# Phosphorylated trimeric SOSS1 complex and RNA polymerase II trigger liquid-liquid phase separation at double-strand breaks

**DOI:** 10.1101/2023.05.10.540130

**Authors:** Qilin Long, Marek Sebesta, Katerina Sedova, Vojtech Haluza, Adele Alagia, Zhichao Liu, Richard Stefl, Monika Gullerova

## Abstract

The most toxic forms of DNA damage are double-strand breaks (DSBs). We have previously shown that RNA polymerase II (RNAPII), phosphorylated at tyrosine 1 (Y1P) on the C-terminal domain, transcribes RNA at DSBs to promote efficient DNA repair. However, it is still unknown how transcription is regulated at DSBs. Here, we show that the trimeric SOSS1 complex (hSSB1, INTS3, and c9orf80) binds to Y1P RNAPII in response to DNA damage, hSSB1 binds to R-loops, and formation of the SOSS1 complex is required for the coexistence of replication protein A (RPA) and hSSB1 at DSBs. The damage-activated tyrosine kinase c-Abl phosphorylates hSSB1 to enable its binding to Y1P RNAPII and its recruitment to DSBs. Finally, we show both *in vitro* and *in vivo* that the SOSS1 complex and RNAPII form dynamic repair compartments at DSBs via liquid-liquid phase separation (LLPS). The loss of the trimeric SOSS1 leads to impaired DNA repair, highlighting its biological importance in the RNA-dependent DNA damage response.

**Teaser:** Trimeric SOSS1 complex and transcription contribute to phase separation at double-strand DNA breaks.

## INTRODUCTION

The stability of the human genome is constantly challenged by numerous endogenous and exogenous insults(*1*). The DNA damage response (DDR) safeguards genome integrity through lesion detection, cell signalling, and DNA repair. The common types of DNA lesions include base conversion(*2*), bulky DNA addition(*3*), single-strand breaks(*4*), and double-strand breaks (DSB)(*5*). DSBs are the most lethal threats to the preservation of genomic information. The persistent, unrepaired DSBs are a major cause of chromosomal aberrations, resulting in genomic instability, cell malfunction, or tumorigenesis(*6*).

Two major mechanisms participating in the repair of DSBs(*5*) are the accurate repair mediated by homologous recombination (HR) and the error-prone repair by non-homologous end-joining (NHEJ). The HR repair pathway is initiated by resection of one of the DNA strands. To protect the exposed ssDNA overhang, single-strand DNA binding (SSB) proteins prevent their degradation. To date, four SSB protein have been characterised: Replication Protein A (RPA), human SSB1 (hSSB1), human SSB2 (hSSB2), and mitochondrial SSB (mtSSB)(*7, 8*). They share evolutionary conserved oligonucleotide binding (OB) domain(s), wherein the heterotrimeric RPA complex belongs to the higher-order SSBs subgroup (with multiple OB-folds) whilst the rest are single-OB fold SSBs(*7*). RPA is well-characterised and plays an essential role in almost all DNA-metabolism pathways(*9*). In contrast, our understanding of the role of the remaining SSBs is limited. hSSB1 was suggested to play a role in HR repair and cell cycle regulation upon activation by ATM(*10*). hSSB1 can form a heterotrimeric sensor of ssDNA (SOSS) complex along with INTS3 and c9orf80, called SOSS1 complex (*11, 12*). Besides its role in DSB repair, hSSB1 also preserves genomic integrity upon hydroxyurea (HU)-mediated(*13*) and oxidative stress(*14*).

Kinases are the key activators of DNA repair pathways. Specifically, three serine/threonine phosphatidylinositol 3-kinase-related kinase (PIKK) family members, Ataxia Telangiectasia mutated (ATM), Ataxia Telangiectasia and Rad3-related (ATR), and DNA-dependent protein kinase (DNA-PK) are critical upstream signal transducers(*15, 16*). They are recruited and activated by corresponding protein complexes, such as MRN (MRE11/RAD50/NBS1), RPA/ATPIP, and KU70/KU86, respectively(*17*). Upon activation, hundreds of proteins are directly phosphorylated in an ATM/ATR- dependent manner, particularly at Ser/Thr-Gln positions(*18, 19*), leading to activation of the checkpoint transducers Chk1 and Chk2(*20, 21*). DNA-PK promotes NHEJ by preventing the end resection and phosphorylating NHEJ machinery factors(*22*). Besides PIKK members, the ubiquitously expressed Abelson kinase c-Abl tyrosine kinase (c-Abl) displays multifaceted roles in regulating phosphorylation status of the DDR factors (*23*). Furthermore, at DSBs, c-Abl phosphorylates RNA polymerase II C-terminal domain (RNAPII CTD) at Tyr1 (Y1) position, which produces damage-responsive transcripts *de novo*, required for efficient repair of DSBs (*24*). R-loops are nucleic acid structures consisting of DNA:RNA hybrid and non-template ssDNA, usually occurring near RNAPII pausing sites. The DSB-associated transcription generates R-loops, which serve as a binding platform for DDR factors to facilitate DNA repair(*24, 25*).

Liquid-liquid phase separation (LLPS) is an important mechanism required for the formation of membrane-less compartments, such as nucleoli, nuclear speckles, RNA granules(*26, 27*), but also gene promoters and super-enhancers(*28*). During LLPS, a part of protein solution condenses into a dense phase, droplets, and the remaining solution forms a dilute phase(*29*). The driving force of LLPS is the weak multivalent interaction of intrinsically disordered regions(*30*). Accumulating evidence shows that LLPS promotes DNA damage repair(*31–33*). Here, we combined *in vivo* and *in vitro* approaches to investigate the role of the SOSS1 complex in regulating the transcription at DSBs. We show that DNA damage-activated tyrosine kinase c-Abl phosphorylates human single-stranded DNA binding (hSSB1) protein. p-hSSB1 binds to INTS3 and c9orf80 leading to the formation of the trimeric SOSS1 complex. The formation of this complex is required for efficient binding of the hSSB1 to R-loops, creating an equilibrium between RPA and hSSB1 binding to ssDNA substrates at DSBs. Furthermore, SOSS1 promotes liquid-liquid phase separation and then serves as a scaffold for Y1P RNAPII, in order to promote transcription at DSBs. The importance of our findings is further supported by the impaired DNA repair observed in cells lacking hSSB1. Thus, this study demonstrates crucial role of the trimeric SOSS1 complex in formation of transient repair compartments and regulation of RNA-dependent DDR.

## RESULTS

### Trimeric SOSS1 complex binds to CTD of RNAPII upon DNA damage

Previously, we identified that RNAPII modified on the C-terminal domain of the catalytic subunit on tyr1 (Y1P RNAPII) actively transcribes RNA at DSBs(*24*). Under non-damage conditions, Y1P RNAPII is mostly detected at the beginning of genes but transcribing in the antisense orientation(*34*). Furthermore, mass spectroscopy (MS) analysis showed that the trimeric SOSS1 complex, but not the SOSS2 complex (hSSB2, INTS3 and c9orf80), interacts with Y1P RNAPII and this interaction is lost when phosphorylation of tyr1 is impaired(*35*).

To investigate the interaction between the trimeric SOSS1 complex and RNAPII, we used proximity ligation assay (PLA), which allows the visualisation of two proteins in their vicinity, and detected the interaction between hSSB1 and RNAPII modified on ser2 (S2P) or ser5 (S5P) RNAPII, respectively. These interactions were sensitive to RNAPII inhibitors, DRB (inhibits CTD phosphorylation by CDK9) and THZ1 (inhibits CTD phosphorylation by CDK7), confirming the specificity of the interaction (fig. S1A and B).

Next, we investigated the interaction between the SOSS1 complex and RNAPII upon DNA damage. First, we used PLA and showed that INTS3 and hSSB1 were in the proximity of DSBs upon ionising irradiation (IR) (fig. S1C). We also detected RNAPII modified on S2P, Y1P, and S5P of the CTD at DSBs (**Fig. 1A**). Furthermore, upon IR, we observed a significant increase in the PLA foci when using hSSB1 or INTS3 and Y1P RNAPII antibodies (**Fig. 1B and C**, single antibodies were used as negative controls), suggesting an increased interaction of the SOSS1 complex with Y1P RNAPII. To support our PLA data, we next performed co-immunoprecipitation experiments, by pulling down hSSB1-GFP in cells stably expressing hSSB1-GFP with or without IR treatment and immunoblotted for Y1P RNAPII. We observed increased levels of Y1P RNAPII in hSSB1-GFP pull downs after IR, which are consistent with the PLA experiments (**Fig. 1D**).

**Figure 1:**
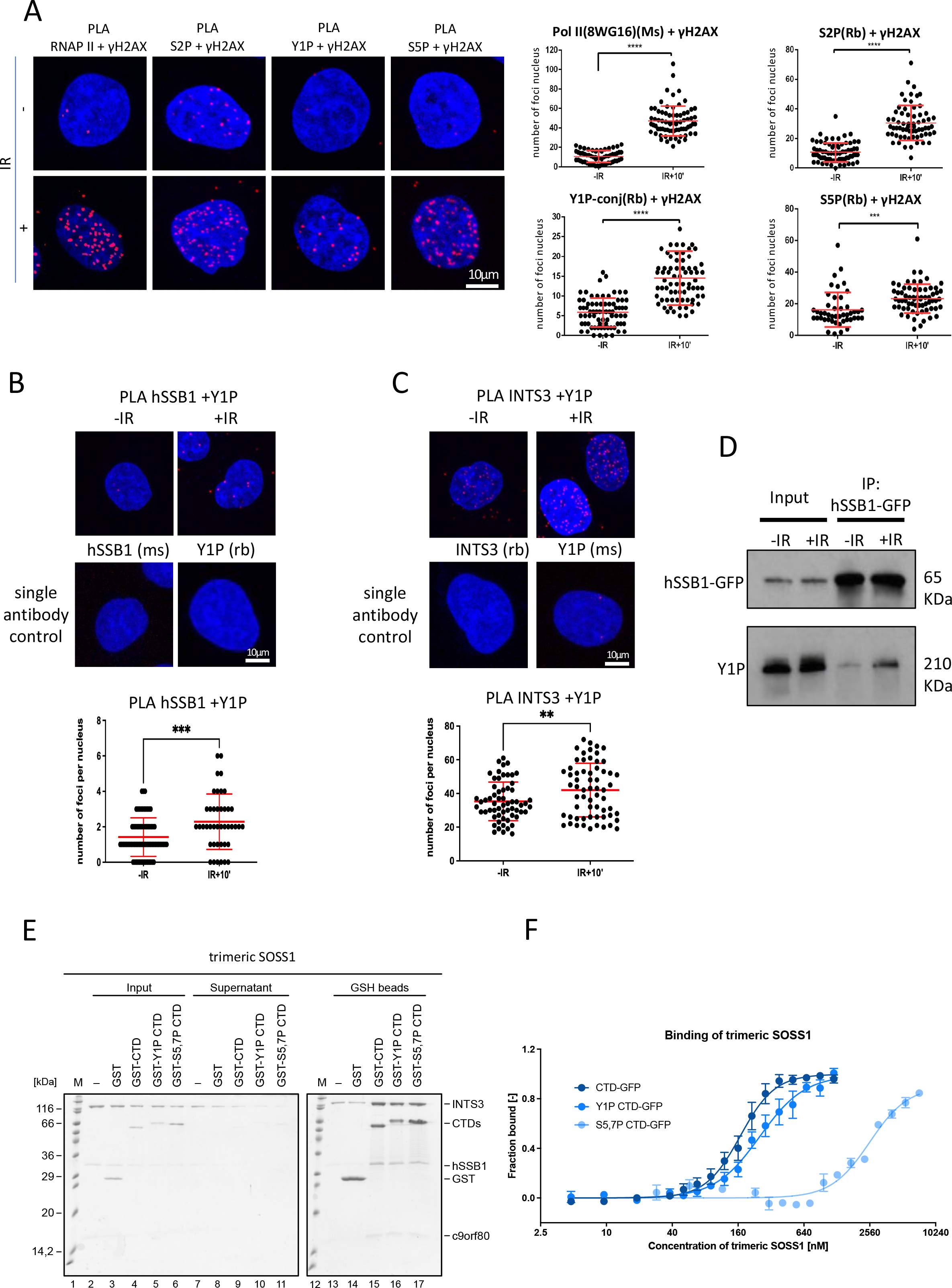
The trimeric SOSS1 complex binds to Y1P CTD of RNAPII upon DNA damage **A)** PLA of RNAPII or S2P or S5P or Y1P and γH2AX in cells with or without IR. IR=10Gy. Left: representative confocal microscopy images; Right: quantification of left, error bar = mean ± SD, significance was determined using non-parametric Mann-Whitney test. \*\*\**p* ≤ 0.001, \*\*\*\**p* ≤ 0.0001. **B)** PLA of hSSB1 and Y1P in cells with or without IR. IR=10Gy. Top: representative confocal microscopy images; bottom: quantification of left, error bar = mean ± SD, significance was determined using non-parametric Mann-Whitney test. \*\*\**p* ≤ 0.001. Single antibody was used as negative control. **C)** PLA of INTS3 and Y1P in cells with or without IR. IR=10Gy. Top: representative confocal microscopy images; bottom: quantification of left, error bar = mean ± SD, significance was determined using non-parametric Mann-Whitney test. \*\**p* ≤ 0.01. Single antibody was used as negative control. **D)** Co-immunoprecipitation of hSSB1-GFP followed by immunoblotting using GFP and Y1P RNAPII antibodies. **E)** *In vitro* pull-down assay of the trimeric SOSS1 complex with GST-CTD, GST-Y1P-CTD or GST-S5,7P-CTD. **F)** Microscale-thermophoresis (MST) binding curves of the trimeric SOSS1 complex with unmodified CTD-GFP, Y1P-CTD-GFP, or S5,7P-CTD-GFP. Measured in triplicates and the lines represent the Hill fit.

To validate the interaction between the SOSS1 complex and RNAPII *in vitro*, we performed pull-down experiments with purified SOSS1 complex and unphosphorylated GST-CTD, Y1P CTD, or S5,7P CTD. All three tested variants of GST-CTD efficiently and specifically pulled-down the trimeric SOSS1 complex (**Fig. 1E**). To determine which subunit of trimeric SOSS1 complex responsible for binding to the CTD of RNAPII, we repeated the pull-down experiments with individual subunits of SOSS1 complex. However, none of the tested proteins (hSSB1, INTS3) interacted with CTD polypeptides (fig. S2A and B). Intriguingly, the trimeric SOSS1 complex assembled from individual, purified subunits failed to interact with CTD as well (fig. S2C), suggesting that the complex formation is required for binding of the trimeric SOSS1 complex to the CTD of RNAPII. To gain quantitative insight into the binding of the trimeric SOSS1 complex to CTD polypeptides, we performed microscale-thermophoresis (MST) analysis, which revealed that the trimeric SOSS1 recognises unphosphorylated CTD and Y1P CTD with similar affinities. Furthermore, the trimeric SOSS1 complex bound with higher affinity to unphosphorylated and Y1P CTD, respectively, than to S5,7P CTD (**Fig. 1F**). Collectively, our *in vivo* and *in vitro* data show that the trimeric SOSS1 complex is associated with RNAPII. The SOSS1 complex preferentially binds Y1 phosphorylated RNAPII and this interaction is enhanced by DNA damage, suggesting that the trimeric SOSS1 may play an important role in DDR.

### hSSB1 binds to DSBs in an R-loop dependent manner

R-loops are transcription-dependent structures found near DSBs. To test whether hSSB1 directly binds to R-loops, we first performed modified PLA(*36*) using antibodies against hSSB1 and S9.6 (recognising R-loop structures) and observed a significant increase in PLA foci upon IR (**Fig. 2A**). We hypothesised that R-loops might act as a binding platform for the hSSB1 recruitment to DSBs. To test this hypothesis, we transiently transfected stably integrated hSSB1-GFP cells with plasmids expressing ribonuclease RNAseH1, an enzyme specifically degrading RNA:DNA hybrids, before subjecting them to laser stripping. First, we confirmed a rapid (from 12sec) recruitment of hSSB1 to DSBs in mock cells. The hSSB1-GFP signal was significantly reduced in presence of RNAseH1 (**Fig. 2B**). To test whether the catalytic activity of RNAseH1 itself caused the reduced hSSB1 occupancy at DSBs, we transfected cells with RNAseH1-GFP WT, WKKD (R-loop binding and catalytic mutant) and D210N (catalytic mutant) plasmids (fig. S3A) and performed PLA. We confirmed a significant increase in RNAseH1-GFP and γH2AX PLA foci upon IR in cells transfected with RNAseH1- GFP WT and D210N mutant, but not with WKKD mutant (fig. S3B). Next, we used the PLA to investigate whether the occupancy of the SOSS1 complex at DSBs is RNAseH1 sensitive. We show that the presence of RNAseH1-GFP WT, but not the mutants, significantly reduced hSSB1 levels at DSBs (**Fig. 2C**).

**Figure 2.**
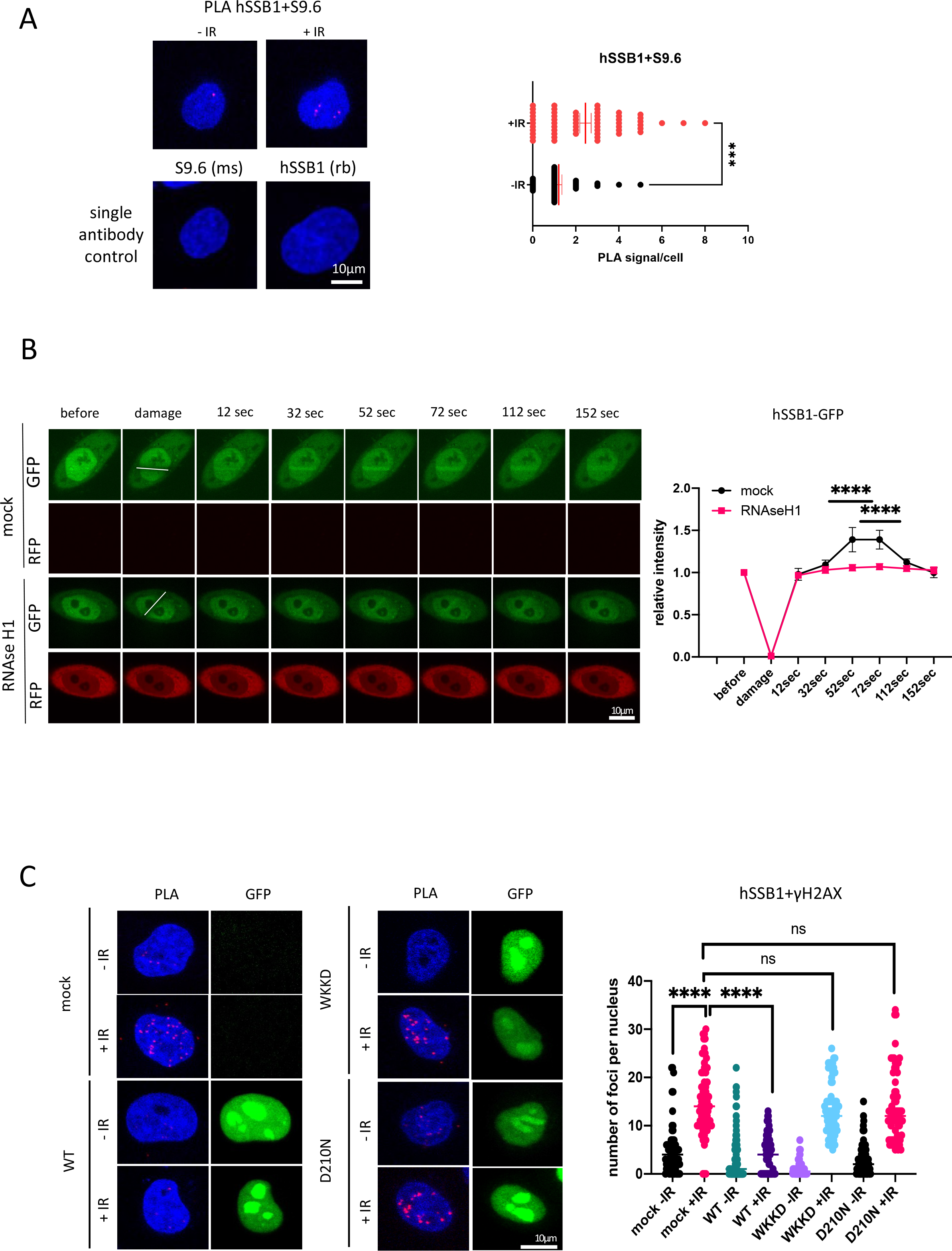
The trimeric SOSS1 complex binds to R-loops upon DNA damage **A)** PLA of hSSB1 and S9.6 (R-loops) with or without IR. IR=10Gy. Left: representative confocal microscopy images; right: quantification of left, error bar = mean ± SD, significance was determined using non-parametric Mann-Whitney test. \*\*\**p* ≤ 0.001. Single antibody was used as negative control. **B)** Laser stripping of stably integrated hSSB1-GFP cells with or without transiently expression of RNAseH1-RFP plasmid. Representative spinning disk confocal microscopy images and quantification (n≥10) showing GFP and RFP signals before and after laser striping at indicated time points; error bar = mean ± SEM; significance was determined using multiple unpaired Student’s *t*-test. \*\*\*\**p* ≤ 0.0001. **C)** PLA of hSSB1 and γH2AX in cells transiently transfected with RNAseH1^wt^-GFP or RNAseH1^WKKD^-GFP (binding and catalytic) or RNAseH1^D210N^-GFP (catalytic) mutants with or without IR. IR=10Gy. Left: representative confocal microscopy images; right: quantification of left, error bar = mean ± SD, significance was determined using non-parametric Mann- Whitney test. \*\*\*\**p* ≤ 0.0001.

These data suggest that the recruitment of hSSB1 to DSBs is transcription and R-loop dependent.

### The formation of the SOSS1 complex is required to suppress the inhibitory effect of RPA on hSSB1 binding to R-loops

hSSB1, together with RPA, belongs to the single-stranded DNA binding (SSB) protein family, which is characterised by the presence of at least one structurally conserved OB-fold domain(*7*). Given the extremely high affinity of RPA to ssDNA, it is unclear how hSSB1 and RPA may coexist at resected DSBs. Interestingly, ssDNA is not only present at the DNA ends but is also part of transcription-associated R-loops (*37, 38*). To understand how RPA and hSSB1 (and the trimeric SOSS1) may coexist at DSBs, we performed a comprehensive DNA binding analysis of trimeric SOSS1 and its subunits on a broad range of nucleic acid (NA) substrates (21nt ssDNA, 61nt ssDNA, 61nt ssRNA, 61nt dsDNA, DNA/RNA hybrid, R-loop, DNA bubble) *in vitro*. By using electrophoretic mobility shift assay (EMSA), we firstly determined the binding preferences of hSSB1 using the abovementioned nucleic acid substrates. As expected from literature(*10*), hSSB1 bound ssDNA in a length-dependent manner: hSSB1 did not bind to 21nt ssDNA nor 61nt dsDNA, whilst exhibiting high affinity towards 61nt ssDNA (**Fig. 3A**, fig. S4A-C). Surprisingly, hSSB1 bound R-loop structures and RNA:DNA hybrids with similar affinity to 61nt ssDNA, but not bubble DNA (R-loop without RNA) (**Fig. 3A**, fig. S4 D-F). As the ssDNA portion in these complex structures is 21nt long, it suggests that hSSB1 binds to R-loops specifically. Finally, hSSB1 binds to ssRNA with similar affinity as to ssDNA (fig. S4G, H). The trimeric SOSS1 exhibited comparable affinities to substrates as hSSB1 (**Fig. 3B**, fig. S5A-H). However, INTS3 did not bind to any of the structures (fig. S6A-D), therefore we conclude that hSSB1 is the subunit of the trimeric SOSS1 responsible for binding to nucleic acids (NA).

**Figure 3:**
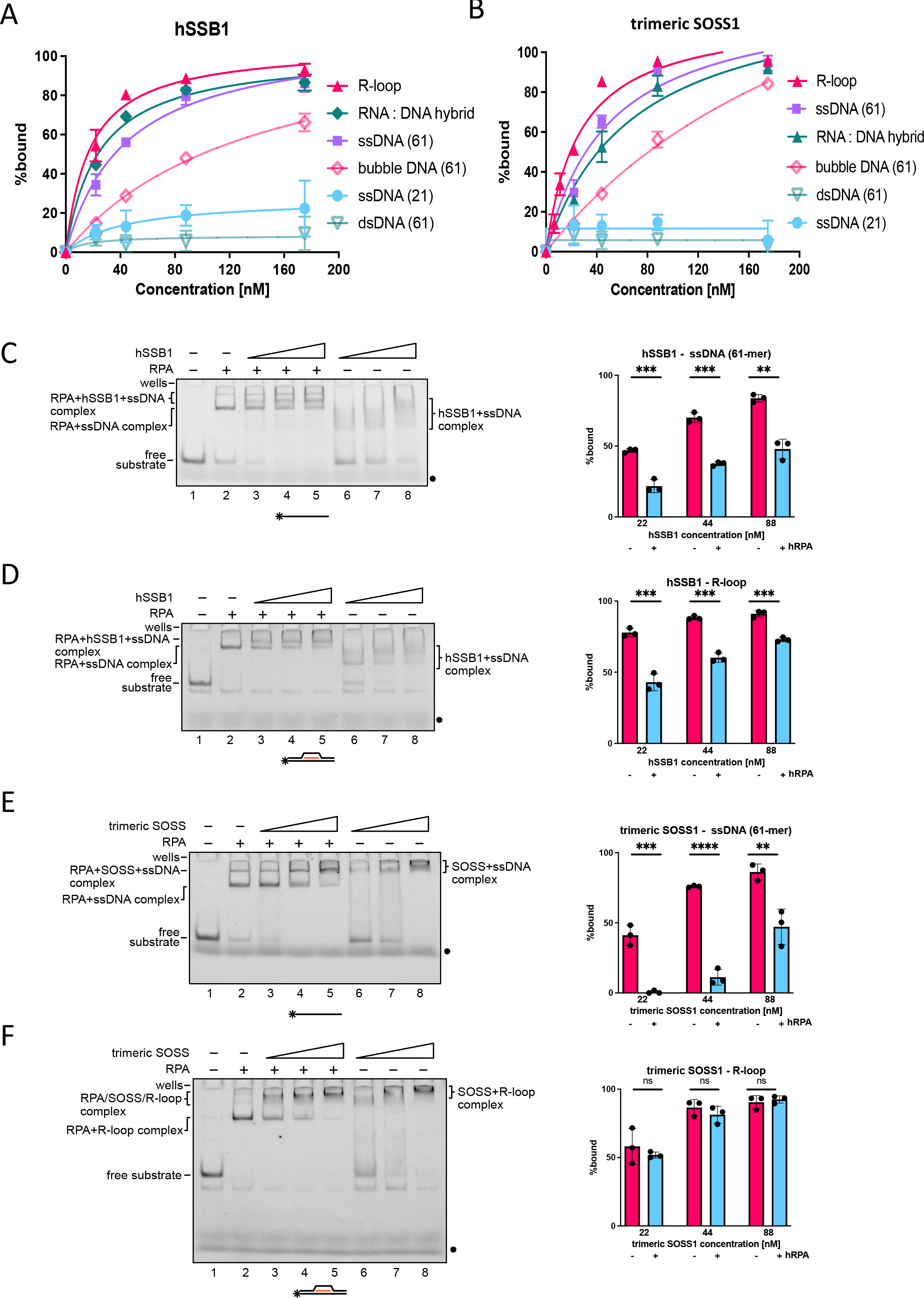
Trimeric SOSS1 complex suppresses the inhibitory effect of RPA in hSSB1- mediated binding of R-loops **A)** Graph representing quantification of EMSA experiments (n=3) conducted between hSSB1 and R-loop, RNA:DNA hybrid, 61-mer ssDNA, bubble DNA, 21-mer DNA, and 61-mer dsDNA respectively. **B)** Graph representing quantification of EMSA experiments (n=3) conducted between trimeric SOSS1 and R-loop, RNA:DNA hybrid, 61-mer ssDNA, bubble DNA, 21-mer DNA, and 61- mer dsDNA respectively. **C)** Scan of representative EMSA experiments (left) and bar chart graph (right) representing quantification of EMSA experiments (n=3) conducted between hSSB1 (at indicated concentrations) and ssDNA (61-mer) in the absence or presence of RPA. **D)** Scan of representative EMSA experiments (left) and bar chart graph (right) representing quantification of EMSA experiments (n=3) conducted between hSSB1 (at indicated concentrations) and R-loop in the absence or presence of RPA. **E)** Scan of representative EMSA experiments (left) and bar chart graph (right) representing quantification of EMSA experiments (n=3) conducted between the trimeric SOSS1 (at indicated concentrations) and ssDNA (61-mer) in the absence or presence of RPA. **F)** Scan of representative EMSA experiments (left) and bar chart graph (right) representing quantification of EMSA experiments (n=3) conducted between the trimeric SOSS1 (at indicated concentrations) and R-loop in the absence or presence of RPA.

Lastly, we tested the binding of RPA to the set of NA substrates. RPA showed high affinity to short (21nt) and long (61nt) ssDNA and ssRNA substrates but no binding to 61nt dsDNA (fig. S7A-C,G,H). We also detected RPA binding to R-loops and RNA:DNA hybrids with high affinity, whilst binding to DNA bubble was less efficient (fig. S7D-F and H).

To investigate the NA binding ability of the trimeric SOSS1 complex and its subunits in the presence of RPA, we performed a set of competitive EMSA. We first pre-incubated the NA substrates with RPA at either 10 or 30 nM and then included the tested individual subunits or SOSS1 complex. RPA significantly inhibited the binding of hSSB1 to ssDNA (61nt), RNA:DNA hybrids, and R-loop substrates, respectively (**Fig. 3C and D**, fig. S8A-D). The outcome contrasted when the trimeric SOSS1 was tested. RPA did significantly reduce the binding of the trimeric SOSS1 complex to ssDNA (**Fig. 3E**, fig. S9A), whilst it did not prevent the binding of the trimeric SOSS1 to R-loop structure (**Fig. 3F**, fig. S9B). Additionally, RPA did not have any effect on SOSS1 binding to RNA:DNA hybrids nor to short ssDNA (21nt) (fig. S9C and D).

Taken together, these data show that hSSB1 alone or as a part of trimeric SOSS1 complex binds specifically and with high affinity to R-loop and RNA:DNA hybrids, and the formation of the trimeric SOSS1 complex is required to overcome the inhibitory binding of RPA to the same NA substrates. We conclude that RPA preferentially binds to resected ssDNA at the site of DSB, whilst hSSB1 and the trimeric SOSS1 complex associate with R-loops.

### c-ABL phosphorylates hSSB1 upon DNA damage

We have previously shown that DNA damage activated c-Abl phosphorylates Y1P CTD RNAPII at DSBs, which leads to the production of strand-specific, damage-responsive transcripts (DARTs)(*24*). To investigate whether c-Abl phosphorylates hSSB1 or other components of the SOSS1 complex, we first performed PLA with antibodies against c-Abl and hSSB1 and detected PLA foci corresponding to c-Abl and hSSB1 interaction upon IR treatment. The number of these foci was significantly reduced in the presence of c-Abl inhibitor Imatinib (**Fig. 4A**, single antibodies were used as a negative control). To extend these data, we repeated PLA using antibody against phosphorylated c-Abl (p-c-Abl) and detected an Imatinib-sensitive interaction between p-c-Abl and hSSB1, suggesting that IR-activated c-Abl interacts with hSSB1 in response to DNA damage (**Fig. 4A**). Next, we performed co- immunoprecitipation assay by pulling down hSSB1-GFP from cells subjected to IR and Imatinib treatment. Immunoblotting with pan-phospho-tyrosine antibody revealed specific Imatinib sensitive damage induced hSSB1 phosphorylation (**Fig. 4B**).

**Figure 4:**
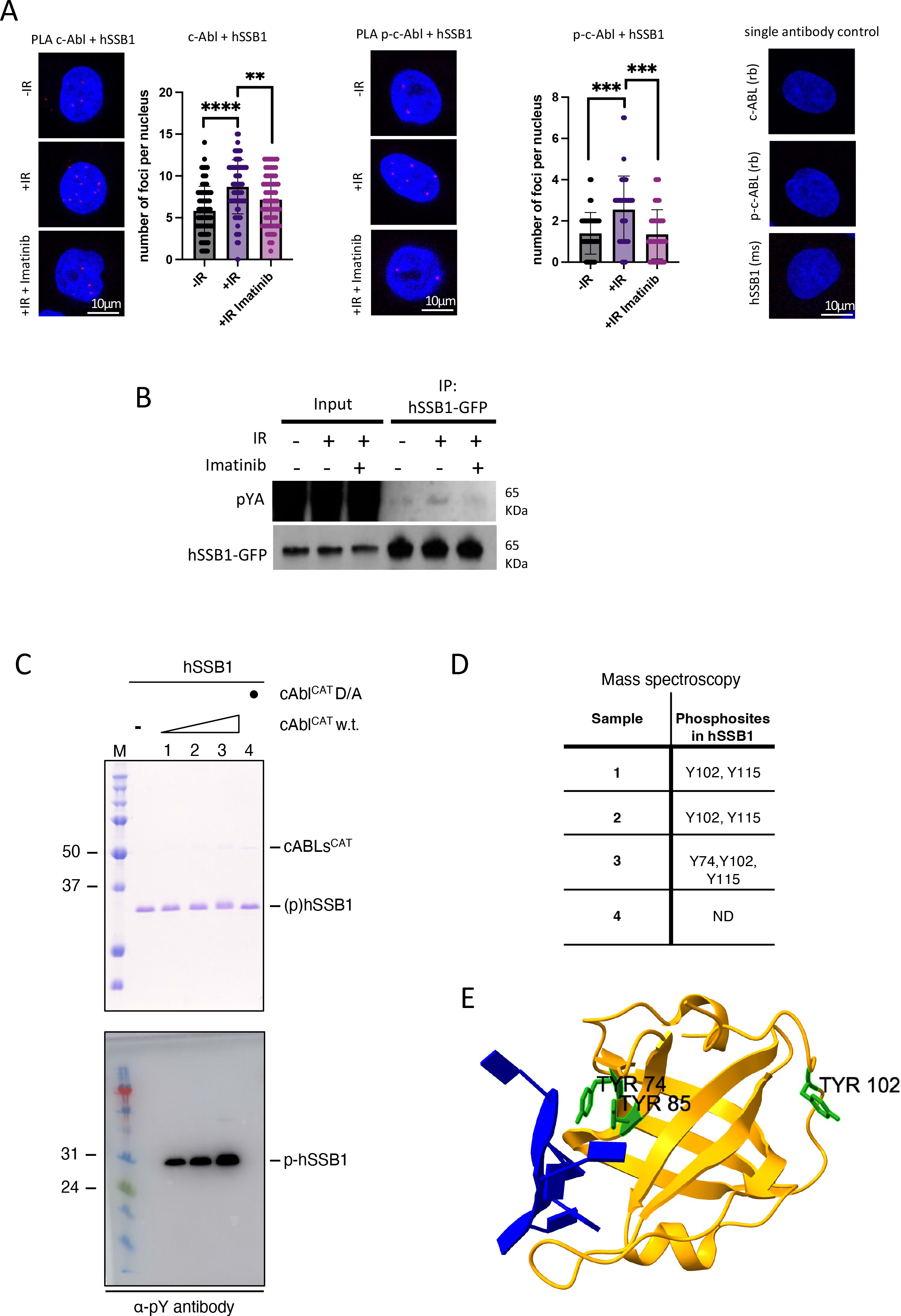
cAbl phosphorylates hSSB1 upon DNA damage **A)** PLA of cAbl /p-cAbl and hSSB1 without IR, with IR, and with IR and Imatinib. IR=10Gy. Cells were treated with 1μM Imatinib for 1h before IR. Left: representative confocal microscopy images; right: quantification of left, error bar = mean ±SD, significance was determined using non-parametric Mann-Whitney test. \**p* ≤ 0.05, \*\**p* ≤ 0.01, \*\*\**p* ≤ 0.001. Single antibody was used as negative control. **B)** Immunoprecipitation of hSSB1-GFP from cells subjected to IR treatment and Imatinib. Immunoblot showing signals for pan-phopsho-tyrosine (α-pY) and hSSB1-GFP. **C)** *In vitro* phosphorylation of hSSB1 by cAbl^CAT^. The upper panel depicts an SDS-PAGE gel of the reaction. The lower panel depicts immunodetection of the reaction by western blotting with an α-pY antibody. **D)** Identification of the cAbl^CAT^ mediated phosphorylation sites on hSSB1 by mass spectrometry. A table with identified residues. The sample number in the table corresponds with the numbering in (C). **E)** Depiction of the position of tyrosine residues (in green) of hSSB1 (yellow) on the structural model with ssDNA (blue) (PDB ID: 4OWW). Residue Y115 is not visible/ highlighted in the structure due to its high flexibility.

In order to validate our *in vivo* data *in vitro*, incubated purified hSSB1 (fig. S10A and B) with purified catalytic domain of c-Abl wt or its kinase dead variant (c-Abl^D363A^). Interestingly, we observed a clear shift of hSSB1 protein bands on PAGE gel, which correlated with an increasing concentration of c-Abl wt. This was not the case when hSSB1 was incubated with c-Abl^D363A^ (**Fig. 4C, top panel**). To confirm that the shift in the protein band on the gel was indeed caused by phosphorylation deposited by c-Abl, we performed immunoblotting with pan-phosphorylated tyrosine antibody (pYA) and detected positive signals, which corresponded to at the size hSSB1 protein (**Fig. 4C, bottom panel**). To test that c-Abl activity on hSSB1 is specific, we incubated c-Abl wt or c-Abl^D363A^ with non-specific substrate GST and as expected, did not detect any GST phosphorylation (fig. S10C and D). Next, we wished to determine the exact tyrosine residues on hSSB1 that are phosphorylated by c-Abl. We cut out the corresponding bands from the PAGE gel (**Fig. 4C)** and subjected them to massspectroscopy (MS) (Supplementary file 1). MS analysis identified phosphorylation on hSSB1 residues Y102, Y115, and Y74, among which the Y102 and Y115 were presented in every sample (**Fig. 4D**). Next, we visualised phosphorylated tyrosine residues on hSSB1 protein structure. Interestingly, Y74 and previously identified Y85 (*39*) residues are located on the ssDNA binding interface, suggesting they might be involved in hSSB1 binding to NA substrates. The Y102 residue is present on hSSB1-INTS3 interphase, possibly playing a role in SOSS1 complex assembly. The Y115 residue is on hSSB1 unstructured domain, hence not visible in the model (**Fig. 4E** and fig. S10E).

Taken together, our *in vivo* and *in vitro* experiments show that damage activated c-Abl phosphorylates hSSB1 at tyrosine residues upon IR.

### hSSB1 phosphorylation on Y102 and Y115 residues is required for its localization at DSBs and interaction with Y1P RNAPII

To test whether c-Abl mediated phosphorylation of hSSB1 plays a role in DDR, we performed laser stripping of stably integrated hSSB1-GFP cells *in vivo*. First, we detected hSSB1-GFP to be rapidly (32 seconds after laser damage) recruited to the sites of DNA damage. This recruitment was significantly abolished in the presence of Imatinib (fig. S11A). To confirm these data, we employed PLA using antibodies against hSSB1 or c-Abl and γH2AX and observed PLA foci upon IR, which were significantly reduced by Imatinib treatment (fig. S11B and C, single antibodies were used as a negative control).

To further investigate the requirement of c-Abl mediated phosphorylation of hSSB1 into more details, we generated Y102, Y115, and double Y102,Y115 hSSB1 non-phosphorylatable mutants (fig. S12A). We transiently transfected cells with plasmids expressing hSSB1-GFP wt, Y102A, Y115A, and Y102A,Y115A respectively and performed PLA using GFP (detecting hSSB1-GFP expressed from the plasmids) and γH2AX antibodies. We observed positive foci in hSSB1 wt cells after IR as before, but the number of foci was significantly reduced in all three mutants (**Fig. 5A**), suggesting that the phosphorylation on hSSB1 on Y102 and Y115 is important for its localisation to DSBs. Additionally, we performed PLA assay using hSSB1 and γH2AX antibodies, as expected, the positive foci were detected only in cells transfected with hSSB1 wt plasmids but not with mutant plasmids (fig. S12B). Next, we generated stable cell lines with integrated hSSB1-GFP wt, Y102A, Y115A, and Y102,Y115A mutants and subjected them to laser stripping. Laser induced DNA damage led to rapid recruitment of hSSB1-GFP wt, but not the mutants to DSBs (**Fig. 5B**). Similarly, laser stripping of transiently transfected cells with hSSB1-GFP wt or its mutants resulted in the recruitment of hSSB1-GFP wt to the proximity of DSBs, whilst the mutants did not (fig. S12C).

**Figure 5:**
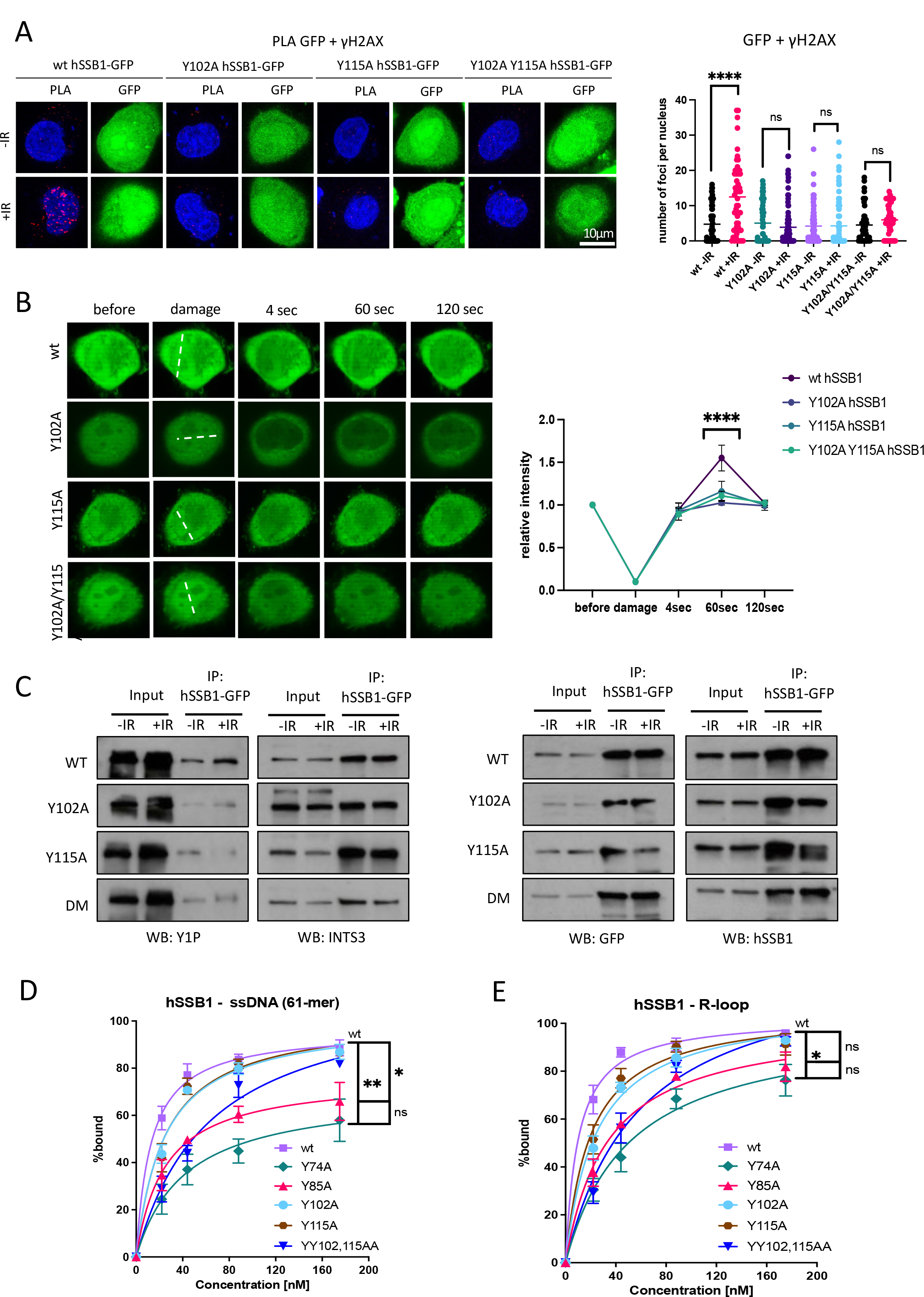
Phosphorylation of hSSB1 is required for its localization at DNA double- strand breaks, interaction with RNAPII Y1P and binding to R-loops. **A)** PLA of GFP and γH2AX in cells transiently transfected with hSSB1^wt^-GFP or hSSB1^Y102A^- GFP, hSSB1^Y115A^-GFP and hSSB1^Y102A/Y115A^-GFP plasmids treated with or without IR. IR=10Gy. Left: representative confocal microscopy images; right: quantification of left, error bar = mean ± SD, significance was determined using non-parametric Mann-Whitney test. \*\*\*\**p* ≤ 0.0001. **B)** Laser stripping of stably integrated hSSB1^wt^-GFP or hSSB1^Y102A^-GFP, hSSB1^Y115A^-GFP and hSSB1^Y102A/Y115A^-GFP cells. Representative spinning disk confocal microscopy images and quantification (n≥10) showing GFP signals before and after laser striping at indicated time points; error bar = mean ± SEM; significance was determined using multiple unpaired Student’s *t*-test. \*\*\*\**p* ≤ 0.0001. **C)** Co-immunoprecipitation of hSSB1-GFP from stably integrated hSSB1^wt^-GFP or hSSB1^Y102A^-GFP, hSSB1^Y115A^-GFP and hSSB1^Y102A/Y115A^-GFP cells with or without IR treatment. IR=10Gy. Immunoblots show signals for Y1P RNAPII, INTS3, GFP and hSSB1. **D)** Graph representing quantification of EMSA experiments (n=3) conducted between hSSB1 wt, Y74A, Y85A, Y102A, Y115A, and YY102,115AA mutants (at indicated concentrations) and ssDNA (61-mer). Significance was determined using unpaired *t*-test. \*\**p* ≤ 0.01 and \**p* ≤ 0.05 represent the comparison between hSSB1 wt and Y74A, Y85A, respectively. **E)** Graph representing quantification of EMSA experiments (n=3) conducted between hSSB1 wt, Y74A, Y85A, Y102A, Y115A, and YY102,115AA mutants (at indicated concentrations) and R-loop. Significance was determined using unpaired *t*-test. \**p* ≤ 0.05 represent the comparison between hSSB1 wt and Y74A.

Next, we asked whether hSSB1 mutants lose their ability to bind to Y1P RNAPII or INTS3. We used co-immunoprecipitation of hSSB1-GFP wt, Y102A, Y115A, and double Y102A,Y115A mutants, followed by immunoblotting using Y1P or INTS3 antibodies. We observed that hSSB1 non-phosphorylatable mutants did not bind Y1P RNAPII as efficiently as hSSB1 wt, whilst their binding to INTS3 was not affected (**Fig. 5C**). Next, we purified the trimeric SOSS1 wt, Y102A, Y115A, and Y102A,Y115A complexes (fig. S13A) and used them for *in vitro* pull-down assay with Y1P RNAPII. Interestingly, all SOSS1 complexes were able to bind to Y1P RNAPII *in vitro* (fig. S13B). We hypothesise that the *in vitro* binding of the mutant SOSS1 variants to Y1P RNAP suggests that the direct, physical interaction is not hampered by the Y to A substitutions, whilst *in vivo* the interaction may take place upon recruitment of the trimeric SOSS1 complex to the sites of DSBs, which is abrogated by the said mutations.

Overall, these data demonstrate that c-Abl mediated phosphorylation of Y102 and Y115 residues is required for hSSB1 recruitment to DSBs and its binding to Y1P RNAPII *in vivo*.

### Phosphorylation on Y74 and Y85 hSSB1 residues is required for its binding to NA substrates

c-Abl mediated phopshorylation of hSSB1 was also detected on Y74 and Y85 residues, which are located at the DNA binding interface of hSSB1 (**Fig. 4E**). The phosphorylation on these two sites on hSSB1 has been implicated in ssDNA binding (*39, 40*). Therefore, we tested whether they are indeed required for hSSB1 binding to NA substrates. We purified hSSB1 wt, Y74A, Y85A, Y102A, Y115A, and Y102A,Y115A and tested them in EMSA with various NA substrates (as in **Fig. 3**). Interestingly, the Y74A and Y85A hSSB1 mutants, but not Y102A, Y115A, nor Y102A,Y115A variants (alone or as part of trimeric SOSS1), bound 61nt ssDNA and R-loops less efficiently than hSSB1 wt (**Fig. 5D and E**, fig. S13D-G, fig. S14A-D).

These data show that c-Abl mediated phosphorylation of hSSB1 on Y74 and Y85 residues is important requirement for hSSB1 binding to NA substrates.

### *In Vitro* phase separation of trimeric SOSS1 is promoted by DNA and CTD of RNAPII

Next, we asked what the role of phosphorylated SOSS1 at DSBs is. As the subunits of the trimeric SOSS1 lack clear enzymatic activity, we conjectured that its role might be indirect. Previous structural work(*40*) suggested that the C-termini of INTS3 and hSSB1 are largely unstructured, intrinsically disordered regions (IDRs). IDRs on other proteins can drive LLPS (*41, 42*). To test, whether SOSS1 is able to phase separate, we purified the trimeric SOSS1 with mCerulean fluorescent protein fused to the C-terminus of INTS3. The trimeric SOSS1 alone, in the presence of ssDNA, nor ssRNA did not phase separate. When crowding agent (5% PEG-8000) was included, we observed robust, concentration-dependent appearance of condensates, which are classic characteristic of LLPS. Furthermore, addition of both ssDNA and ssRNA resulted in a statistically significant increase in the number of condensates, suggesting that ssDNA may promote LLPS of the trimeric SOSS1 (**Fig. 6A and B**, fig. S15A- C).

**Figure 6:**
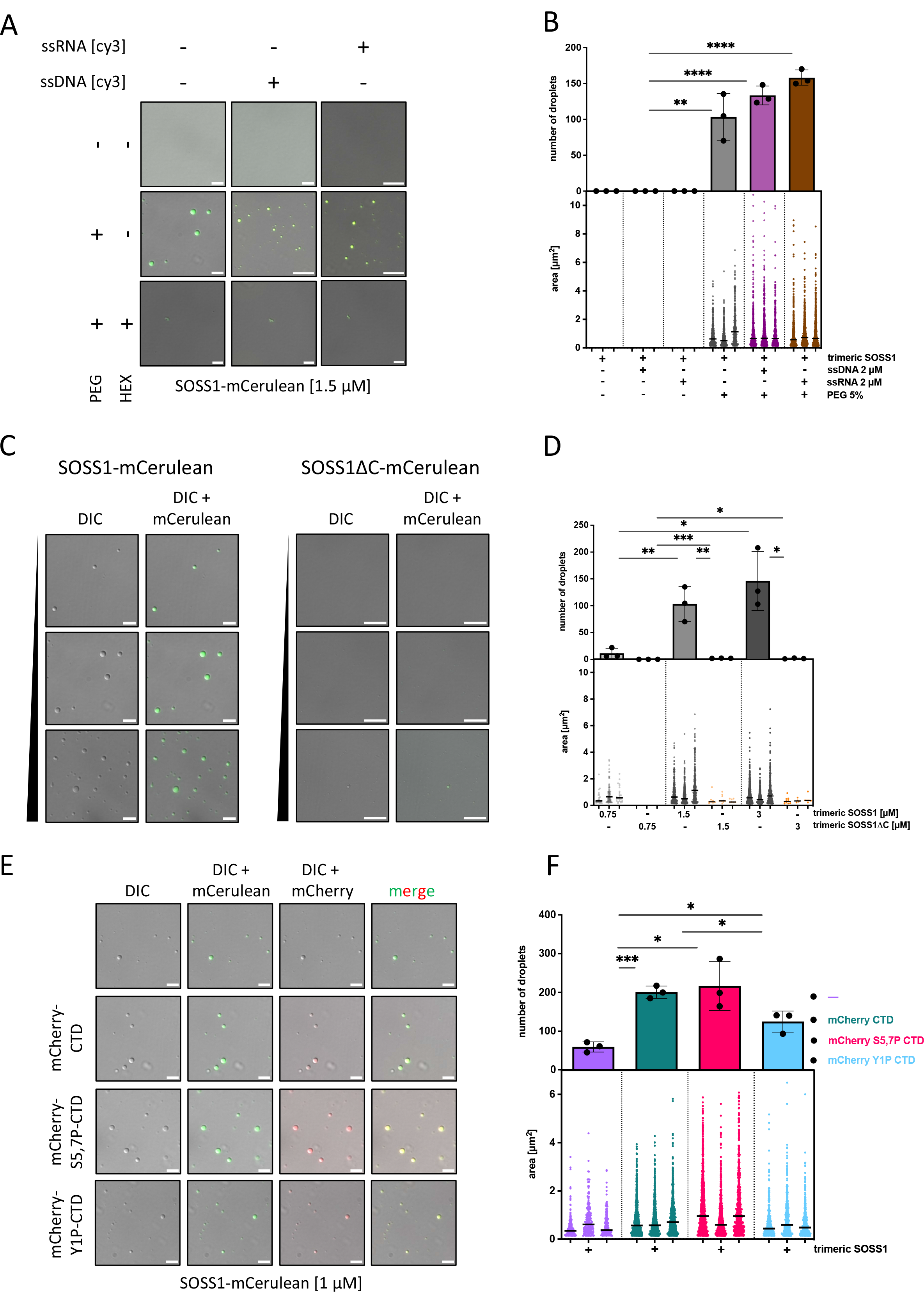
Phase-separation of the trimeric SOSS complex is promoted by nucleic acids and CTD of RNAPII *in vitro* **A)** LLPS experiments of purified trimeric, fluorescently labelled SOSS1 complex (on INTS3 subunit), determining the effect of crowding agent (5% PEG-8000), ssDNA (at 2 µM), and ssRNA (at 2 µM) on the efficiency of phase separation of the complex. Representative images from three experiments are depicted as overlay of differential interference contrast (DIC), mCerulean, and cy3 (where nucleic acids are present) channels. Hexane-1,6-diol (at 10%, HEX) was added to inhibit hydrophobic interactions. Scale bar: 5µM. PEG: PEG-8000. **B)** Bar chart (upper panel) represents quantification (n=3) of number of droplets from LLPS experiments shown in A). Statistical significance was determined by unpaired *t*-test. Nested scatter plot (lower panel) represents quantification (n=3) of area of individual droplets from three independent experiments shown in A), with median area determined per dataset. Statistical significance was determined by nested *t*-test. **C)** Determination of the domain responsible for LLPS of the trimeric SOSS1 complex. Representative images from three experiments with SOSS1 complexes with either wild-type INTS3 or INTS3^1-961^ (INTS3ΔC), labelled with mCerulean at 0.75, 1.5, and 3 µM. The images are depicted as overlay of differential interference contrast (DIC) and mCerulean channels. **D)** Bar chart (upper panel) represents quantification (n=3) of number of droplets from LLPS experiments shown in C). Statistical significance was determined by unpaired *t*-test. Nested scatter plot (lower panel) represents quantification (n=3) of area of individual droplets from three independent experiments shown in C), with median area determined per dataset. Statistical significance was determined by nested *t*-test. **E)** LLPS experiments investigating the effect of mCherry-labelled CTDs: unmodified, phosphorylated on serines 5,7 (S5,7P), and tyrosine 1 (Y1P) at 0.75µM, respectively, on phase separation with the trimeric SOSS1 complex (1µM). Representative images from three experiments are depicted as DIC, overlay of DIC and mCerulean, overlay of DIC and mCherry, and overlay of all three channels. Scale bar: 5µM. **F)** Bar chart (upper panel) represents quantification (n=3) of number of droplets from LLPS experiments shown in C). Statistical significance was determined by unpaired *t*-test. Nested scatter plot (lower panel) represents quantification (n=3) of area of individual droplets from three independent experiments shown in C), with median area determined per dataset. Statistical significance was determined by nested *t*-test.

To identify the region within the trimeric SOSS1 complex responsible for phase-separation, we constructed variant of the trimeric SOSS1 complex, in which the IDR region of INTS3 (AA961-1043) was absent (SOSS1ΔC). When tested across concentrations, the trimeric SOSS1ΔC complex did not phase-separate (**Fig. 6C,D**), suggesting that the domain responsible for phase-separation of the trimeric SOSS1 complex is indeed located within the IDR region of INTS3.

To rule-out the possibility that only INTS3 enters the droplets, we generated a variant trimeric SOSS1 complex in which also the hSSB1 subunit was tagged with a different fluorescent tag (mOrange), along INTS3, tagged with mCerulean. We observed that such complex did indeed phase separate (fig. S15D,F), albeit to a lesser extent compared to the trimeric SOSS1 complex tagged only on INTS3 subunit (compare fig. 15C and 15F). In the observed droplets we did detect signal for both INTS3 and hSSB1, suggesting that the entire complex phase-separate into droplets. Fittingly, when the IDR regions of both INTS3 and hSSB1 (AA106-221) were absent, the resulting trimeric SOSS1 complex did not phase-separate (fig. 15E).

We next wondered whether trimeric SOSS1 may form heterotypic condensates, thereby serving as a scaffold for additional proteins. As it is widely accepted that RNAPII, via its CTD, may be one of such proteins(*43–45*). We therefore tested this hypothesis. Whilst S5,7P-CTD and Y1P-CTD efficiently entered the condensates, the unmodified CTD entered to a lower extent (**Fig. 6E**). Importantly, all three forms of CTD promoted phase separation of trimeric SOSS1, as evidenced by the increase in the numbers of condensates formed (**Fig. 6F**). This effect is not caused by phase separation of CTD itself, since at the tested conditions, none of CTD alone phase separated (fig. S16A).

Next, we investigated whether both ssDNA and CTD may enter the same condensates. To perform this experiment, we used unmodified trimeric SOSS1. Based on our data, both S5,7P CTD and ssDNA alone entered the same condensates of trimeric SOSS1 (fig. S16B and C).

Finally, we tested whether IDR on INTS3 is required for LLPS formation.

Collectively, our results suggest that the trimeric SOSS1 efficiently phase separates *in vitro*, and these condensates simultaneously accommodate RNAPII CTD and DNA, thereby they may be promoting efficient transcription at DSBs.

### hSSB1 and INTS3 phase separate into optoDroplets *in vivo*

In order to test whether the SOSS1 complex is able to phase separate *in vivo*, we used the optoDroplet system(*46*), which employs the photolyase homology region (PHR) of *Arabidopsis thaliana,* Cry2, fusion to protein of interest. PHR self-associates upon blue light induction, hence enabling to visualise and recapitulate phase separation of tested proteins in cells. First, we cloned full length hSSB1 and INTS3 into mCherry-PHR (Cry2) plasmids(*46*). As positive controls, we used Cry2-fused to IDRs of RNA-binding proteins FUS and hnRNPA1(*46*). We stably integrated all constructs into HeLa cells and subjected them to light induction and time-lapse confocal microscopy using red channel (561 nm) acquisition and green channel (488 nm) channel for blue light induction every 10 seconds in 40 frames (**Fig. 7A**). As expected, in cells expressing Cry2 alone, no droplet formation was detected upon light induction (**Fig. 7B** and Supplementary movie 1). Fusion of FUS and hnRNPA1 IDRs with Cry2 resulted in a time-dependent optoDroplet formation, as it was shown previously(*46*) (**Fig. 7C and D** and Supplementary movies 2 and 3). Interestingly, the expression of Cry2-hSSB1 and Cry2-INTS3 in cells resulted in light induced optoDroplet formation (**Fig. 7E and F** and Supplementary movies 4 and 5).

**Figure 7:**
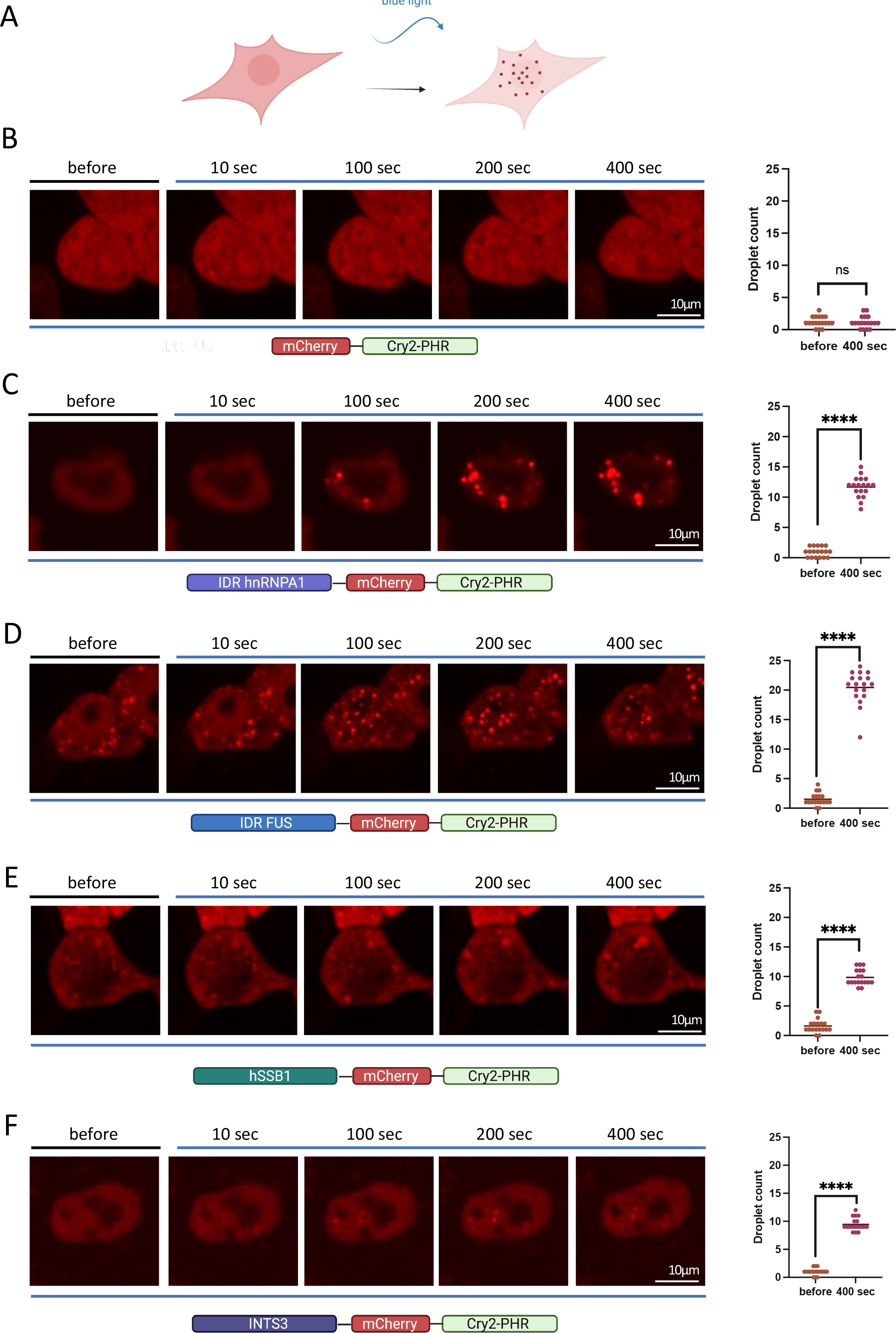
SOSS1 phase separates *in vivo* **A)** Diagram showing the optoDroplet strategy. **B)** Light induced optoDroplet formation of Cry2-mCherry. Cells were imaged at 10 seconds intervals 40 times. Representative images of optoDroplets before and during light induction are shown, specific time points are indicated. Quantification of optoDroplets from 3 independent experiments show values for optoDroplets before and at 400 seconds after light induction. **C)** As in B for IDR-hnRNAPA1-Cry2-mCherry. Significance was determined parametric student t- test. \*\*\*\**p* ≤ 0.0001 **D)** As in B for IDR-FUS-Cry2-mCherry. Significance was determined parametric student t- test. \*\*\*\**p* ≤ 0.0001 **E)** As in B for hSSB1-Cry2-mCherry. Significance was determined parametric student t- test. \*\*\*\**p* ≤ 0.0001 **F)** As in B for INTS3-Cry2-mCherry. Significance was determined parametric student t- test. \*\*\*\**p* ≤ 0.0001

All together, these results show that hSSB1 and INTS3 subunits of the SOSS1 complex are able to phase separate *in vivo*.

### hSSB1 and INTS3 contribute to phase separation at DSBs

A recent study showed that RNAPII CTD is able to phase separate in order to form hubs at actively transcribed genes(*28*). We showed that the SOSS1 complex interacts with RNAPII upon DNA damage (**Fig.1** and fig. S1). It has been proposed that at DSBs phase separation integrate localised DNA damage repair into compartments(*32*). Therefore, we investigated whether the SOSS1 complex might contribute to phase separation at DSBs. To do so, we modified the light inducible optoDroplet protocol and added laser (405 nm) striping prior the light induction (**Fig. 8A**). Neither Cry2, Cry2-FUS-IDR, nor Cry2-hnRNPA1-IDR were recruited to the laser stripes, suggesting that the IDR domain alone is not sufficient for DSB recruitment (fig. S17A-C and Supplementary movies 6, 7 and 8). As expected, both Cry2-hSSB1 and Cry2-INTS3 were rapidly recruited to DSBs. Interestingly, we observed fission and optoDroplet formation in a time-dependent manner within the laser stripe area (**Fig. 8B-D** and Supplementary movies 9 and 10).

**Figure 8:**
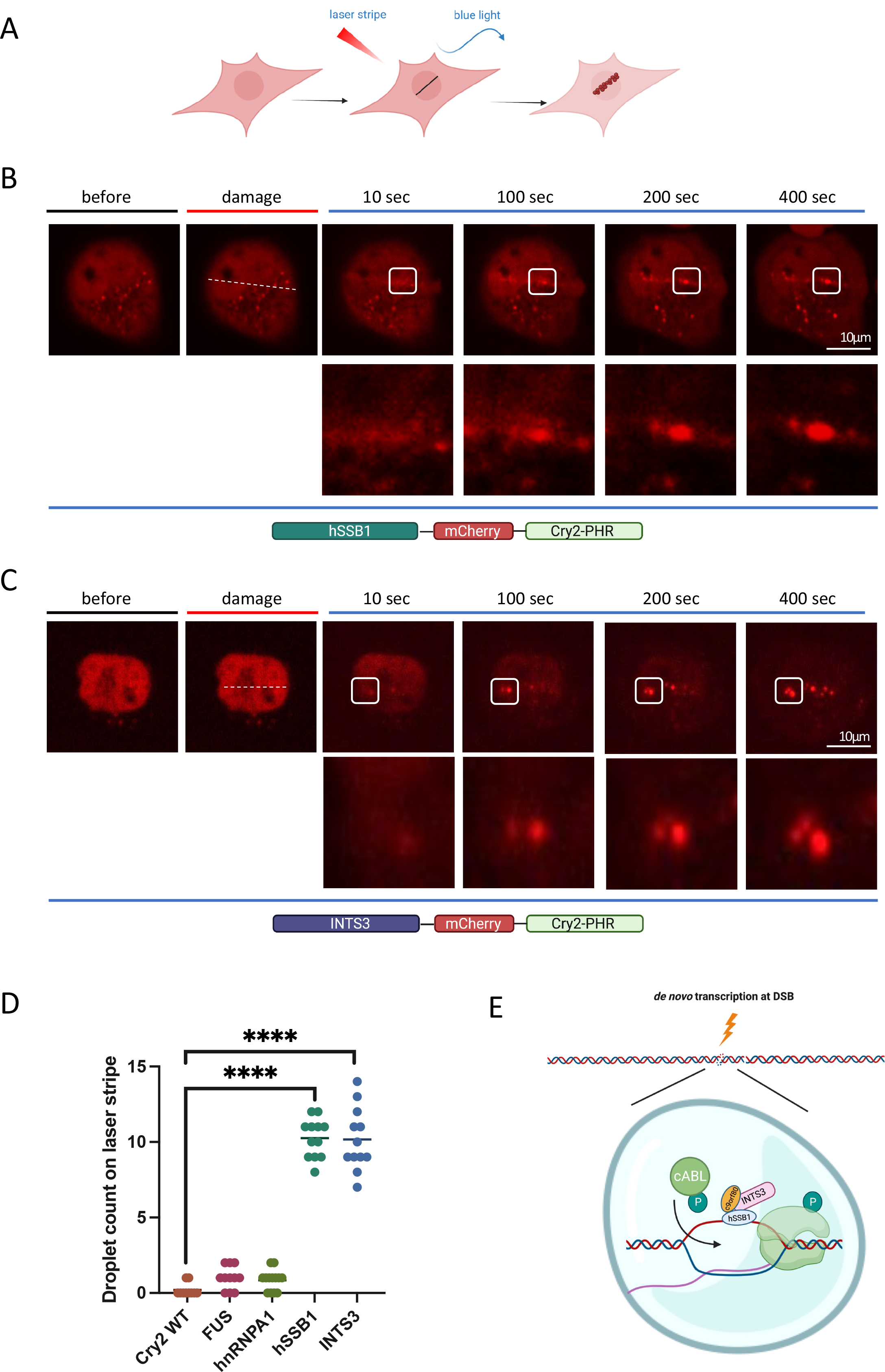
SOSS1 phase separates at double strand breaks *in vivo* **A)** Diagram showing the optoDroplet strategy combined with laser stripping. For global activation, cells were imaged by use of two laser wavelengths (488 nm for Cry2 activation /560 nm for mCherry imaging) at 25% of laser power, in 10 seconds pulses 40 times. Laser stripe was induced using 405 nm laser, followed by light induction at 488 nm and acquisition at 561 nm. **B-C)** Damage induced optoDroplet formation of hSSB1-Cry2-mCherry (B) and INTS3-Cry2- mCherry (C) cells. Representative images of optoDroplets before and after laser stripe and during light induction at indicated time points. Position of the laser stripe is marked with dashed white line. White rectangles show area of the image that is enlarged below. **D)** Quantification of optoDroplets from 3 independent experiments show values for optoDroplet number on the laser stripe at 400 seconds after light induction. Significance was determined student *t*- test. \*\*\*\**p* ≤ 0.0001. **E)** Model: In response to DNA damage, c-abl phosphorylates hSSB1, which forms trimeric SOSS1 complex with INTS3 and c9orf80, binding to Y1P RNAPII via R-loops triggering phase separation at DSBs.

These data suggest that the SOSS1 complex contributes to phase separation at DSBs.

### Trimeric SOSS1 is required for efficient DNA damage repair

Intrigued by these findings, we were wondering whether the trimeric SOSS1 complex is indeed important for DDR. To test this, we monitored γH2AX clearance after IR at several time points in wt and hSSB1 knock-down cells and observed a significant delay in DNA damage repair (fig. S18A). To test which DSB repair pathway the SOSS1 complex might be involved in, we employed reporter cell lines. DR-GFP HR HeLa reporter cells are used to study the HR repair efficiency. The stably integrated DR-GFP cassette has a special designed SceGFP sequence, which contains an I-SceI cutting site followed by a stop codon to forbid the NHEJ and an iGFP sequence was used as the in-frame repair template. Via the transient expression of pCBASceI plasmid, only cells that undergo HR will generate functional GFP, which can be measured by FACS. In this system, we used depletion of BRCA1 as a positive control and observed significant inhibition of HR. Interestingly, we also detected significant reduction in HR efficiency in INTS3 and hSSB1 KD cells (fig. S18B, C and F).

Additionally, we also employed another HeLa-based reporter system, in which the disrupted GFP is re-activated by NHEJ. In this system, we used DNA-PK inhibitor, wortmannin, as a positive control and observed a significant reduction in NHEJ efficiency. The depletion of INTS3 caused only mild inhibition of NHEJ, whilst depletion of hSSB1 led to increased NHEJ efficiency (fig. S18D and F). It is possible that depletion of hSSB1 might be partially compensated by hSSB2. Finally, we performed clonogenic assay and detected a growth defect caused by hSSB1 depletion further exacerbated by IR treatment (fig. S18E).

Collectively, these data show that lack of trimeric SOSS1 complex results in delayed DNA repair.

## DISCUSSION

ATM and DNA-PK mediated phosphorylation of hSSB1 on its T117 and S134 residues implicated its role in DNA repair(*10, 47*). Here, we demonstrate that, upon IR, c-Abl specifically phosphorylates hSSB1 on residues Y102, Y115, Y74 and Y85 (**Fig. 4**). The residues Y74 and Y85 are located within the OB-fold domain, mediating ssDNA binding(*40*). The Y102 is located inside the binding interphase with INTS3, possibly functioning in the trimeric SOSS1 complex assembly. The Y115 residue is present in the unstructured domain. (**Fig 4E**, fig. S10E). c-Abl present at DSBs(*24*) exhibits multiple roles in DDR(*23*). However, its phosphorylation of hSSB1 has not been identified yet. In our study, we show that the phosphorylation of hSSB1 by cAbl is critical for its presence at DSBs, which emphasises the importance of c-Abl and hSSB1 in the repair of DSBs. The phosphorylated hSSB1 works as a signal transducer, initiating the assembly of the trimeric SOSS1 complex at DSBs. This aligns with previously reported role of c-Abl in Y1P CTD RNAPII phosphorylation at DSBs and places c-Abl in the front line of DSBs signalling proteins, stimulating transcription at DSBs and also activating RNAPII binding complexes, such as the trimeric SOSS1 complex. Our *in vitro* data show that the SOSS1 complex interacts with both non-phosphorylated and active RNAPII. Furthermore, active transcription is associated with R-loop formation. Our EMSA experiments revealed that hSSB1 preferentially binds to R-loops and this binding is not outcompeted by RPA when hSSB1 is part of the trimeric SOSS1 complex (**Fig.3**, fig. S8,9). The question how RPA and hSSB1 co-exist and function together has been discussed in the field with no clear answer so far(*8*). Our data suggest that RPA preferentially binds to exposed single-strand DNA at the ends, whilst hSSB1 binds to R-loops, followed by the assembly of the trimeric SOSS1 in close proximity to RNAPII at DSBs.

Our *in vitro* LLPS assay shows that trimeric SOSS1 itself phase separates into condensates (**Fig.6**, fig. S15). The addition of purified S5,7P-CTD and ssDNA into the pre-formed condensates of SOSS1 leads to increased stability and the size of droplets. These data uncover a novel role for the trimeric SOSS1 complex in promoting the transcription through phase separation. A previous *in vivo* LLPS assay(*31*) reported that the S2P-mediated damage-induced long non-coding RNAs (dilncRNA) synthesised at DSBs could stabilise the condensate formation and are vital for cell repair, however, the detailed signalling remains elusive. We provide evidence to show that the trimeric SOSS1 complex is one of the elements for LLPS at DSBs, most likely through the unstructured domain of INTS3, which enables dynamic transient structural shifts for diverse biological purposes. Interestingly, the depletion of hSSB1 resulted in a significant delay in DNA damage repair (fig. S18).

In a summary, we propose a model (**Fig. 8E**) in which the trimeric SOSS1 complex, phosphorylated on hSSB1 by damage activated c-Abl, binds to RNAPII and R-loops to promote efficient repair of DSBs through liquid-liquid phase separation.

## MATERIALS AND METHODS

### Synthetic RNA/DNA substrates

Oligonucleotides for preparing synthetic fluorescently-labelled (Cy3) RNA/DNA substrates were purchased from Sigma (HPLC purified) and their sequences are available in Table S1. Substrates were prepared by mixing 3 pmol of labelled oligonucleotides with a 3-fold excess of the unlabelled oligonucleotides in the annealing buffer [25 mM Tris-HCl, pH 7.5, 100 mM NaCl, 3 mM MgCl2], followed by initial denaturation at 75 °C for 5 min. Substrates were then purified from a native PAGE gel.

### Plasmids

Fragment of DNA containing the ORF of hSSB1 was cloned into plasmid 2BT (pET His6 LIC cloning vector, Addgene plasmid #29666) via ligation independent cloning (LIC). Fragments of DNA containing the ORFs of hSSB1, INTS3, and c9orf80, respectively, were cloned into plasmid 438B (pFastBac His6 TEV cloning vector with BioBrick Polypromoter LIC subcloning, Addgene plasmid #55219). Constructs 438B-INTS3, 438B-hSSB1, and 438B- c9orf80 were combined using BioBrick Polypromoter LIC subcloning into a single construct enabling co-expression of the three subunits of the trimeric SOSS1 complex from a single virus in insect cells. To fluorescently tag INTS3 and INTS3^1-961^ (INTS3ΔC), the ORFs were cloned into plasmid H6-mCerulean (pET Biotin His6 TEV mCerulean LIC cloning vector, Addgene plasmid #29726). In the second step, the ORFs for the fused, fluorescent-tagged INTS3s- mCerulean were cloned into 438B vector. Analogously, hSSB1 and hSSB11-106 (hSSB1ΔC) were fluorescently tagged in two steps by first cloning the ORFs into plasmid H6-mOrange (pET Biotin His6 mOrange LIC cloning vector, Addgene plasmid #29723) and then into plasmid 438B.

The plasmids enabling co-expression of the fluorescent-labelled trimeric SOSS complexes were assembled identically, as described above. A full list of generated plasmids is available in Table S2. Plasmids 2BT,2BcT, 438B, 438C, H6-mCerulean, and H6-mOrange were purchased directly from QB3 Macrolab (UC Berkeley).

To generate plasmids enabling expression of the kinase module of TFIIH complex in insect cells, the ORFs for CDK7, MAT1, and CCNH were cloned into plasmid 438B and later combined into a single construct. Plasmid enabling expression of cABL^CAT^ (AA 83–534), alongside PTP1b^1-238^ was generously provided by Gabriele Fendrich and Michael Becker at the Novartis Institutes for Biomedical Research, Basel. Plasmid expressing catalytically inactive cABL^CAT^ D363A was generated by site-directed mutagenesis. Plasmids 2BcT-GFP-hCTD and 2BcT-mCherry-hCTD (provided by Katerina Linhartova) were used to express and purify the full-length C-terminal domain of the catalytic subunit of RNAPII (hCTD) fused with msfGFP and mCherry respectively. Plasmid pGEX4T1-(CTD)26-(His)7 (provided by Olga Jasnovidova) was used to express and purify GST-(CTD)26-(His)7. All constructs (listed in Table S3) were verified by sequencing.

### Insect cell work

To generate viruses enabling the production of proteins in insect cells, the coding sequences and the necessary regulatory sequences of the constructs were transposed into bacmid using *E. coli* strain DH10bac. The viral particles were obtained by transfection of the bacmids into the *Sf*9 cells using FuGENE Transfection Reagent and further amplification. Proteins were expressed in 300 ml of Hi5 cells (infected at 1×10^6^ cells/ml) with the corresponding P1 virus at a multiplicity of infection >1. The cells were harvested 48 hours post-infection, washed with 1x PBS, and stored at -80°C.

### Protein purification

#### Purification of hSSB1

Five grams of *E. coli* BL21 RIPL cells expressing hSSB1 were resuspended in ice-cold lysis buffer [50 mM Tris-HCl, pH 8; 0.5 M NaCl; 10 mM imidazole; 1 mM DTT], containing protease inhibitors (0.66 μg/ml pepstatin, 5 μg/ml benzamidine, 4.75 μg/ml leupeptin, 2 μg/ml aprotinin) at +4°C. Cells were opened up by sonication. The cleared lysate was passed through 2 mL of Ni-NTA beads (Qiagen), equilibrated with buffer [50 mM Tris-HCl, pH 8; 500 mM NaCl; 10 mM imidazole; and 1 mM DTT]. hSSB1 was eluted with an elution buffer [50 mM Tris-HCl, pH 8; 500 mM NaCl; 1 mM DTT and 400 mM imidazole]. The elution fractions containing hSSB1 were pooled, concentrated, and further fractioned on Superdex S-75 column equilibrated with SEC buffer [25 mM Tris-Cl pH7.5; 200 mM NaCl, 1 mM DTT]. Fractions containing pure hSSB1 were concentrated, glycerol was added to a final concentration of 10 % before they were snap-frozen in liquid nitrogen, and stored at −80 °C.

#### Purification of INTS3

Pellets of Hi5 insect cells were resuspended in ice-cold lysis buffer [50 mM Tris pH 8.0; 500 mM NaCl; 0.4% Triton X-100; 10% (v/v) glycerol; 10 mM imidazole; 1 mM DTT; protease inhibitors (0.66 μg/ml pepstatin, 5 μg/ml benzamidine, 4.75 μg/ml leupeptin, 2 μg/ml aprotinin); and 25 U benzonase per ml of lysate]. The resuspended cells were gently shaken for 10 min at 4°C. To aid the lysis, cells were briefly sonicated. The cleared lysate was passed through 2 mL of Ni-NTA beads (Qiagen), equilibrated with buffer [50 mM Tris-HCl, pH 8; 500 mM NaCl; 10 mM imidazole; and 1 mM DTT]. Proteins were eluted with an elution buffer [50 mM Tris-HCl, pH 8; 500 mM NaCl; 1 mM DTT and 400 mM imidazole]. The elution fractions containing proteins were pooled, concentrated, and further fractioned on Superdex S- 200 column equilibrated with SEC buffer [25 mM Tris-Cl pH7.5; 200 mM NaCl, 1 mM DTT]. Fractions containing pure INTS3 and MBP-INTS6 respectively were concentrated, glycerol was added to a final concentration of 10 % before they were snap-frozen in liquid nitrogen, and stored at −80 °C.

#### Purification of the trimeric SOSS1 complex

Pellets of Hi5 insect cells were resuspended in ice-cold lysis buffer [50 mM Tris pH 8.0; 500 mM NaCl; 0.4% Triton X-100; 10% (v/v) glycerol; 10 mM imidazole; 1 mM DTT; protease inhibitors (0.66 μg/ml pepstatin, 5 μg/ml benzamidine, 4.75 μg/ml leupeptin, 2 μg/ml aprotinin); and 25 U benzonase per ml of lysate]. The resuspended cells were gently shaken for 10 min at 4°C. To aid the lysis, cells were briefly sonicated. The cleared lysate was passed through 2 mL of Ni-NTA beads (Qiagen), equilibrated with buffer [50 mM Tris-HCl, pH 8; 500 mM NaCl; 10 mM imidazole; and 1 mM DTT]. Proteins were eluted with an elution buffer [50 mM Tris-HCl, pH 8; 500 mM NaCl; 1 mM DTT and 400 mM imidazole]. The elution fractions containing proteins were pooled, concentrated, and further fractioned on Superose 6 column equilibrated with SEC buffer [25 mM Tris-Cl pH7.5; 200 mM NaCl, 1 mM DTT]. Fractions containing pure complexes were concentrated, glycerol was added to a final concentration of 10 % before they were snap-frozen in liquid nitrogen, and stored at −80 °C.

#### Purification of proteins for the in vitro LLPS assays

The trimeric SOSS1 complexes (labelled or not) were purified as described above, with the following modification: affinity tags were cleaved-off by TEV protease, followed by reverse Ni-NTA affinity chromatography. Additionally, proteins were frozen in the absence of glycerol.

#### Purification of kinases

The kinase module of TFIIH was purified as described for the trimeric SOSS1 complexes. cABL^CAT^ w.t. and D363A mutant were purified as described for hSSB1.

#### Purification of CTD polypeptides

GST-(CTD)26-(His)7 was purified from *E. coli* cells was purified as described for hSSB1. GFP- hCTD and mCherry-hCTD were purified as described for hSSB1, with the following modification: affinity tags were cleaved-off by TEV protease, followed by reverse Ni-NTA affinity chromatography. Proteins were frozen in the absence of glycerol.

#### Purification of RPA

RPA was purified as described in(*48*).

### *In vitro* phosphorylation assay

#### Analytical phosphorylation of hSSB1 and SOSS complex by cABL^CAT^

hSSB1 and GST (both at 5µM) were phosphorylated with increasing concentrations of cABL^CAT^ (0.14, 0.26, and 0.58 µM) or cABL^CAT^ D363A (0.58 µM) in buffer K [25 mM Tris-Cl pH7.5, 5 mM MgCl2, 2 mM ATP, 1 mM DTT] for 30 min at 37°C (final volume 10 µl). Reactions were stopped by adding 2xSDS loading dye and boiling at 95°C for 5 min. Samples were subsequently analysed on a 12% SDS-PAGE gel. The presence of modification was detected either by western blotting, followed by immunodetection with pan α-pY antibody or by mass-spectrometry (see below).

#### Preparative phosphorylation and purification of CTD polypeptides

Two and half mg of GST-(CTD)26-(His)7, GFP-hCTD, and mCherry-hCTD were phosphorylated by 350 µg of cABL^CAT^ (to phosphorylate Y1 on the CTD) or 250 µg of the kinase module of TFIIH (to phosphorylate S5 and S7 on the CTD) in the presence of 2 mM ATP and 3.5 mM MgCl2 for 60 min at 30°C. Reactions were stopped by placing the reactions at +4°C. CTD peptides were purified from the kinases and ATP by size-exclusion chromatography on Superdex S-200, equilibrated with 25 mM Tris-Cl, pH 7.5; 220 mM NaCl, 1 mM DTT.

### Identification of residues phosphorylated by cABL^CAT^ by mass-spectrometry

The procedure was performed as described earlier(*49*). Briefly, protein samples in the gel pieces were alkylated and digested by trypsin. The digested peptides were extracted from gels. One half of the peptide mixture was directly analysed, and the rest of the sample was used for phosphopeptide enrichment. Both peptide mixtures were separately analysed on LC-MS/MS system (RSLCnano connected to Orbitrap Exploris 480; Thermo Fisher Scientific).

MS data were acquired in a data-dependent strategy using survey scan (350-2000 m/z). High-resolution HCD MS/MS spectra were acquired in the Orbitrap analyser. The analysis of the mass spectrometric RAW data files was carried out using the Proteome Discoverer software (Thermo Fisher Scientific; version 1.4) with in-house Mascot (Matrixscience, London, UK; version 2.4.1) search engine utilization. The phosphoRS feature and manual check of the phosphopeptide spectrum was used for the localisation of phosphorylation.

### Electrophoretic Mobility-Shift Assay (EMSA)

Increasing concentrations of the tested proteins (22, 44, 88, 167 nM; for RPA the following concentrations were used: 5, 10, 20, 40 nM) were incubated with fluorescently labelled nucleic acid substrates (final concentration 10 nM) in buffer D [25 mM Tris-HCl, pH 7.5, 1 mM DTT, 5 mM MgCl2 and 100 mM NaCl] for 20 min at 37°C. Loading buffer [60 % glycerol in 0.001% Orange-G] was added to the reaction mixtures and the samples were loaded onto a 7.5 % (w/v) polyacrylamide native gel in 0.5 x TBE buffer and run at 75 V for 1h at +4°C. The different nucleic acid species were visualised using an FLA-9000 Starion scanner and quantified in the MultiGauge software (Fujifilm). To calculate the relative amount of bound nucleic acid substrate the background signal from the control sample (without protein) was subtracted using the *band intensity - background* option. Nucleic acid-binding affinity graphs were generated with Prism-GraphPad 7.

In the EMSA experiments assessing the effect of RPA on the binding of hSSB1 and the trimeric SOSS complex, respectively, the substrate (10 nM) was first pre-coated with 10 or 30 nM RPA, respectively, for 10 min at 37°C. Subsequently, increasing concentrations (22, 44, 88 nM) of the tested proteins were incorporated and the reaction mixtures were further incubated for 10 min at 37°C. Reactions were next processed as described above. The statistical significance was determined by unpaired *t*-test analysis.

### *In vitro* pull-down experiments

Purified GST, GST-CTD, GST-Y1P-CTD, or GST-S5,7P-CTD (5 μg each), respectively, were incubated with the trimeric SOSS1 complex (5 μg) in 30 μl of buffer T [20 mM Tris–HCl, 200 mM NaCl, 10 % glycerol, 1 mM DTT, 0.5 mM EDTA, and 0.01% Nonidet P-40; pH 7.5] for 30 min at 4°C in the presence of GSH-beads. After washing the beads twice with 100 μl of buffer T, the bound proteins were eluted with 30 μl of 4xSDS loading dye. The input, supernatant, and eluate, 7 μl each, were analysed on SDS-PAGE gel.

### Micro-scale thermophoresis (MST)

Binding affinity comparisons via microscale thermophoresis were performed using the Monolith NT.115 instrument (NanoTemper Technologies). The CTD polypeptides (CTD, Y1P-CTD and S5,7P CTD, respectively) were fused with msfGFP and served as ligands in the assays. Affinity measurements were performed in the MST buffer [25 mM Tris–HCl buffer, pH 7.5; 150 mM NaCl; 1 mM DTT; 5% glycerol; and 0.01% Tween-20]. Samples were soaked into standard capillaries (NanoTemper Technologies). Measurements were performed at 25°C, 50% LED, medium IR-laser power (laser on times were set at 3 s before MST (20 s), and 1 s after), constant concentration of the labelled ligand (20 nM), and increasing concentration of the trimeric SOSS complex (4.8–1200 nM, CTD-GFP and Y1P-CTD-GFP; 28.7-7250 nM, S5,7P CTD). The data were fitted with Hill Slope in GraphPad Prism software.

### *In vitro* LLPS assays

Condensate formation assays were performed in the buffer H [25 mM HEPES, pH 7.5; 220 mM NaCl; 0.5 mM TCEP] in the presence of a crowding agent (5% PEG-8000). Where indicated, ssDNA or ssRNA was added to a final concentration 2 µM. Upon the addition of the indicated proteins (mCherry-CTD peptides at 0.75 µM; the trimeric SOSS1 complex at 0,75, 1, 1,5, and 3µM), the mixtures were immediately spotted onto a glass slide, and the condensates were recorded on Zeiss Axio Observer Z1 with a 63x water immersion objective. Analyses and quantifications of the micrographs were performed in Cell-profiler(*50*). First, four micrographs (2048 pixels (px) per 2048 px; 1 px= 0.103 µm) per condition and per experiment were analysed. Objects (droplets) were identified based on diameter (4-70 px; 0,413-7.5 µm) and intensity using Otsu’s method for thresholding. Picked objects were further filtered based on shape and intensity. For the filtered objects the area and the object count per picture were calculated. The values for droplets were converted from the px to µm based on the metadata of the micrographs. The data were plotted in GraphPad Prism.

The statistical significance of the object counts per picture was determined by unpaired *t*-test analysis, while for the area, a nested *t*-test was used.

### Cell Culture and Cell Line Generation

Cells were cultured at 37 °C with 5% CO2 in high-glucose DMEM medium (Life Technologies, 31966047) supplemented with 10% (vol/vol) fetal bovine serum (FBS) (Merck, F9665), 2 mM L-glutamine (Life Technologies, 25030024) and 100 units/ml penicillin-streptomycin solution (Life Technologies, 15140122). Cell morphology was frequently assessed via microscopy, and regular mycoplasma authentication was conducted. HeLa cells were obtained from ATCC. The stable wild-type hSSB1-GFP and 102A/115A/102&115A hSSB1-GFP mutants were generated with Lipofectamine 3000 (Invitrogen, L3000001) transfection followed by 500 μg/ml hygromycin B (Gibco, 10687010) selection for 10 days. Single-cell sorting was performed to ensure monoclonal-based growth in a 96-well plate (supplemented with 1:1 conditioned HeLa media to fresh media). HeLa HR/NHEJ reporter cell lines were generated with linearized DRGFP and EJ5 cassettes via Lipofectamine LTX (Invitrogen, 15338100) transfection followed by 2 μg/ml puromycin selection for 2 weeks before being single-cell sorted. Monoclonals were progressively grown until sufficient confluency. The correct cassette integration was validated by transfecting the I-SceI overexpression plasmid (Addgene, 26477) for 48 h and measuring GFP induction by flow cytometry. The colony with the highest GFP signal was further validated with western blot by siRNA ablation of the target proteins. To produce stable optoDroplet cell lines expressing Cry2 fusion constructs, lentiviral constructs were transfected with Lipofectamine 3000 (Invitrogen, L3000001) into 293T cells and incubated at 37 °C, 5% CO2 for 48h. Viral supernatants were collected 48h after transfection and filtered with 0.45 μm syringe filters (Sigma, SLHV033R). HeLa cells seeded at ∼70% confluency, were infected by adding 1 ml of filtered viral supernatant directly to the cell medium. Viral medium was replaced with normal growth medium 48 h after infection. The list of cell line is shown in Table S9.

### Chemicals and Antibodies

The DNA damage was generated with γ-rays by CS-137 source (Gravatom, RM30/55). Cells were treated with 1μM cAbl inhibitor Imatinib (Stratech Scientific, B2171-APE-10mM) for 1h, 100μM 5,6-Dichloro-1-beta-D-ribofuranosylbenzimidazole (DRB) (Cayman Chemical, 10010302) for 2h, 1μM THZ1 (Stratech Scientific, A8882-APE-10mM) for 2h prior to the induction of DNA damage, and cells were harvested 10 min post-irradiation (IR) unless stated differently. The antibody list is shown in Table S4.

### Transfection of siRNA and Plasmids

RNAi was performed with Lipofectamine RNAiMax (Life technologies, 13778075), delivered at 30 or 60nM final concentration by using reverse transfection with 1 × 10^6^ cells. The used siRNAs are listed in Table S5. Plasmids delivery was achieved with Lipofectamine 3000 or Lipofectamine LTX with the forward transfection. The details of plasmids source and usage are listed in Table S6. For site-directed mutagenesis, pCMV3-hSSB1-GFP plasmid from Sino Biological (HG22790-ACG-SIB-1Unit) was amplified with Q5® Hot Start High-Fidelity DNA Polymerase (NEB, M0493L) with primers listed in Table S7. PCR products were circularized with T4 kinase (NEB, M0201L) and T4 ligase (NEB, M0202S). The parental plasmid was digested with 5U DpnI (NEB, R0176S). Plasmid transformation was achieved by using the heat shock method (42°C, 47s) in DH5α competent cells (NEB, C2987H), then purified with QIAGEN Plasmid *Plus* Midi Kit (QIAGEN, 12943). Mutations were confirmed by sanger sequencing. For constructing optoDroplet plasmids, Gibson assembly method was applied. DNA fragments encoding human hSSB1 and INTS3 were amplified by PCR from NABP2- GFPspark plasmid (Sino Biological, HG22790-ACG) and INTS3- GFPspark plasmid (Sino Biological, HG15926-ACG) with primers listed in Table S8, then inserted into PHR-mCh- CryWT plasmid (Adgene, 101221) by using NEBuilder HiFi DNA Assembly Cloning Kit (NEB, E5520S). The generated constructs were fully sequenced to confirm the absence of any mutations or stop codons. Control plasmids containing IDRs from FUS or hnRNPA1 were purchased from Adgene (101223, 101226 respectively).

### In situ Proximity Ligation Assay (PLA)

Duolink™ In Situ Red Starter Kit Mouse/Rabbit (Merck, DUO92101-1KT) was used to detect protein-protein interactions. Cells were fixed by 4% paraformaldehyde (PFA) in PBS (Alfa Aesar, J61899) for 10min, followed by 10 min permeabilization with 0.1% TritonX-100 (Merck, X100-100ML) before blocking with 100μL blocking buffer from the kit for 1h. The specific primary antibodies (listed in Table S4) were diluted in Duolink™ dilution buffer and incubated overnight at 4°C. Following primary antibody incubation, PLA probe incubation, ligation and amplification followed the manufacturer’s instructions. Duolink™ In Situ Probemaker PLUS kit (Merck, DUO92009-1KT) was applied to conjugate PLA oligonucleotides (PLUS) to Y1P rat antibody for use in Duolink® PLA experiments. For the detection of cAbl and R-loop, the pre-extraction with CSK buffer was performed as described previously(*36*). Image acquisition was performed on Olympus FluoView Spectral FV1200 Laser Scanning Microscope (IX83) with 60X oil immersion objective. The red PLA dots was quantified with CellProfiler 4.2.1 with sparkle function. Non-parametrical two-tailed Mann-Whitney *u*-test was applied for PLA analysis. Statistical variability was estimated with the standard deviation (SD) and the significance was established at *p* < 0.05 with Graphpad Prism (Version 9).

### Cell Lysis, Immunoprecipitation and Western Blot

Approximately 1 × 10^7^ cells were lysed in 200 μL lysis buffer [50mM Tris pH 8 (Merck, T6066), 150mM NaCl (Merck, S3014), 1mM EDTA (Merck, E9884), 5mM MgCl2 (Merck,PHR2486), 0.5% NP40 (Merck, I8896-100ML), 1X protease inhibitors (Merck, 11873580001)/1X phosphatase inhibitors (Thermo Fisher, A32961)(PPI)] for 20min at 4°C with vortex every 10min. Cytoplasmic supernatant was collected by centrifuge at 500g, 4°C for 5min. The cell chromatin pellet was resuspended with 200 μL lysis buffer and digested with 2μl per sample Pierce Universal Nuclease for Cell Lysis (Thermo Fisher, 88702) and 1μL per sample Benzonase® Nuclease (Merck, E1014-25KU) for 30min at 4°C on wheel with vigorous pipetting every 10min. The soluble nuclear lysate was collected by 10 min centrifuge at 17000g (4⁰C). 300 μL dilution buffer [50mM Tris pH 8, 150mM NaCl, 1mM EDTA, 5mM MgCl2, 1X PPI) was added to the both cytoplasmic and nuclear lysate before take 50 μL Input. GFP-Trap Magnetic Agarose (Proteintech, gtma-20) were washed 3X in cold dilution buffer before adding to the cell lysate for 2h. After pull-down, GFP-Trap beads were wash with high salt washing buffer [50 mM Tris pH 8, 500 mM NaCl, 1 mM EDTA, 5 mM MgCl2, 1X PPI] twice and low salt washing buffer [50mM Tris pH 8, 150mM NaCl, 1mM EDTA, 5mM MgCl2, 1X PPI] twice before eluted with 1X Laemmli Buffer [62.5 mM Tris pH6.8, 2% sodium dodecyl sulphate (SDS), 2% β-mercaptoethanol, 10% glycerol, 0.005% bromophenol blue] (Alfa Aesar, J61337.AD) and boiled for 10min at 95°C. NuPAGE™ 4 to 12% Bis-Tris midi protein gels (Invitrogen™, WG1402BOX) and 4–15% Mini-PROTEAN® TGX™ precast protein gels (BioRad, 4561083/4561086) were used for the standard western blot process. The details of antibodies are listed in Table S1. Blots were imaged with Amersham™ Hyperfilm™ ECL™ (VWR, 28-9068-35). Band intensity was quantified with ImageJ. Statistical analysis was performed with the two-tailed student’s *t*-test and * is refers to p < 0.05, ** is refers to p < 0.01, and *** is refers to p < 0.001.

### Co-Immunoprecipitation (CoIP)

When cells reach 50-70% confluency (approximately 7-10 million cells) in 15cm dishes, they were washed three times by ice-cold PBS before scrapped into PBS and collected by centrifugation (500g, 4°C, 5min). The cell pellet was lysed in 5 volumes (200-300μL) of lysis buffer [50mM Tris pH 8, 150mM NaCl, 1mM EDTA, 5mM MgCl2, 0.5% NP40, 1X Protease inhibitors and 1X Phosphatase inhibitors (PPI)] plus 2μL per sample Pierce Universal Nuclease for Cell Lysis and 1μL per sample Benzonase® Nuclease (Merck, E1014-25KU) for 30min at 4°C on wheel with vigorous pipetting every 10min. The supernatant was collected with 10 min centrifuge (17000g, 4⁰C). 1.5X volume (300-450 μL) of dilution buffer [50mM Tris pH 8, 150mM NaCl, 1mM EDTA, 5mM MgCl2, 1X PPI] was added to the supernatant. Take 0.1X volume of diluted supernatant for Input. GFP-Trap Magnetic Agarose were washed 3X in cold dilution buffer before adding to the cell lysate. The stably overexpressed GFP-tagged proteins were captured by rotating on 4°C for 1.5h. After pull-down, beads were wash with dilution buffer three times before eluted with 2X Laemmli Buffer [62.5 mM Tris pH6.8, 2% sodium dodecyl sulphate (SDS), 2% β-mercaptoethanol, 10% glycerol, 0.005% bromophenol blue] and boiled for 10min at 95°C.

### HR/NHEJ Reporter Assay with FACS

For HR/NHEJ Reporter Assay, HeLa reporter cells stably expressing DRGFP cassette and EJ5 cassette were used. 1×10^6^ cells were reverse transfected with 60nM siRNA in a well of a 6- well plate. After 24h, 1×10^5^ cells were reseeded into a new 6-well plate and cultured for another 24h. 1.5 µg pCBASceI plasmid (I-SceI endonuclease expression vector) (Addgene, 26477) was transfected into cells via Lipofectamine 3000 for a further 48h before cells were harvested to run FACS. As a NHEJ reporter cell line positive control, 1µM Wortmannin (sigma, W3144- 250UL, a DNA-PK inhibitor) was added to cell culture media after 6h of reseeding and maintained until harvest.

### Laser Microirradiation

2×10^5^ HeLa cells stably expressing wild-type hSSB1-GFP and 102A/115A/102&115A mutants were seeded onto CELLview™ Culture dish (35mm) (Greiner, 627860. After 16h, 10 µM Hoescht 33342 (Thermo Scientific, H3570) was used to sensitize cells for 10 min before laser damage. To abolish R-loop, pFRT-TODestRFP_RNAseH1 (Addgene, 65785) plasmid was transfected into stable hSSB1-GFP cells via Lipofectamine 3000 for 16h before pre-sensitization. For plasmid-based laser microirradiation, 1×10^5^ HeLa cells were seeded and transfected with hSSB1-GFP and mutant plasmids for 16h before Hoescht treatment. The laser microirradiation was performed with Nikon SoRa microscope and cells were maintained at 37 °C and 5% CO2 during the experimental procedures. Laser tracks were made by a 405 pulsedlaser with laser power set to 20% at 80 repetitions. The 488 nm channel was monitored every 4 seconds tracking the GFP intensity.

### *In vivo* optoDroplets Live cell imaging

Stably integrated optoDroplets HeLa cell lines (listed in Table S9) were seeded on the 35-mm glass-bottom dish (CELLview™ Culture dish, Greiner, 627860) and grown overnight in normal growth medium to reach ∼ 50% confluency. All live cell imaging was performed using 60X oil immersion objective (NA 1.4) on an Olympus SoRA spinning disk confocal microscope equipped with a temperature stage at 37°C and CO2 chamber. For global activation, cells were imaged by use of two laser wavelengths (488 nm for Cry2 activation /560 nm for mCherry imaging) at 25% of laser power, in 10 seconds pulses 40 times. For DNA damage induced laser stripping, cells were subjected to incubation with 10 µM Hoechst (Thermo Scientific, H3570) for 30 minutes prior to imaging. Laser stripe was induced using 405 nm laser, followed by light induction at 488 nm and acquisition at 561 nm.

### Clonogenic assay

1000 cells were seeded into a 12-well plate and incubated overnight before being irradiated with 2 Gy. The cells were then cultured for 10-14 days until colonies formed. Subsequently, colonies were fixed and stained with a mixture of 0.5% crystal violet and 20% methanol for 30min. Images were scanned and quantified by ImageJ with the ColonyArea plugin.

### Statistical analyses

Statistical tests were performed in GraphPad Prism 9.3.1 and Excel. All error bars represent mean ± SD, unless stated differently. Each experiment repeats at least 3 times (N=3). Statistical testing was performed using the Student’s *t*-test, one-way ANOVA, Mann–Whitney test (non-parametric comparison for PLA foci analysis). Significance are listed as \**p* ≤ 0.05, \*\**p* ≤ 0.01, \*\*\**p* ≤ 0.001, \*\*\*\**p* ≤ 0.0001.

## ACKNOWLEDGEMENTS

We are grateful to Dr Sue Tan-Wang from Prof. Nicholas Proudfoot’s lab (University of Oxford) for sharing the pRNH1-GFP, pRNH1D210N-GFP and pRNH1WKKD-GFP plasmids; Katerina Linhartova (CEITEC, Masaryk University) for sharing 2BcT-GFP-hCTD and 2BcT- mCherry-hCTD plasmids and for assistance with the analysis of the LLPS experiments; Olga Jasnovidova (University of Tallin) for sharing pGEX4T1-(CTD)26-(His)7 plasmid. We also thank Alan Wainman for his help with microscopy, and Ruth F. Ketley for CoIP protocol.

## AUTHOR CONTRIBUTIONS

Conceptualization: MS, MG

Investigation: QL, MS, AA, ZL, KS, VH, MG Visualization: QL, MS, VH, MG

Funding acquisition: MS, RS, MG Project administration: MS, RS, MG Supervision: MS, MG

Writing – original draft: QL, MS, MG Writing – review & editing: QL, MS, RS, MG

## DATA AND MATERIAL AVAILABILITY

All data are available in the main text or the supplementary materials.

---

**Supplementary Figure 1.**
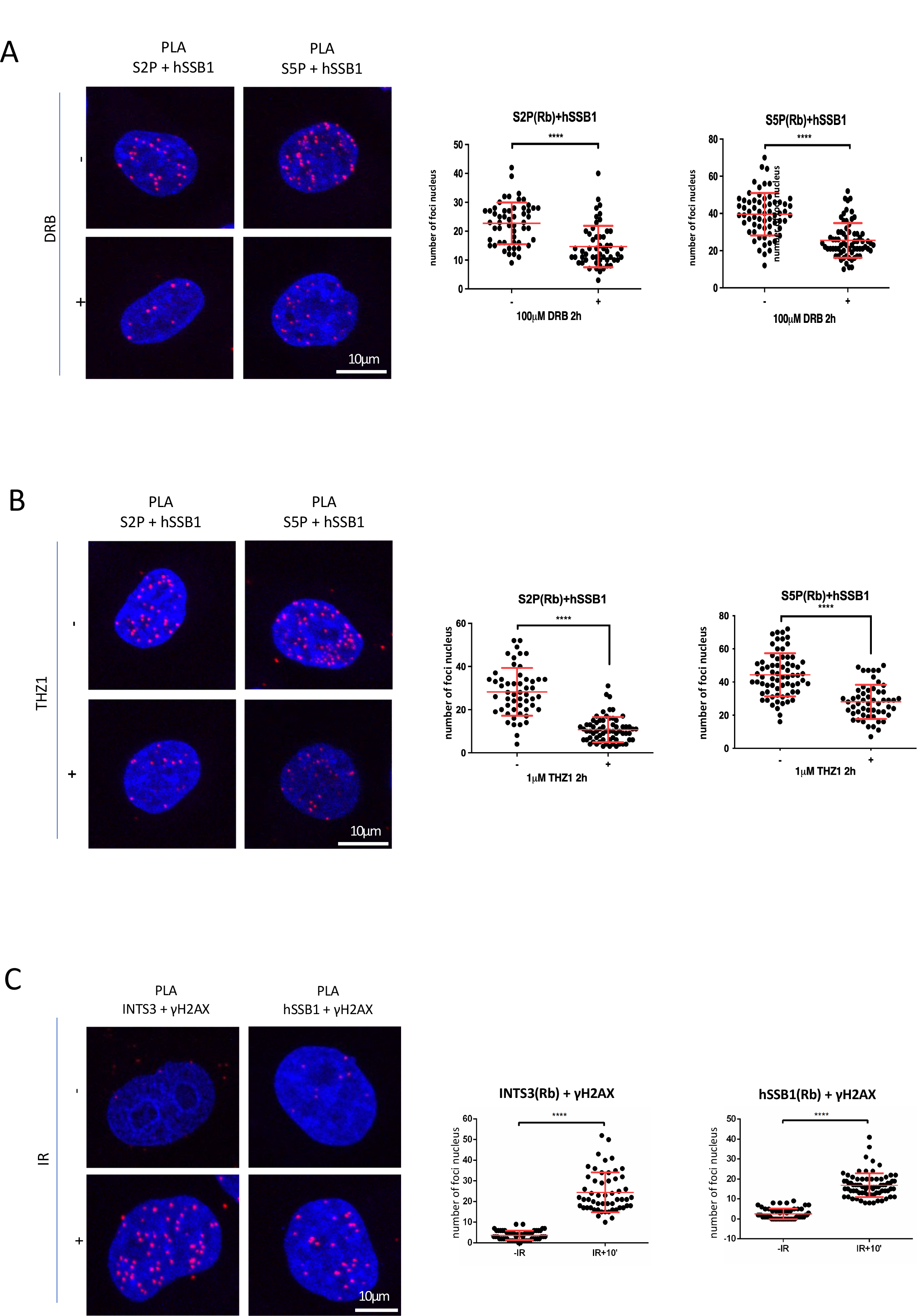
A) PLA of hSSB1 and S2P or S5P in cells with or without DRB treatment (100μM, 2h). Left: representative confocal microscopy images; right: quantification of left, error bar = mean ± SD, significance was determined using non-parametric Mann-Whitney test. \*\*\*\**p* ≤ 0.0001. **B)** PLA of hSSB1 and S2P or S5P in cells with or without THZ treatment (1μM, 2h). Left: representative confocal microscopy images; right: quantification of left, error bar = mean ± SD, significance was determined using non-parametric Mann-Whitney test. \*\*\*\**p* ≤ 0.0001. **C)** PLA of INTS3 or hSSB1 and γH2AX in cells with or without IR. IR=10Gy. Left: representative confocal microscopy images; right: quantification of left, error bar = mean ± SD, significance was determined using non-parametric Mann-Whitney test. \*\*\*\**p* ≤ 0.0001.

**Supplementary Figure 2.**
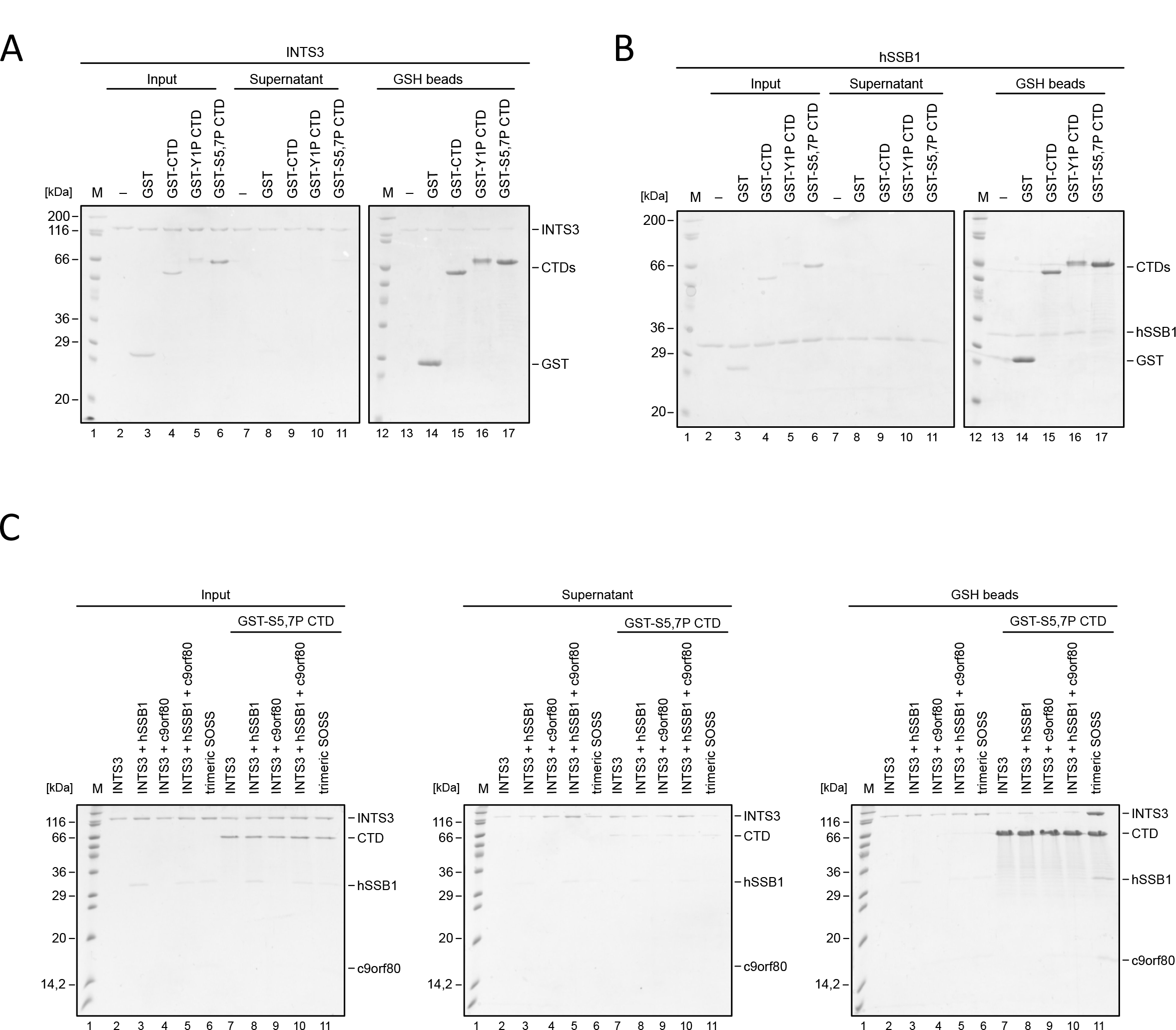
A) A representative SDS-PAGE gel depicting *in vitro* pull-down assay of INTS3 with GST-tagged CTD, GST-tagged CTD modified on tyrosine 1 (Y1P), or GST-tagged CTD modified on serine 5 and 7 (S5,7P). **B)** A representative SDS-PAGE gel depicting *in vitro* pull-down assay of hSSB1 with GST-tagged CTD, GST-tagged CTD modified on tyrosine 1 (Y1P), or GST-tagged CTD modified on serine 5 and 7 (S5,7P). **C)** A representative SDS-PAGE gel depicting *in vitro* pull-down assay of the individual subunits of the SOSS1 complex alone and in combination with GST-tagged CTD, GST-tagged CTD modified on tyrosine 1 (Y1P), or GST-tagged CTD modified on serine 5 and 7 (S5,7P). Purified trimeric SOSS1 complex was used as a positive control.

**Supplementary Figure 3.**
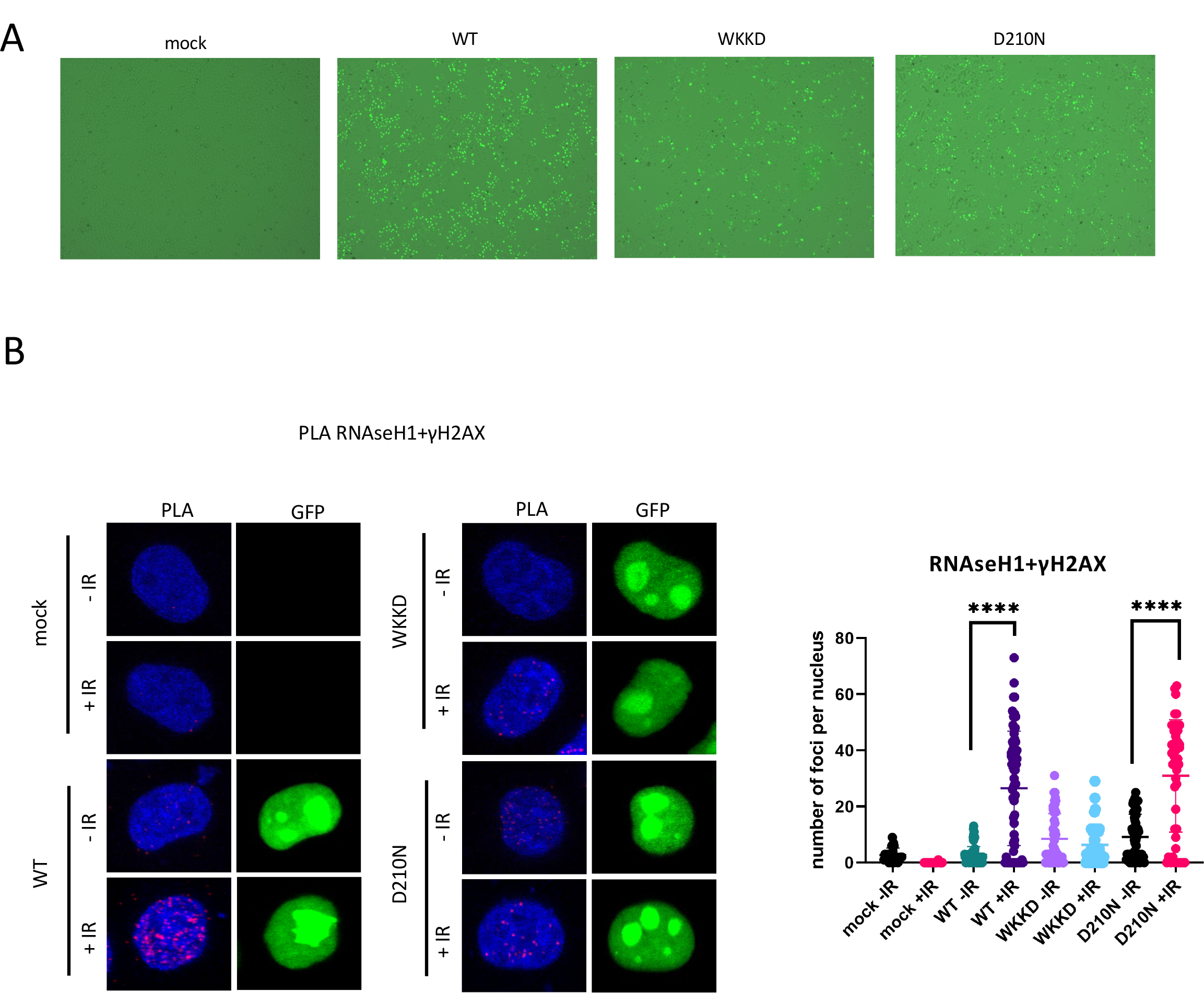
A) Confocal microscopy images showing GFP expression of transiently transfected RNAseH1- GFP wt or WKKD (binding and catalytic) or D210N (catalytic) mutants. Mock is used as negative control. **B)** PLA of RNAseH1-GFP and γH2AX in cells with transiently transfected RNAseH1^wt^-GFP or RNAseH1^WKKD^-GFP (binding and catalytic) or RNAseH1^D210N^-GFP (catalytic) mutants with or without IR. IR=10Gy. Left: representative confocal microscopy images; right: quantification of left, error bar = mean ± SD, significance was determined using non-parametric Mann-Whitney test. \*\*\*\**p* ≤ 0.0001.

**Supplementary Figure 4.**
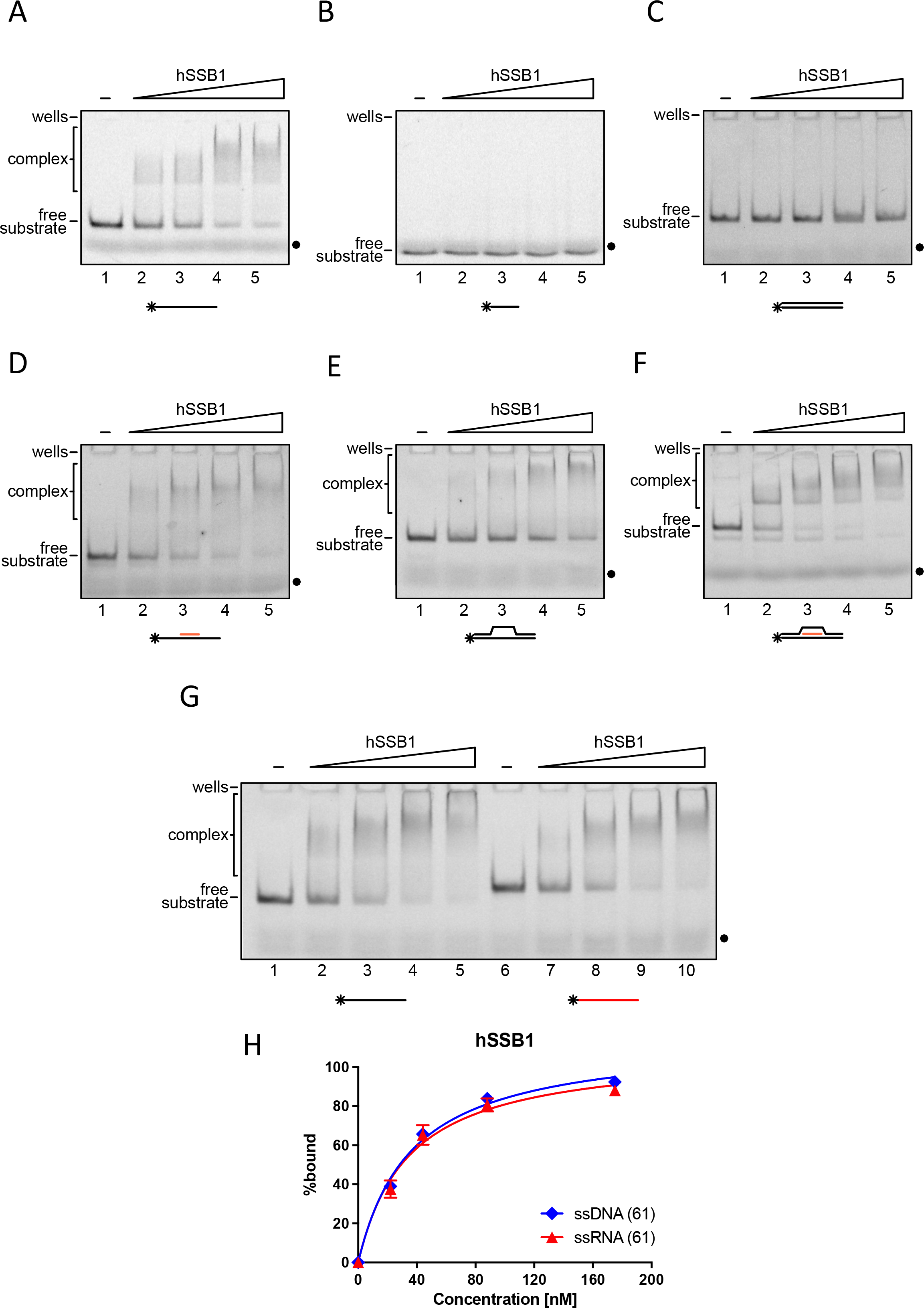
A) Scans of representative EMSA experiments of hSSB1 with 61-mer ssDNA. **B)** Scans of representative EMSA experiments of hSSB1 with 21-mer ssDNA. **C)** Scans of representative EMSA experiments of hSSB1 with 61-mer dsDNA. **D)** Scans of representative EMSA experiments of hSSB1 with RNA:DNA hybrids. **E)** Scans of representative EMSA experiments of hSSB1 with DNA bubble. **F)** Scans of representative EMSA experiments of hSSB1 with R-loops. **G)** Scans of representative EMSA experiments of hSSB1 with 61-mer ssDNA (black) and ssRNA (red). **H)** Graph representing quantification of EMSA experiments from G (n=3).

**Supplementary Figure 5.**
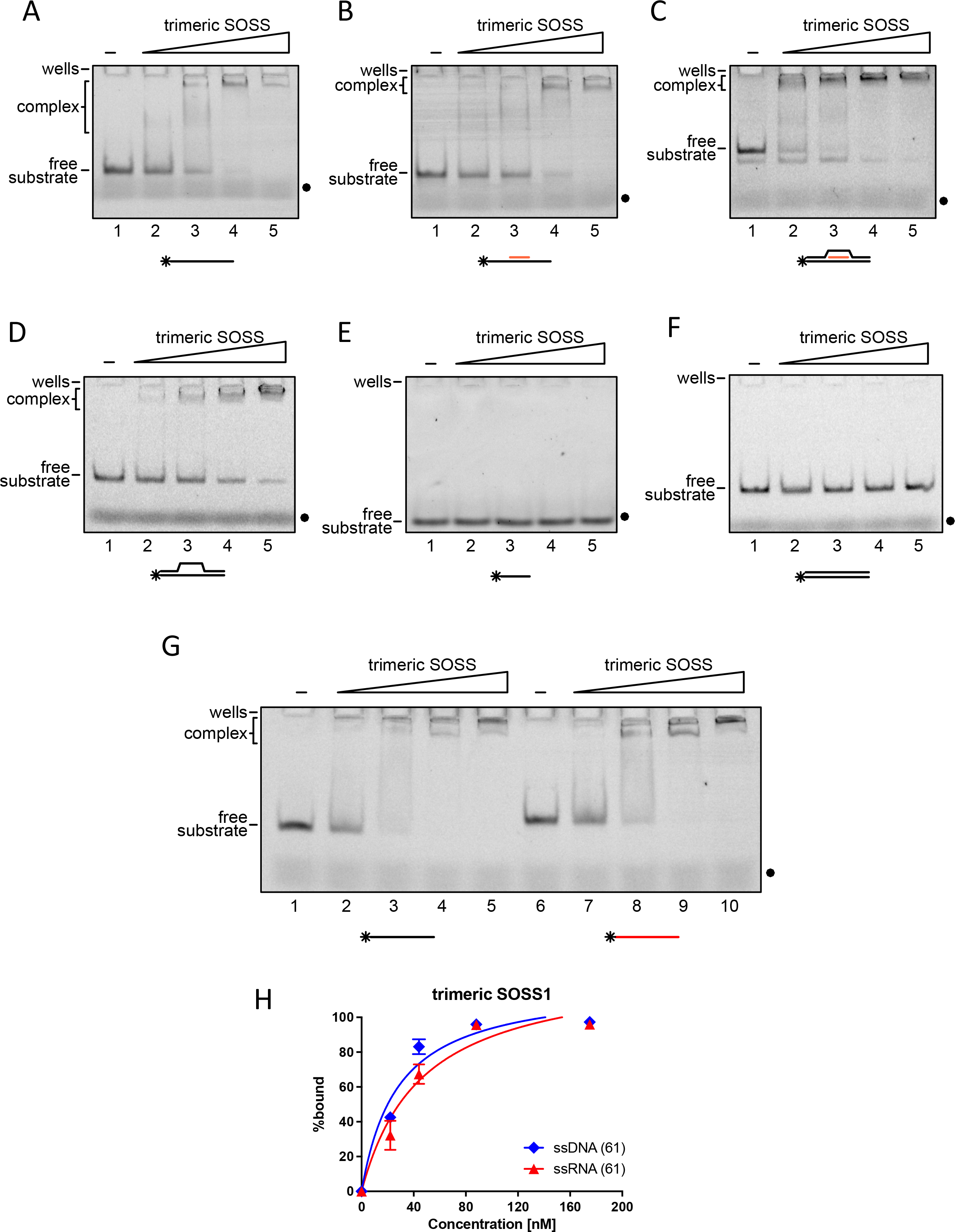
A) Scans of representative EMSA experiments of trimeric SOSS1 with 61-mer ssDNA. **B)** Scans of representative EMSA experiments of trimeric SOSS1 with RNA:DNA hybrids. **C)** Scans of representative EMSA experiments of trimeric SOSS1 with R-loops. **D)** Scans of representative EMSA experiments of trimeric SOSS1 with DNA bubble. **E)** Scans of representative EMSA experiments of trimeric SOSS1 with 21-mer ssDNA. **F)** Scans of representative EMSA experiments of trimeric SOSS1 with 61-mer dsDNA. **G)** Scans of representative EMSA experiments of trimeric SOSS1 with 61-mer ssDNA (black) and ssRNA (red). **H)** Graph representing quantification of EMSA experiments from G (n=3).

**Supplementary Figure 6.**
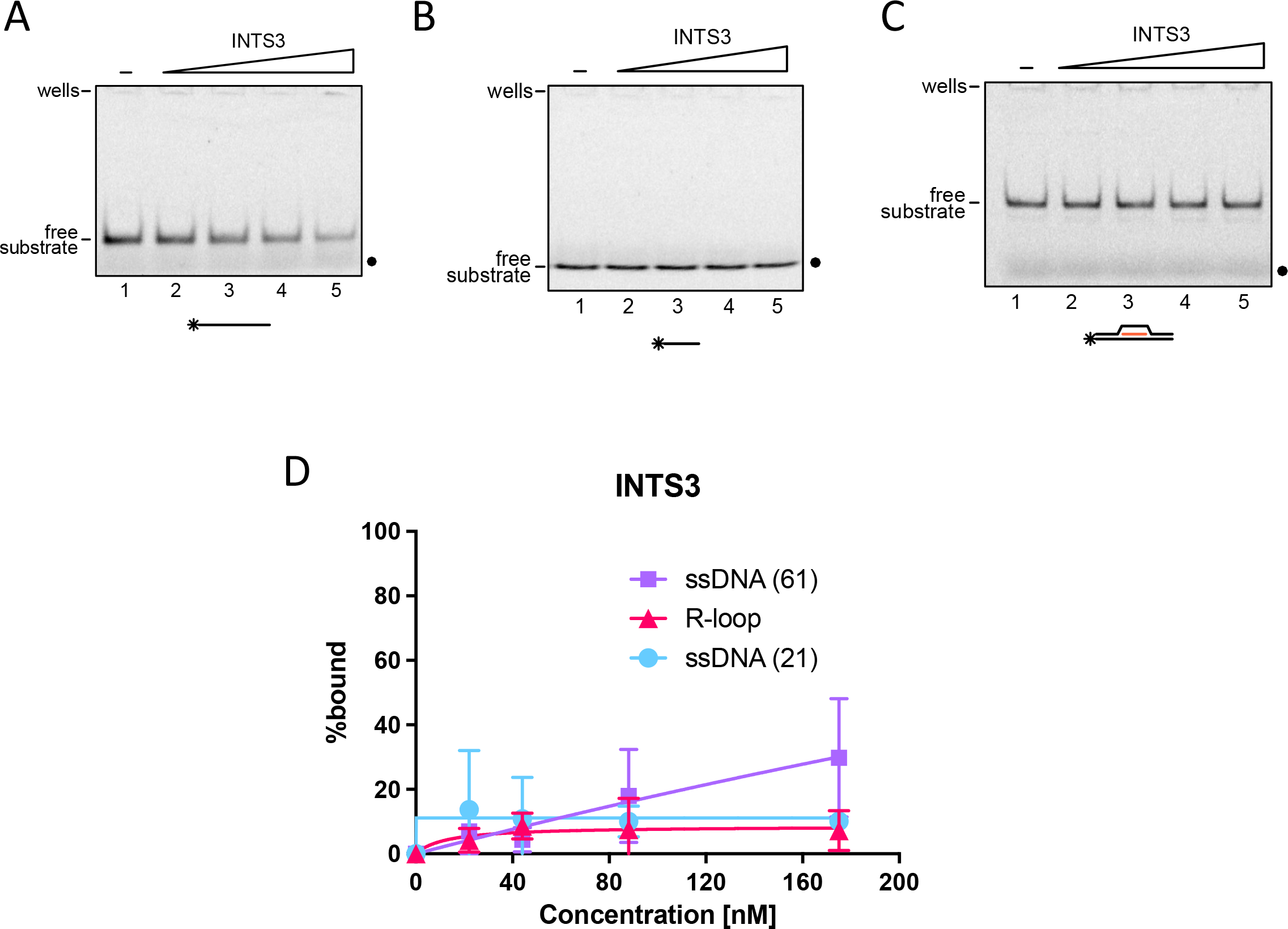
A) Scans of representative EMSA experiments and conducted between INTS3 with 61-mer ssDNA. **B)** Scans of representative EMSA experiments and conducted between INTS3 with 21-mer ssDNA. **C)** Scans of representative EMSA experiments and conducted between INTS3 with R-loop. **D)** Graph representing quantification of EMSA experiments from A-C (n=3).

**Supplementary Figure 7.**
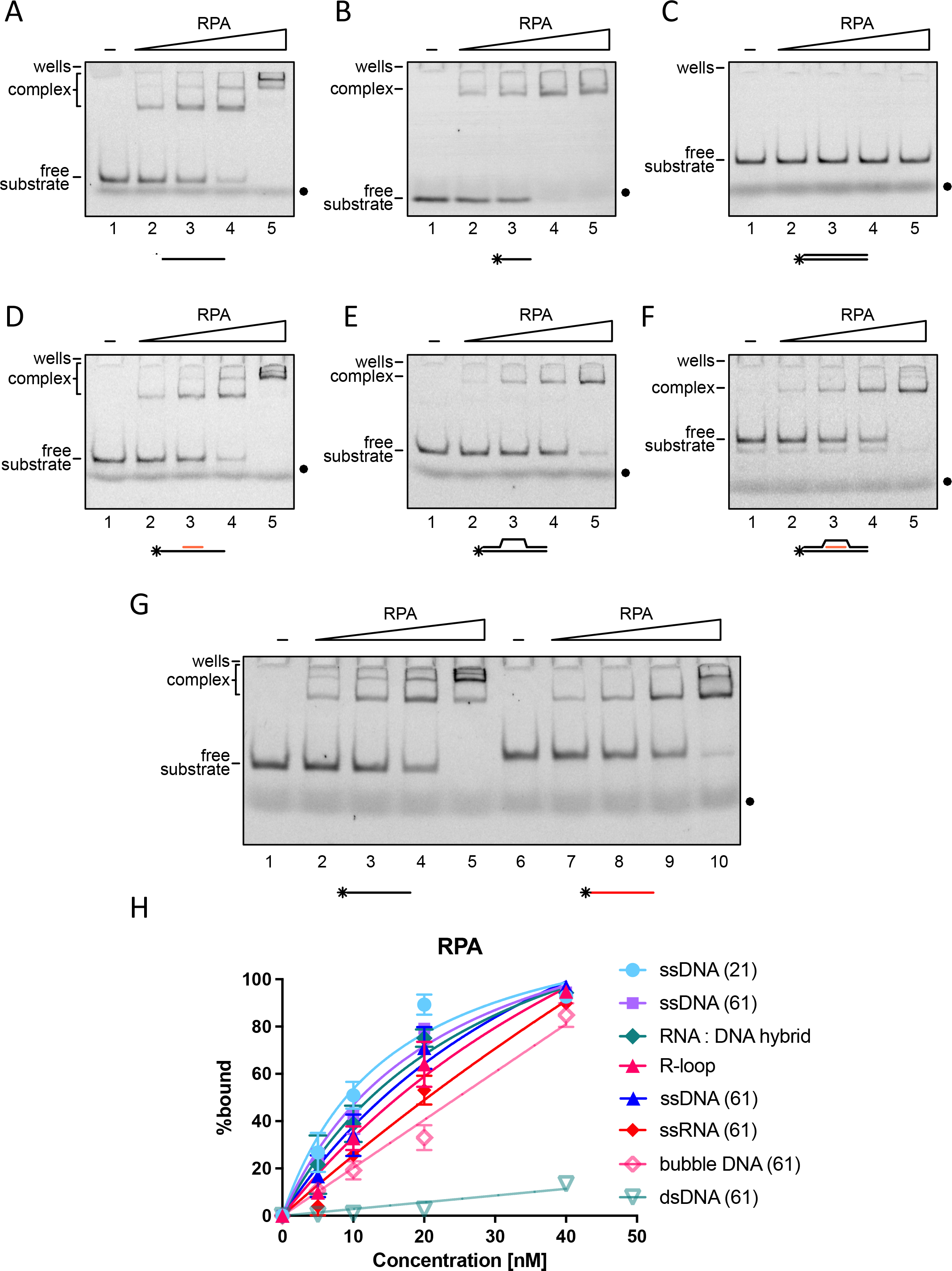
A) Scans of representative EMSA experiments of RPA with 61-mer ssDNA. **B)** Scans of representative EMSA experiments of RPA with 21-mer ssDNA. **C)** Scans of representative EMSA experiments of RPA with 61-mer dsDNA. **D)** Scans of representative EMSA experiments of RPA with RNA:DNA hybrids. **E)** Scans of representative EMSA experiments of RPA with DNA bubble. **F)** Scans of representative EMSA experiments of RPA with R-loops. **G)** Scans of representative EMSA experiments of RPA with 61-mer ssDNA (black) and ssRNA (red). **H)** Graph representing quantification of EMSA experiments from A-G (n=3).

**Supplementary Figure 8.**
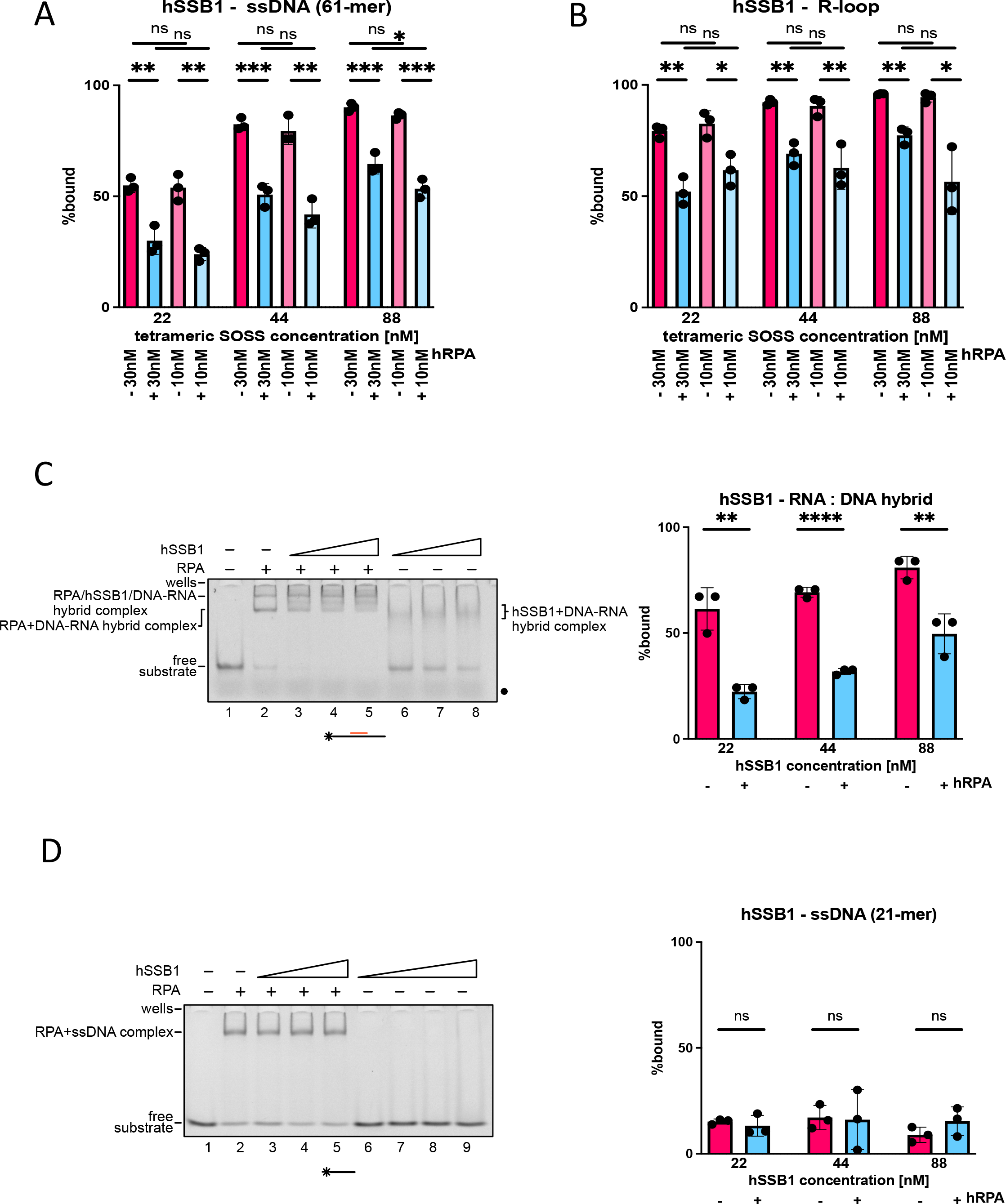
A) Graph representing quantification of EMSA experiments of hSSB1 with 61-mer ssDNA in absence or presence of RPA at various concentrations (n=3). **B)** Graph representing quantification of EMSA experiments of hSSB1 with R-loop in absence or presence of RPA at various concentrations (n=3). **C)** Scans of representative EMSA experiments of hSSB1 with RNA:DNA hybrids in absence or presence of RPA at various concentrations (left) and graph representing quantification of EMSA experiments (n=3) (right). **D)** Scans of representative EMSA experiments of hSSB1 with 21-mer ssDNA in absence or presence of RPA at various concentrations (left) and graph representing quantification of EMSA experiments (n=3) (right).

**Supplementary Figure 9.**
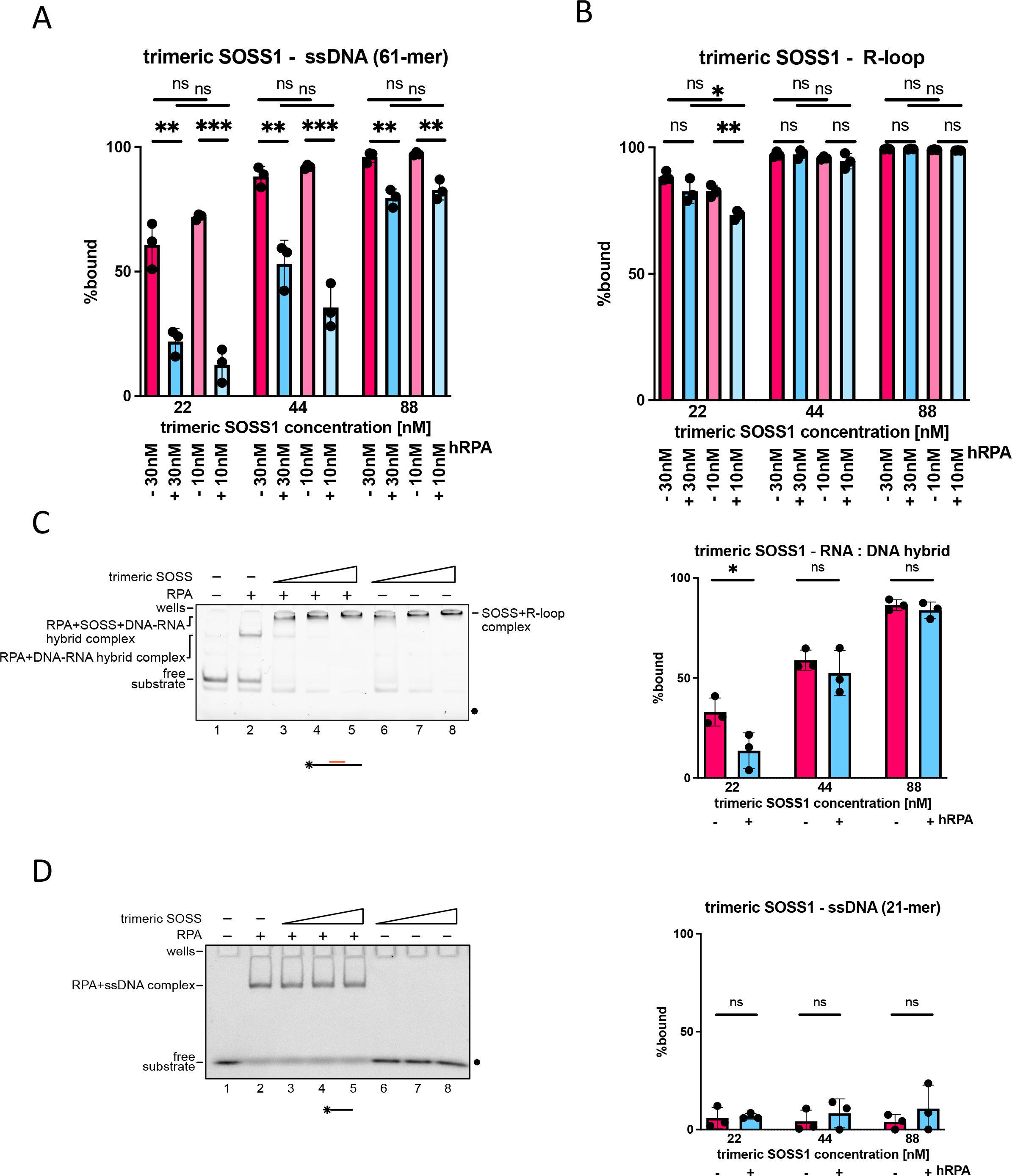
A) Graph representing quantification of EMSA experiments of trimeric SOSS1 with 61-mer ssDNA in absence or presence of RPA at various concentrations (n=3). **B)** Graph representing quantification of EMSA experiments of trimeric SOSS1 with R-loop in absence or presence of RPA at various concentrations (n=3). **C)** Scans of representative EMSA experiments of trimeric SOSS1 with RNA:DNA hybrids in absence or presence of RPA at various concentrations (left) and graph representing quantification of EMSA experiments (n=3) (right). **D)** Scans of representative EMSA experiments of trimeric SOSS1 with 21-mer ssDNA in absence or presence of RPA at various concentrations (left) and graph representing quantification of EMSA experiments (n=3) (right).

**Supplementary Figure 10.**
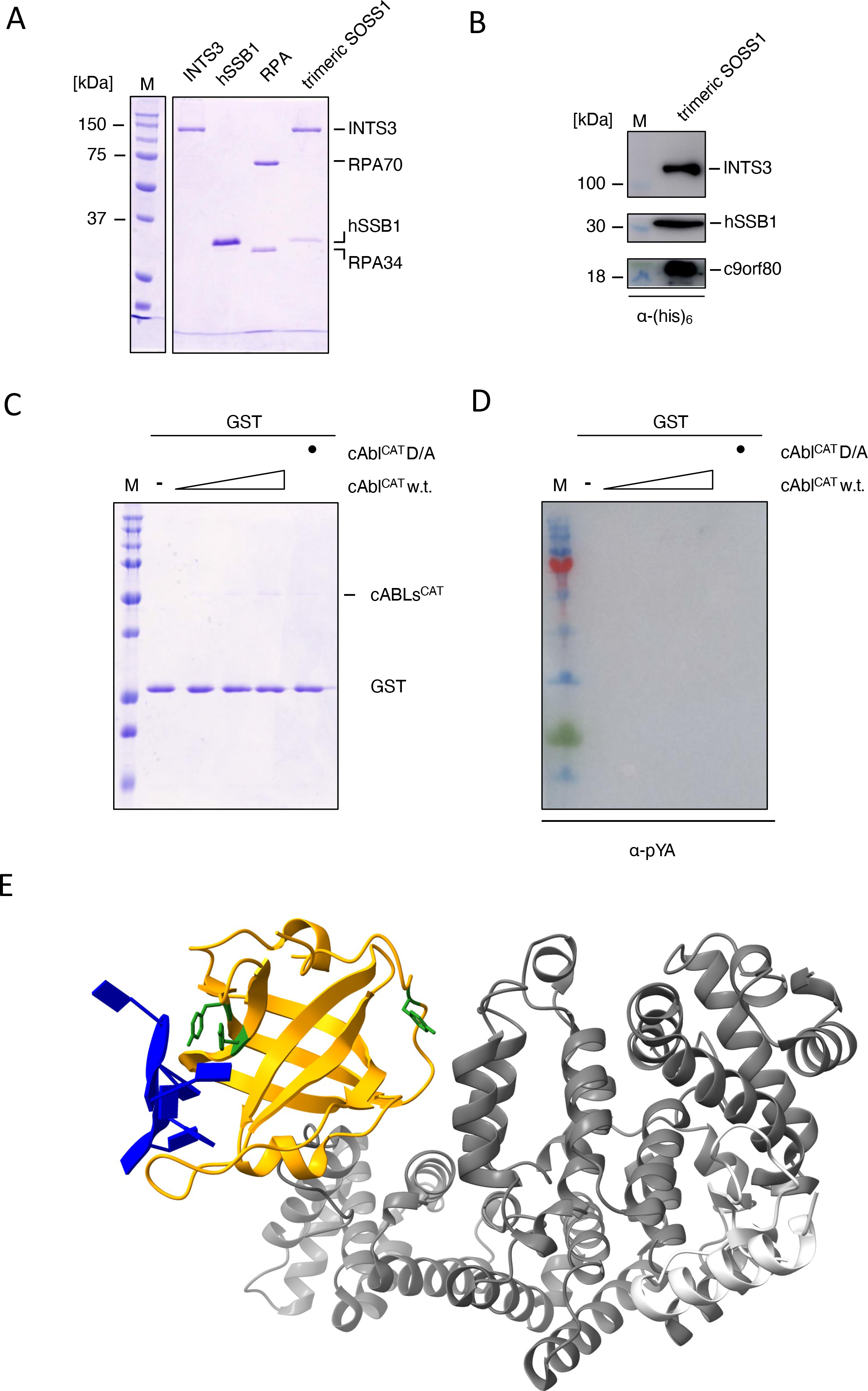
A) An SDS-PAGE gel depicting purified INTS3, hSSB1, RPA, and the trimeric SOSS1 complex. **B)** Western blot detection of the individual subunits of the SOSS1 complex. **C)** *In vitro* phosphorylation of GST by cAbl^CAT^ as depicted by an SDS-PAGE gel of the reaction. **D)** Western blot of samples from C) detected with α-pY antibody. **E)** Depiction of the position of tyrosine residues (in green) of hSSB1 (yellow) on the structural model of the trimeric SOSS1 complex with ssDNA (PDB ID: 4OWW). Residue Y115 is not visible in the structure and could not be highlighted. Dark grey represents INTS3, and light grey represents c9orf80.

**Supplementary Figure 11.**
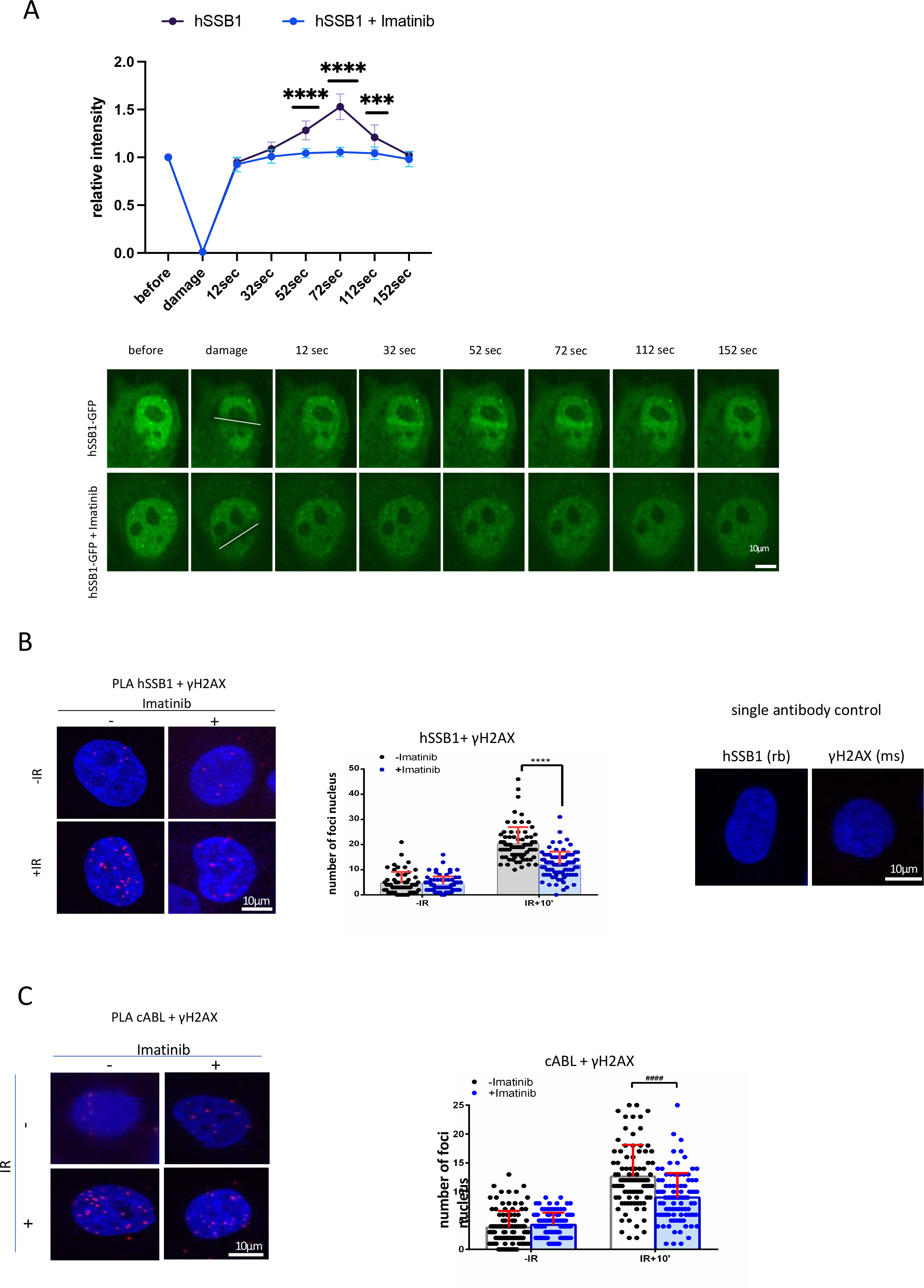
A) Laser stripping of stably integrated hSSB1-GFP cells with and without Imatinib (1μM, 1h). Representative spinning disk confocal microscopy images and quantification (n≥10) showing GFP signals before and after laser striping at indicated time points; error bar = mean ± SEM; significance was determined using multiple unpaired Student’s *t*-test. \*\*\**p* ≤ 0.001 \*\*\*\**p* ≤ 0.0001. Upper panel is the quantification data for bottom representative images. **B)** PLA of hSSB1 and γH2AX in cells without IR, with IR and with IR and Imatinib. IR=10Gy. Cells were treated with 1μM Imatinib for 1h before IR. Left: representative confocal microscopy images; right: quantification of left, error bar = mean ± SD, significance was determined using non-parametric Mann-Whitney test. \*\*\*\**p* ≤ 0.0001. Single antibody was used as negative control. **C)** PLA of cAbl and γH2AX in cells with or without IR. IR=10Gy. Cells were treated with 1μM Imatinib for 1h before IR. Left: representative confocal microscopy images; right: quantification of left, error bar = mean ± SD, significance was determined using non-parametric Mann-Whitney test. ^####^*p* ≤ 0.0001

**Supplementary Figure 12.**
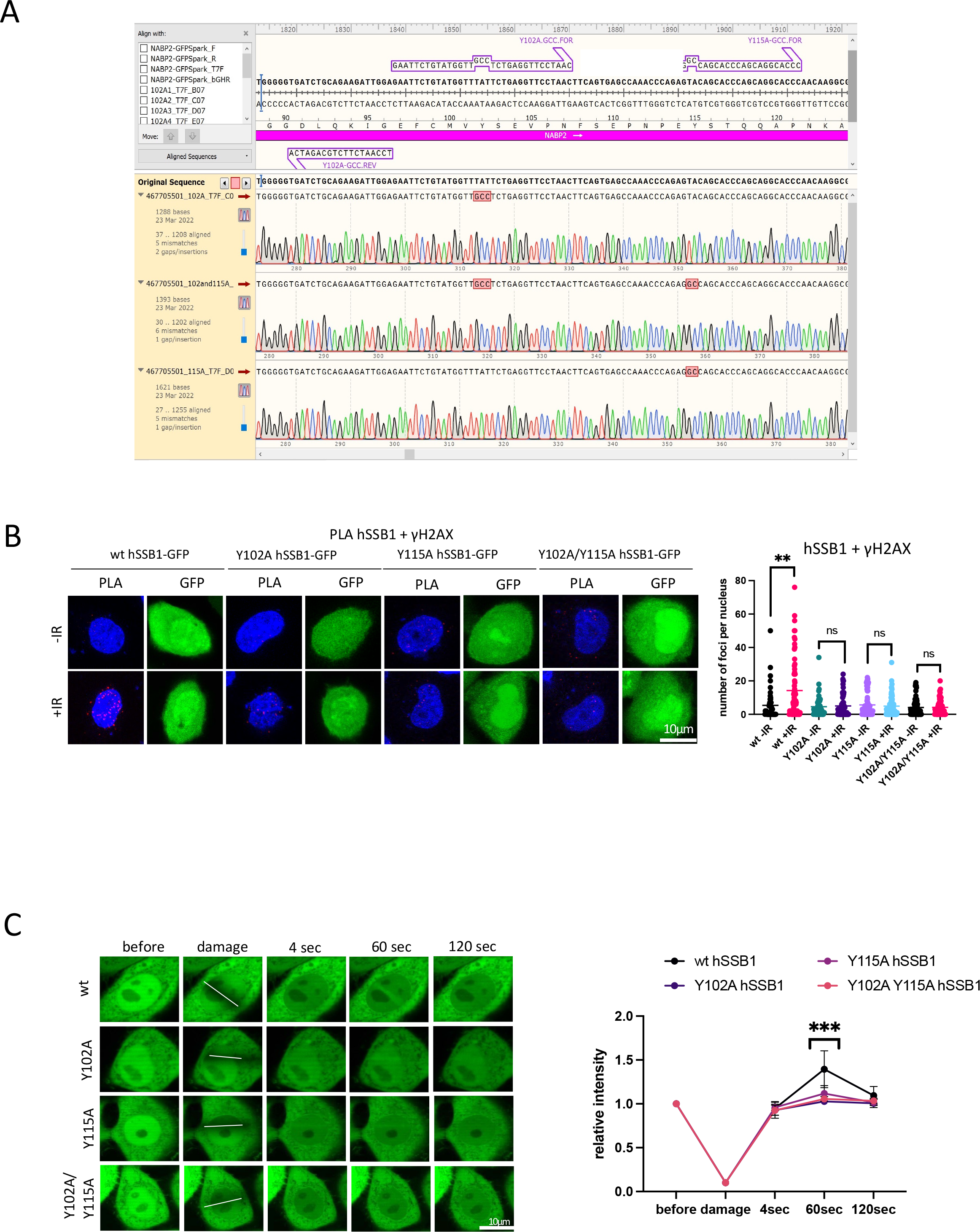
A) Screenshot of sequencing validation corresponding to hSSB1^Y102A^-GFP, hSSB1^Y115A^-GFP and hSSB1^Y102A/Y115A^-GFP plasmids. **B)** PLA of hSSB1 and γH2AX in cells transiently transfected with hSSB1^wt^-GFP or hSSB1^Y102A^-GFP, hSSB1^Y115A^-GFP and hSSB1^Y102A/Y115A^-GFP plasmids treated with or without IR. IR=10Gy. Left: representative confocal microscopy images; right: quantification of left, error bar = mean ± SD, significance was determined using non-parametric Mann-Whitney test. \*\**p* ≤ 0.01. **C)** Laser stripping of cells transiently transfected with hSSB1^wt^-GFP or hSSB1^Y102A^-GFP, hSSB1^Y115A^-GFP and hSSB1^Y102A/Y115A^-GFP plasmids. Representative spinning disk confocal microscopy images and quantification (n≥10) showing GFP signals before and after laser striping at indicated time points; error bar = mean ± SEM; significance was determined using multiple unpaired Student’s *t*-test. \*\*\**p* ≤ 0.001.

**Supplementary Figure 13.**
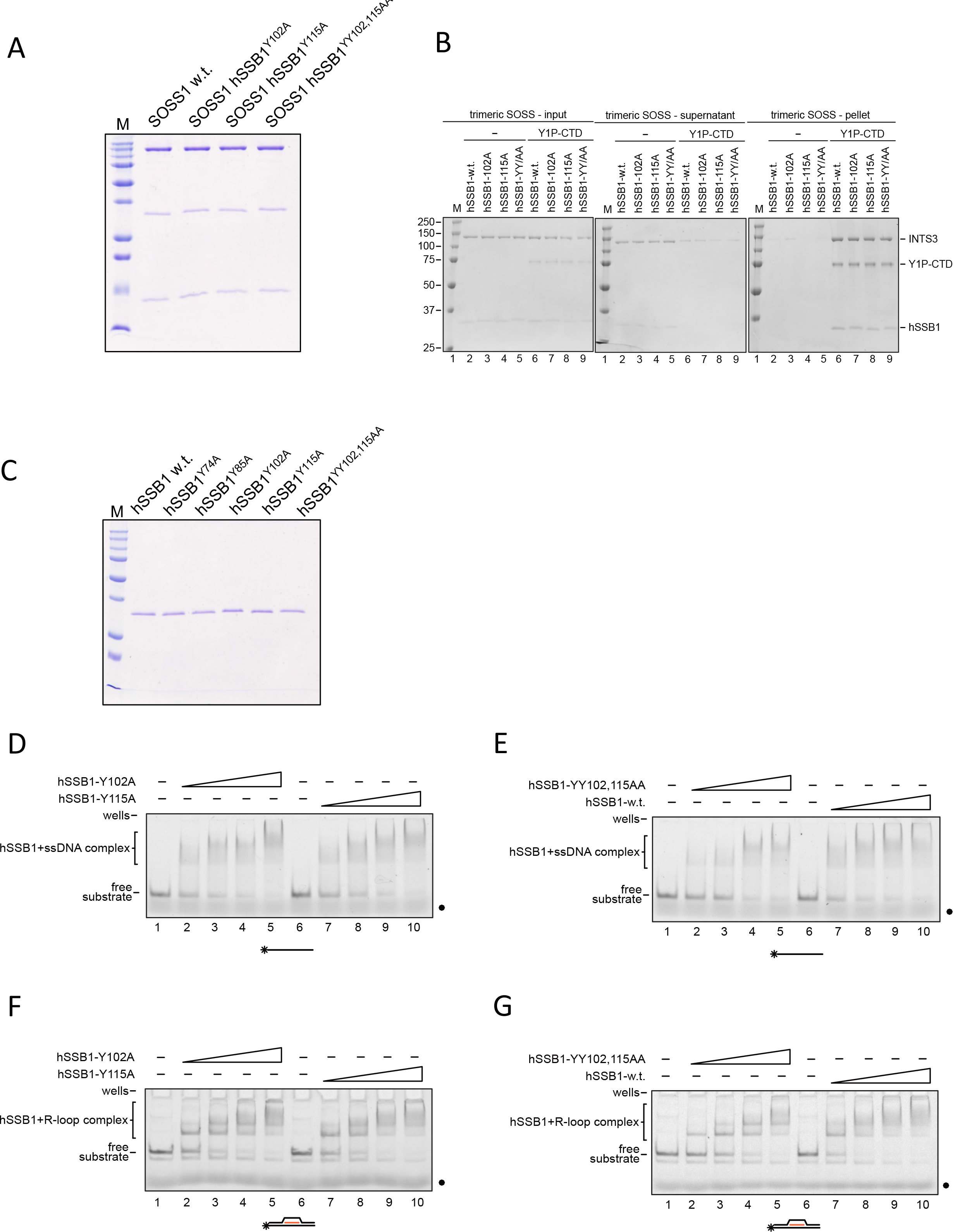
A) An SDS-PAGE gel depicting purified trimeric SOSS1 complexes containing hSSB1 wt, Y102A, Y115A, or YY102,115AA mutants, respectively. **B)** A representative SDS-PAGE gel depicting *in vitro* pull-down assay of purified trimeric SOSS1 complexes containing hSSB1 wt, Y102A, Y115A, or YY102,115AA (YY/AA) mutants with GST-tagged CTD modified on tyrosine 1 (Y1P). **C)** An SDS-PAGE gel depicting purified hSSB1 wt, Y102A, Y115A, or YY102,115AA mutants. **D)** Scans of representative EMSA experiments of hSSB1-Y102A and Y115A mutants with 61- mer ssDNA. **E)** Scans of representative EMSA experiments of hSSB1- YY102,115AA mutant hSSB1 and wt with 61-mer ssDNA. **F)** Scans of representative EMSA experiments of hSSB1-Y102A and Y115A mutants with R-loop. **G)** Scans of representative EMSA experiments of hSSB1- YY102,115AA mutant and hSSB1 wt with R-loop.

**Supplementary Figure 14.**
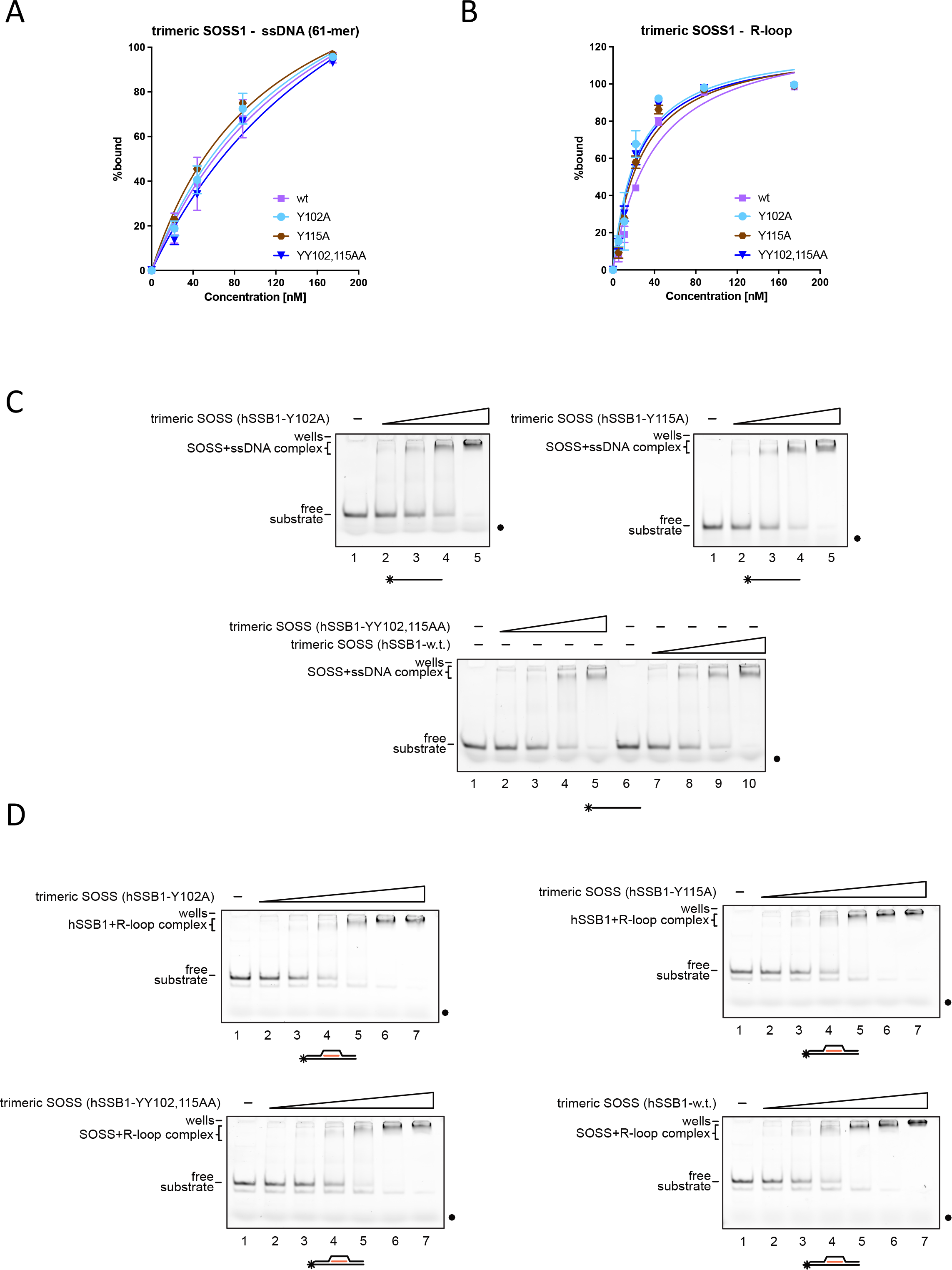
A) Graph representing quantification of EMSA experiments of trimeric SOSS1 containing hSSB1 wt, Y102A, Y115A, or YY102,115AA mutants with 61-mer ssDNA (n=3). **B)** Graph representing quantification of EMSA experiments of trimeric SOSS1 containing hSSB1 wt, Y102A, Y115A, or YY102,115AA mutants with R-loop (n=3). **C)** Scans of representative EMSA experiments of trimeric SOSS1 containing hSSB1 wt, Y102A, Y115A, or YY102,115AA mutants with 61-mer ssDNA. **D)** Scans of representative EMSA experiments of trimeric SOSS1 containing hSSB1 wt, Y102A, Y115A, or YY102,115AA mutants with R-loop.

**Supplementary Figure 15.**
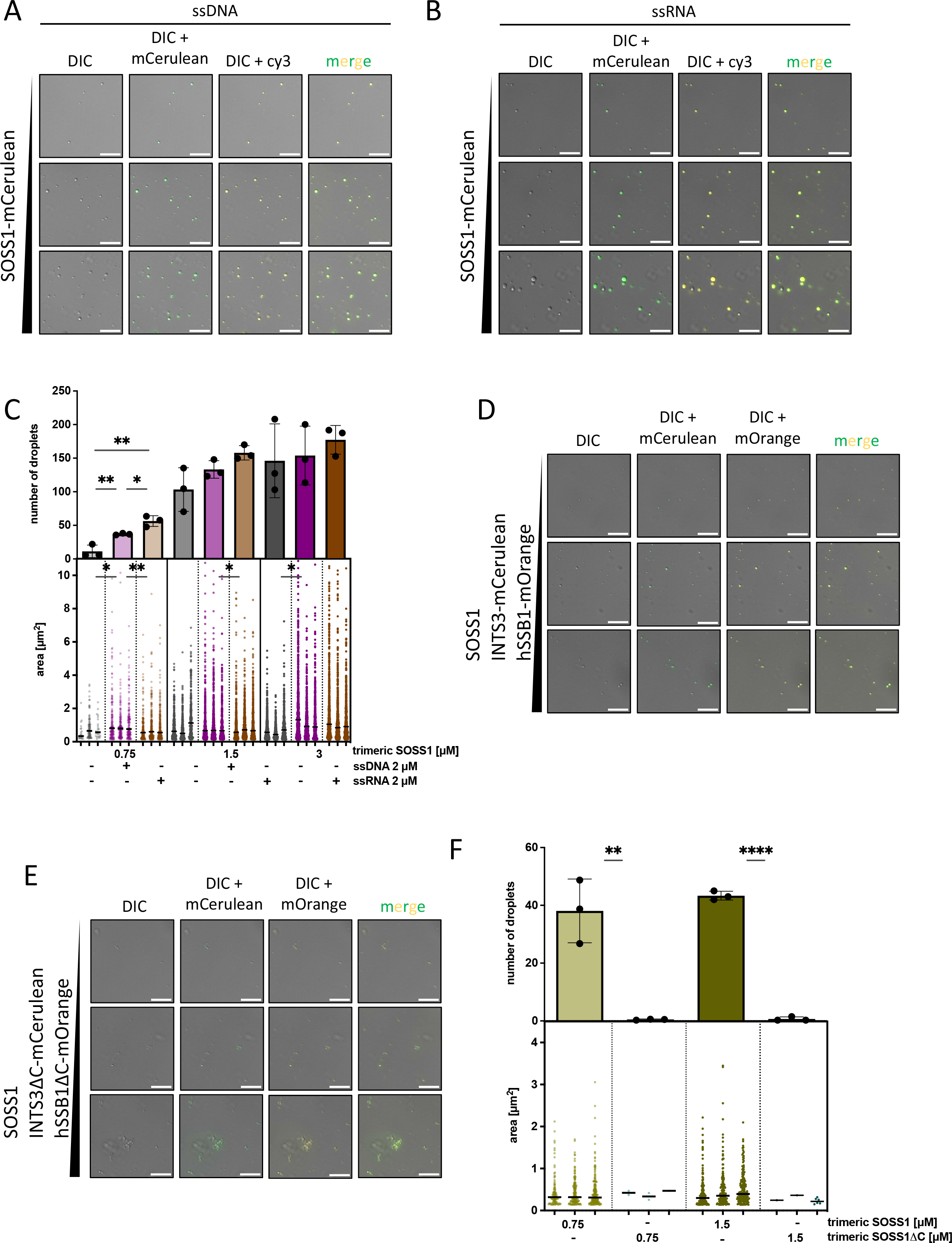
A) LLPS experiments determining concentration-dependent phase separation of fluorescently-labelled SOSS1 complex in the presence of 5% PEG-8000 and cy3-labelled ssDNA (2µM). Representative images from three experiments are depicted as differential interference contrast (DIC), overlay of DIC and mCerulean, overlay of DIC and cy3, and overlay of all three channels. Scale bar: 5µM. **B)** LLPS experiments determining concentration-dependent phase separation of fluorescently-labelled SOSS1 complex in the presence of 5% PEG-8000 and cy3-labelled ssRNA (2µM). Representative images from three experiments are depicted as differential interference contrast (DIC), overlay of DIC and mCerulean, overlay of DIC and cy3, and overlay of all three channels. Scale bar: 5µM. **C)** Bar chart (upper panel) represents quantification (n=3) of number of droplets from LLPS experiments shown in A), B), and experiments shown in Figure 6C. Statistical significance was determined by unpaired *t*-test. Nested scatter plot (lower panel) represents quantification (n=3) of area of individual droplets from three independent experiments, with median area determined per dataset. Statistical significance was determined by nested *t*-test. **D)** LLPS experiments determining phase separation of the trimeric SOSS1 complex labelled on INTS3 (INTS3-mCerulean) and hSSB1 (hSSB1-mOrange). Representative images from three experiments are depicted as differential interference contrast (DIC) and overlay of DIC and mCerulean, DIC and mOrange, and overlay of the three channels. Scale bar: 5µM. **E)** LLPS experiments determining phase separation of double-labelled trimeric SOSS1 complex containing INTS3^1-961^ (INTS3ΔC-mCerulean) and hSSB1^1-106^ (hSSB1ΔC-mOrange. Representative images from three experiments are depicted as differential interference contrast (DIC) and overlay of DIC and mCerulean. Scale bar: 5µM. **F)** Bar chart (upper panel) represents quantification (n=3) of number of droplets from LLPS experiments shown in D), E). Statistical significance was determined by unpaired *t*-test. Nested scatter plot (lower panel) represents quantification (n=3) of area of individual droplets from three independent experiments shown in D), E), with median area determined per dataset. Statistical significance was determined by nested *t*-test.

**Supplementary Figure 16.**
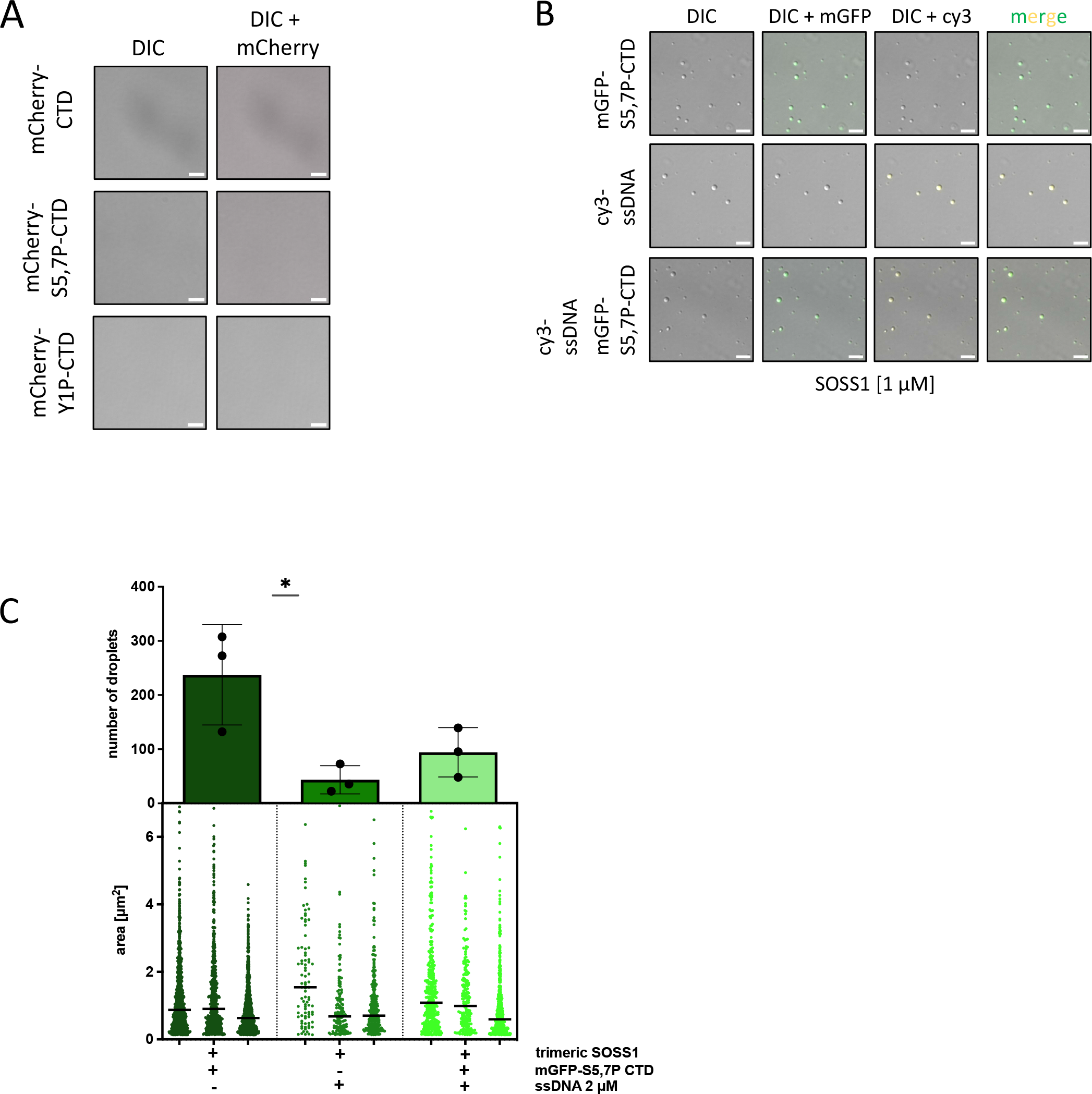
A) Control LLPS experiments determining phase separation of mCherry-CTDs (0.75 µM) in the presence of 5% PEG-8000. Representative images from three experiments are depicted as differential interference contrast (DIC) and overlay of DIC and mCherry. Scale bar: 5µM. **B)** LLPS experiments investigating the effect of mGFP-S5,7P-CTD (0,75 µM) and cy3- labelled ssDNA (2µM) on phase separation of unlabelled trimeric SOSS1 complex (1 µM). Representative images from three experiments are depicted as DIC, overlay of DIC and mGFP, overlay of DIC and cy3, and overlay of all three channels. Scale bar: 5µM. **C)** Bar chart (upper panel) represents quantification (n=3) of number of droplets from LLPS experiments shown in B). Statistical significance was determined by unpaired *t*-test. Nested scatter plot (lower panel) represents quantification (n=3) of area of individual droplets from three independent experiments shown in B), with median area determined per dataset. Statistical significance was determined by nested *t*-test.

**Supplementary Figure 17.**
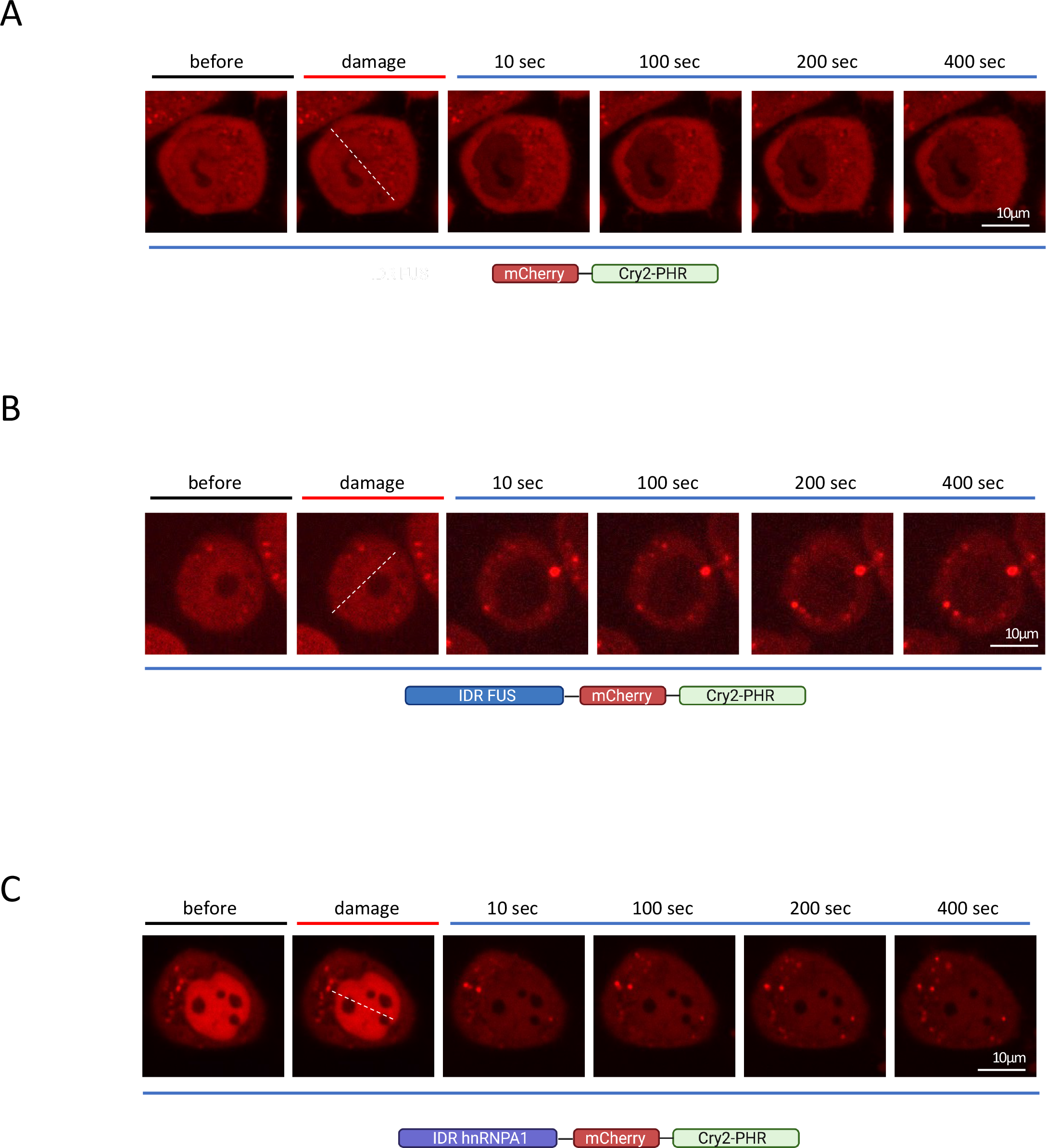
A-C) Damage induced optoDroplet formation in Cry2-mCherry (A), IDR-FUS-Cry2-mCherry (B) and IDR-hnRNPA1-Cry2-mCherry (C) cells. Representative images of optoDroplets before and after laser stripe and during light induction with indicated time points. Position of the laser stripe is marked with dashed white line. White rectangles show area of the image that is enlarged below.

**Supplementary Figure 18.**
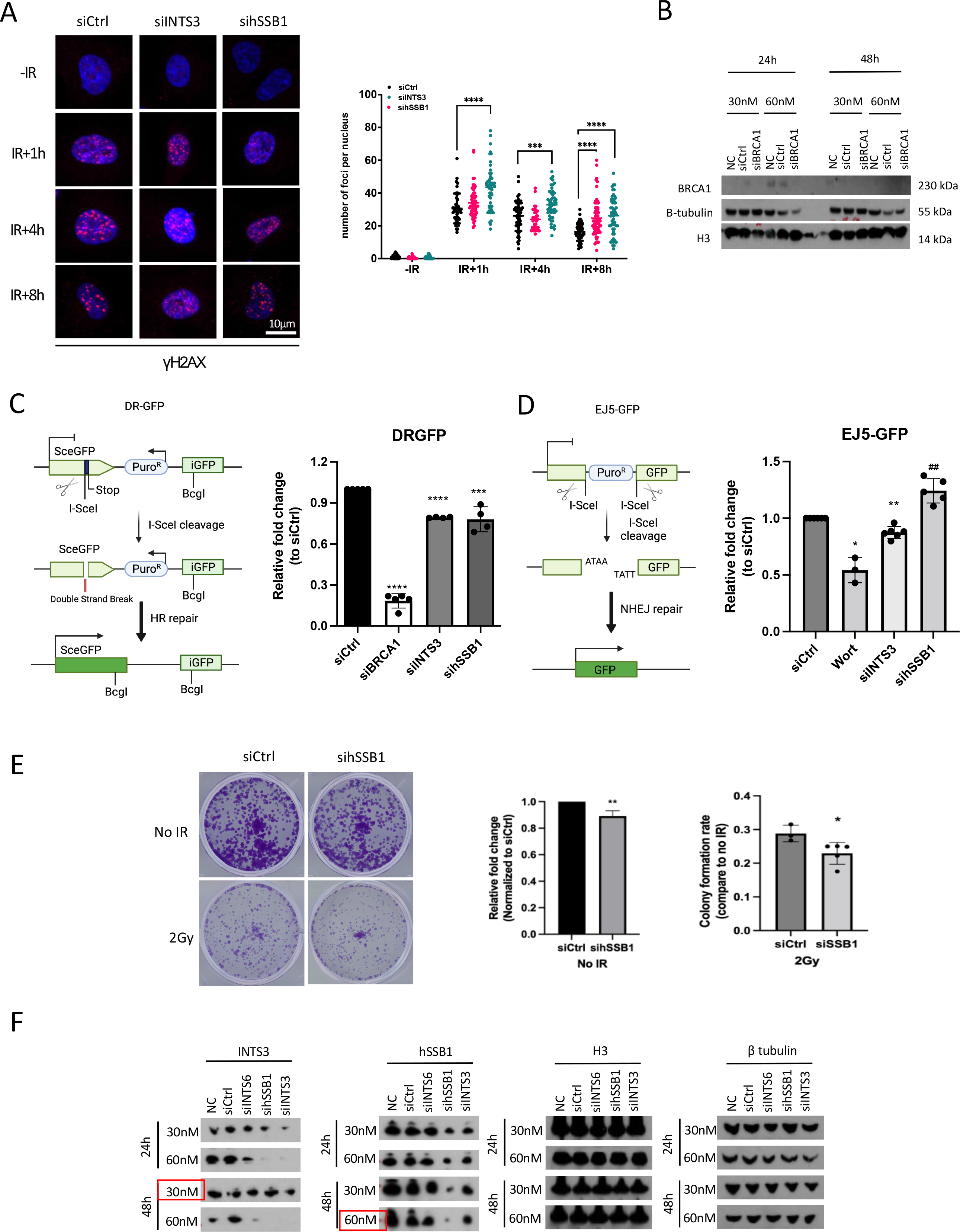
A) Representative IF images showing the levels of γH2AX in siControl, siINTS3 or sihSSB1 cells. Bar chart showing quantification of IF images, \*\*\*\**p* ≤ 0.0001. **B)** Representative images of western blot showing the protein level of BRCA1 in wt (NC), siCtrl or siBRCA1 cells at 24h or 48h after indicated concentrations of siRNA treatment. β- tubulin and H3 were used as loading controls. **C)** Drawing of DR-GFP HR reporter strategy. Bar chart shows the efficiency of HR repair in DR-GFP HeLa reporter cells, as measured by FACS. BRCA1 knockdown was used as the positive control. \*\*\**p* ≤ 0.001, \*\*\*\**p* ≤ 0.0001. **D)** Drawing of EJ5-GFP NHEJ reporter strategy. Bar chart showing the efficiency of NHEJ repair in EJ5 HeLa reporter cells, as measured by FACS. Wortmannin (DNA-PK inhibitor) was used as the positive control. \*\**p* ≤ 0.01, \**p* ≤ 0.05, compare with siCtrl significantly decreased; ^##^ *p* ≤ 0.01, compare with siCtrl significantly increased. **E)** Left: representative of clonogenic assay of siCtrl and sihSSB1 cells. The cells were stained and counted after 10 days of growing. Right: quantification of left (\*\**p* ≤ 0.01, \**p* ≤ 0.05). **F)** Representative images of western blot showing the protein levels of INTS3, hSSB1, H3 and β-tubulin in DR-GFP HeLa reporter cells after siINTS3 or siINTS6 or sihSSB1 treatment (24h or 48h). β-tubulin and H3 were used as loading controls. Chosen time point and siRNA concentration for each protein in HR reporter assay is indicated in red box.

**Supplementary table 1:**
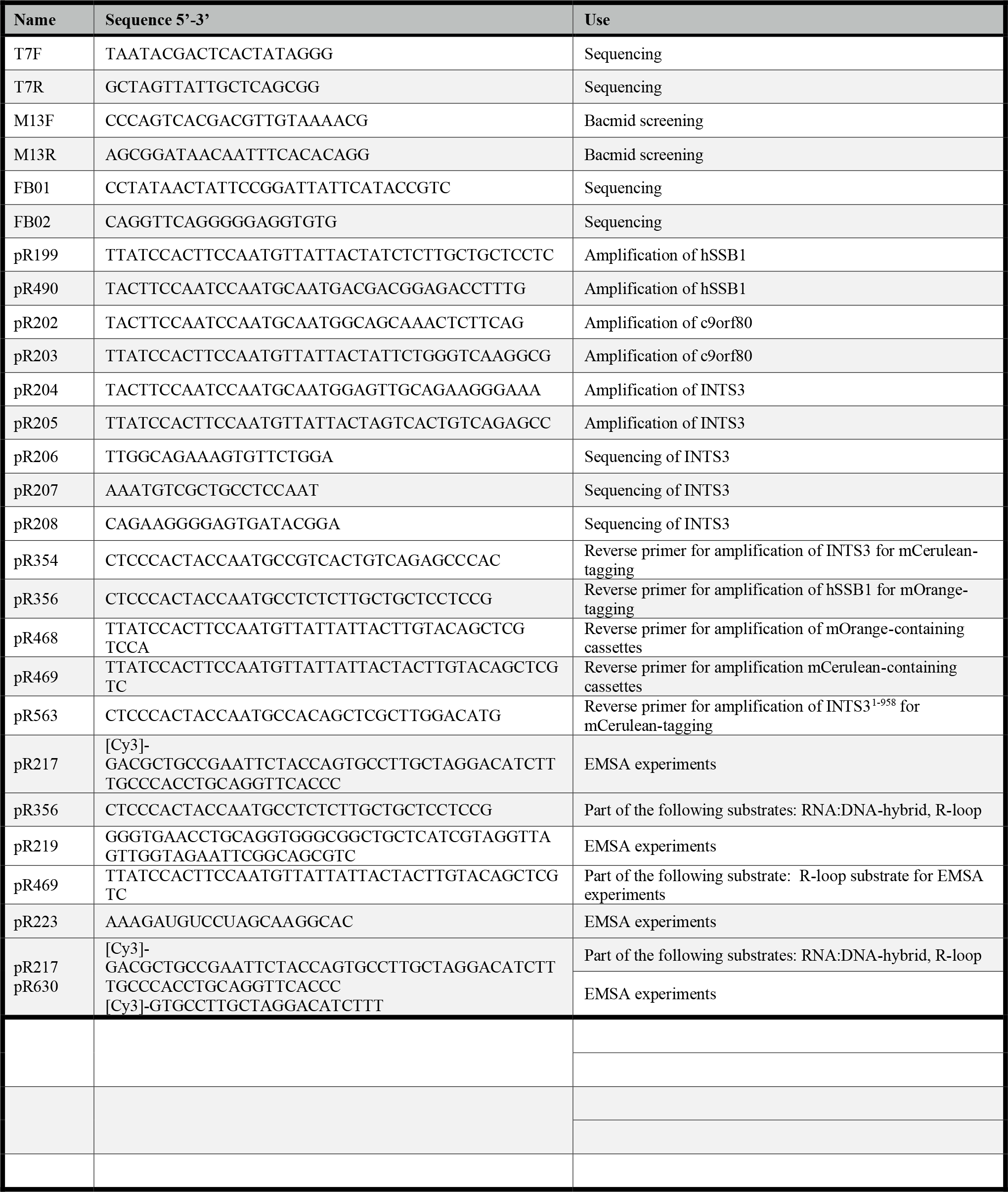
List of oligonucleotides used in *in vitro* work

**Supplementary table 2:**
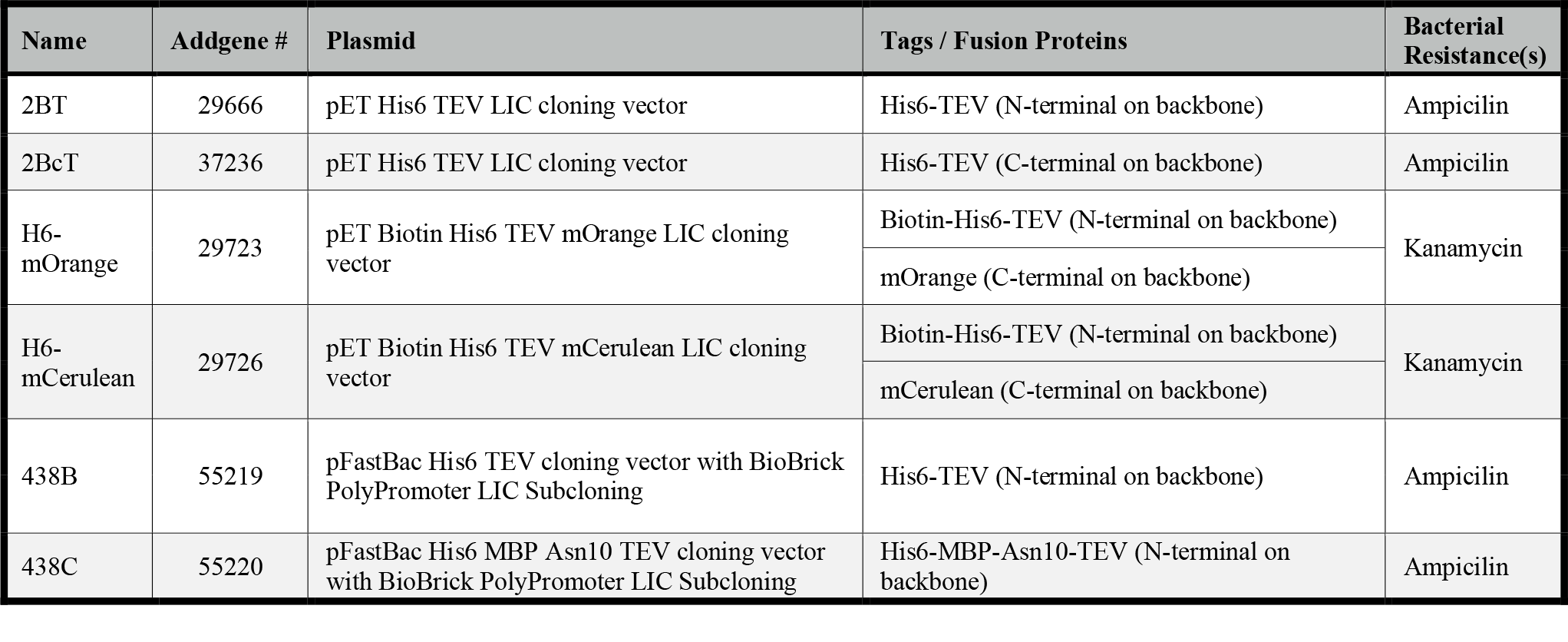
List of plasmids used in *in vitro* work

**Supplementary table 3:**
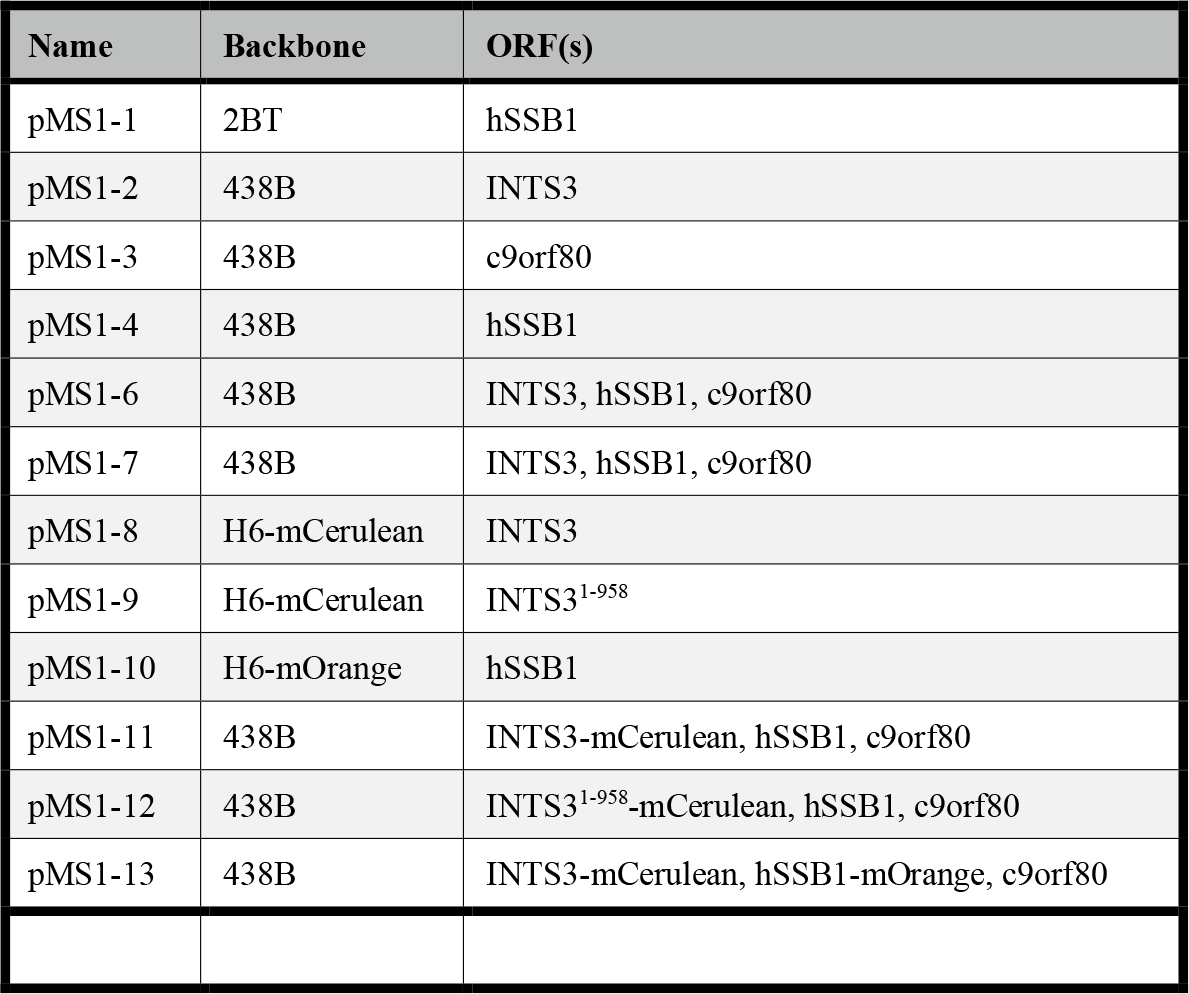
List of recombinant plasmids used in *in vitro* work

**Supplementary table 4:**
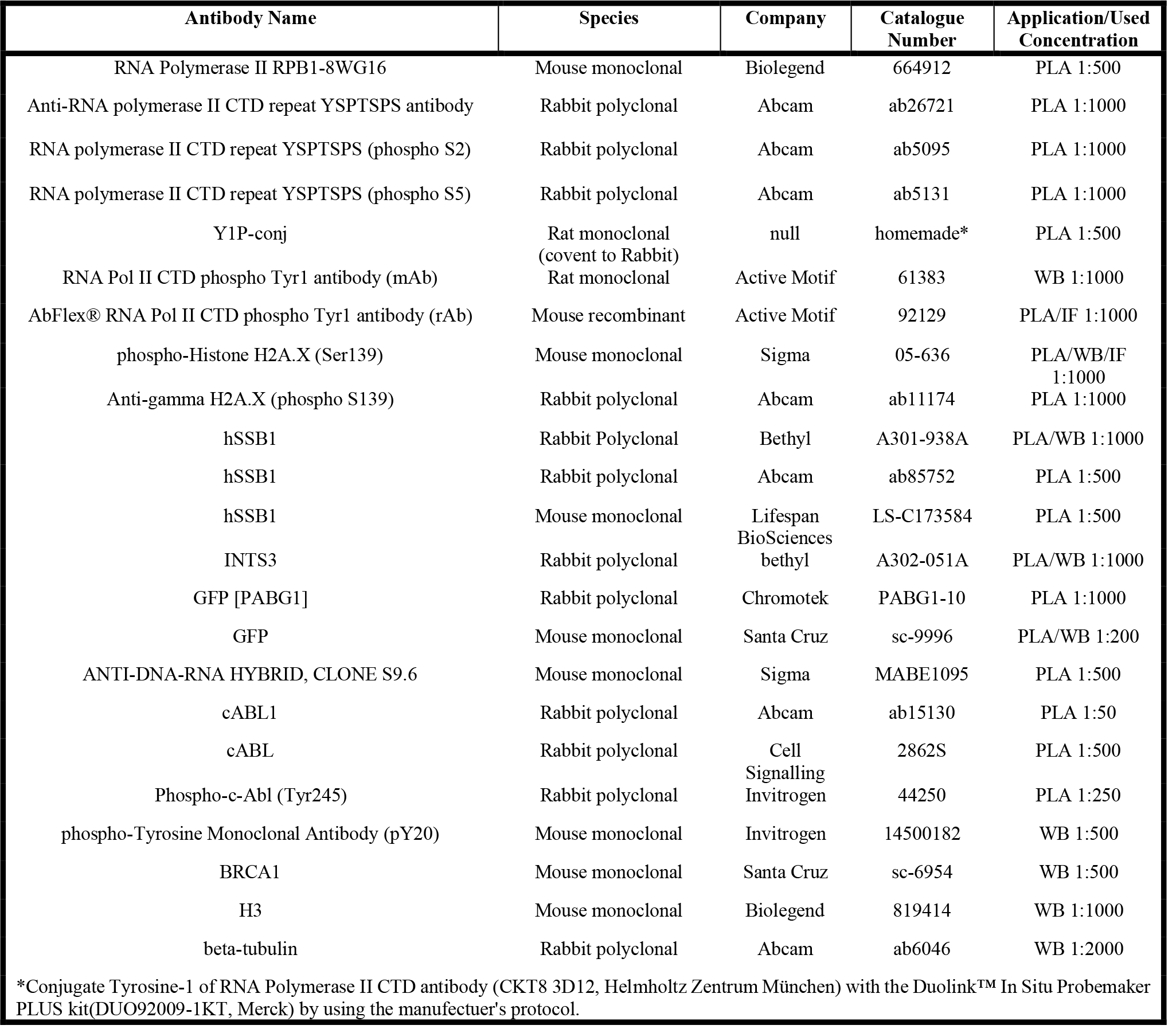
List of antibodies used in *in vivo* work

**Supplementary table 5:**
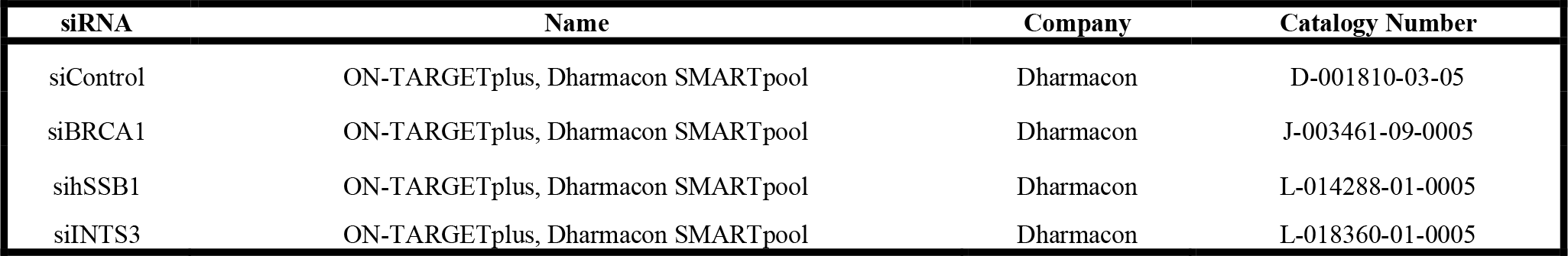
List of siRNA used in *in vivo* work

**Supplementary table 6:**
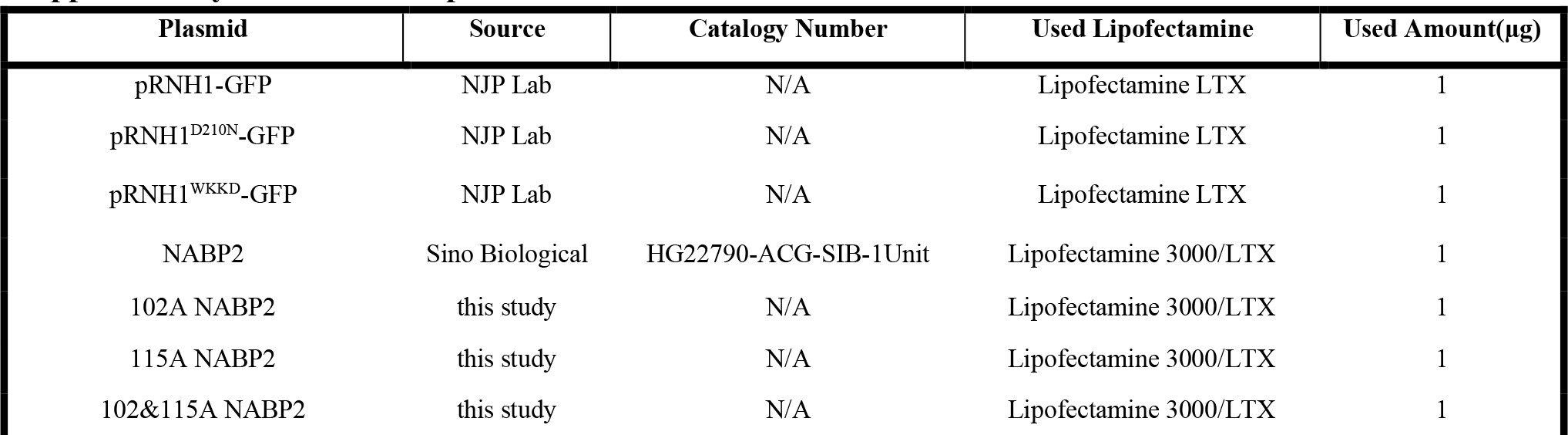
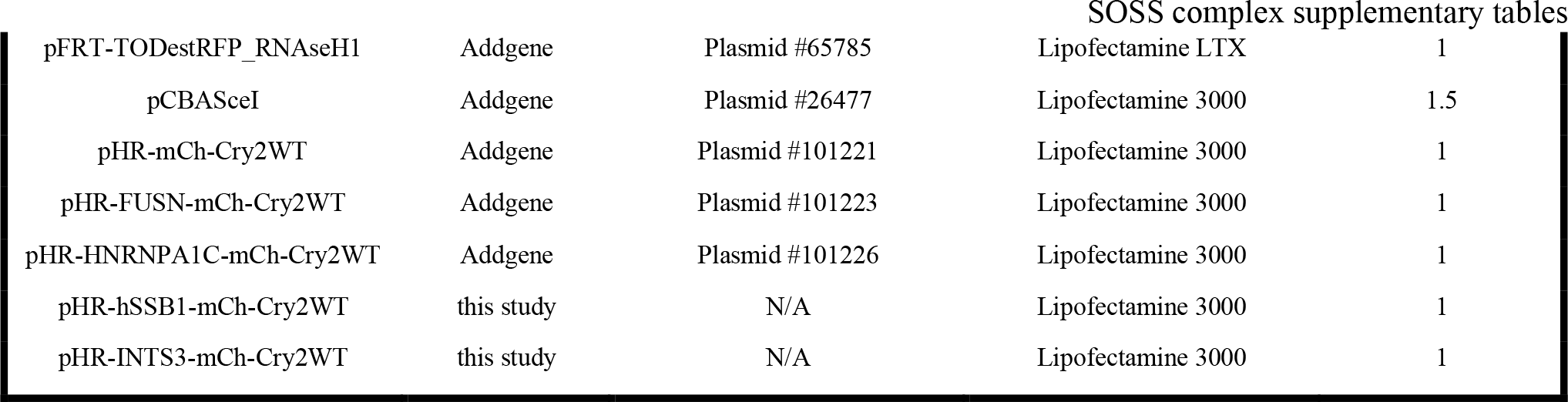
List of plasmids used in *in vivo* work

**Supplementary table 7:**
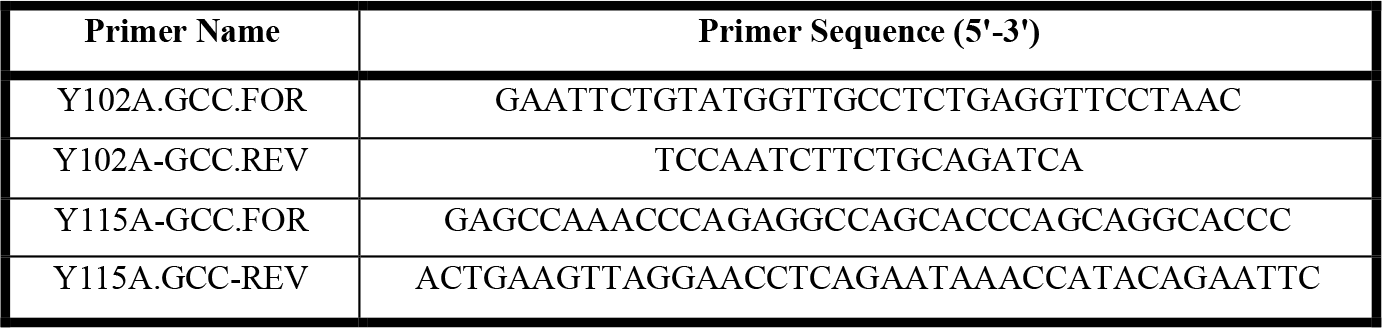
List of primers used for mutagenesis in this work

**Supplementary table 8:**
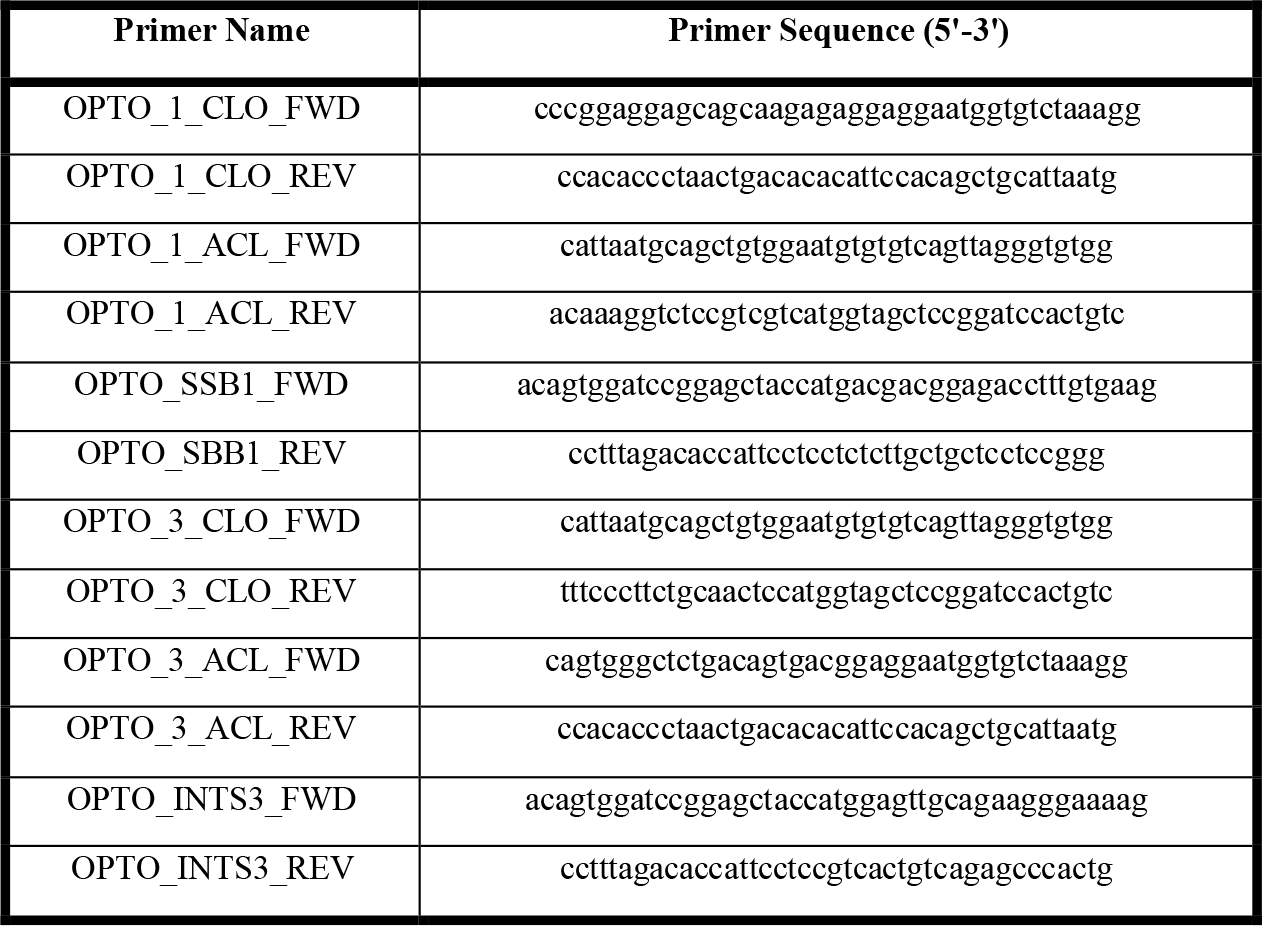
List of primers used for Gibson cloning in this work

**Supplementary table 9:**
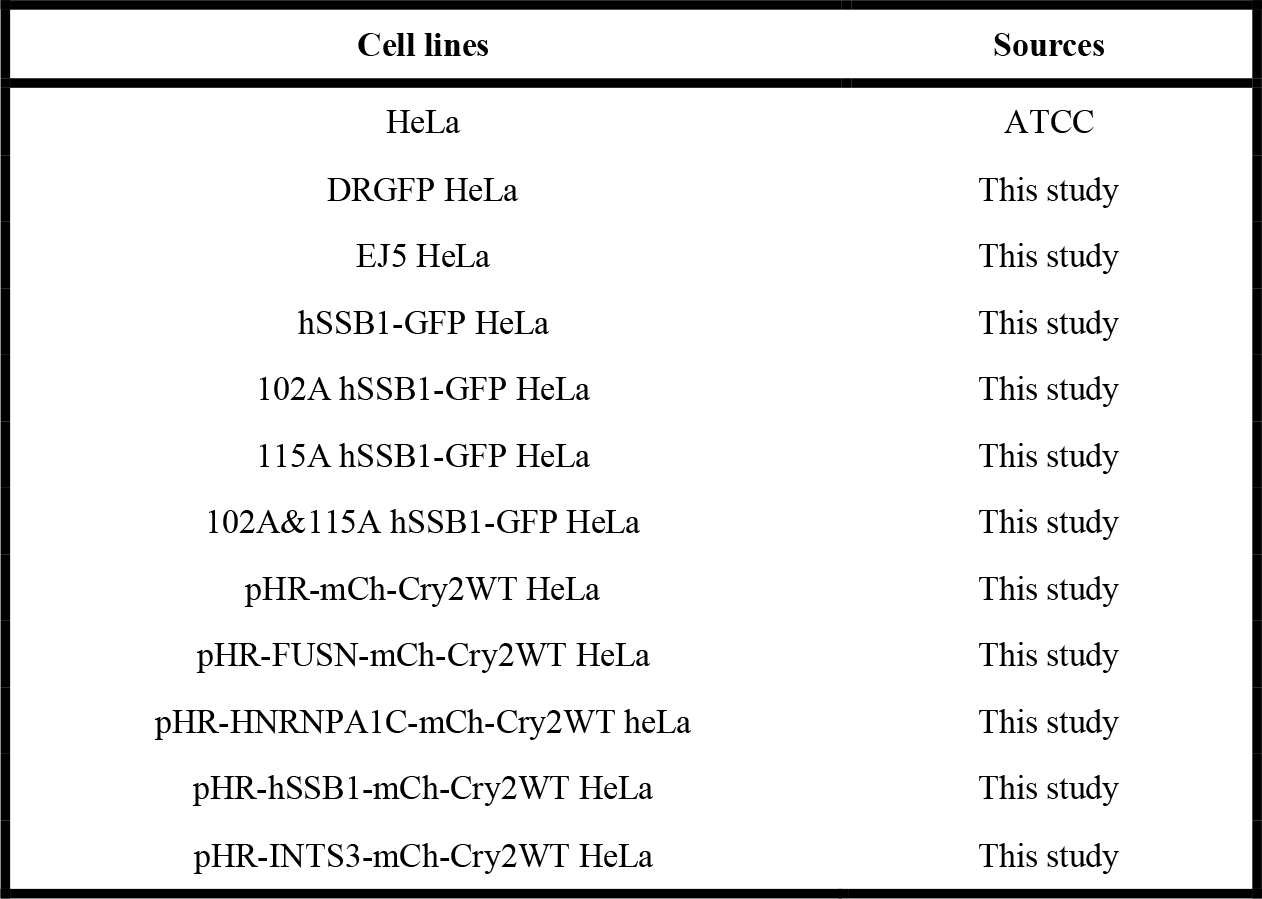
List of cell lines used in this work.

## REFERENCES

1. Q. Long, Z. Liu, M. Gullerova, Sweet Melody or Jazz? Transcription Around DNA Double-Strand Breaks. Front Mol Biosci 8, 655786 (2021).

2. H. E. Krokan, M. Bjoras, Base excision repair. Cold Spring Harb Perspect Biol 5, a012583 (2013).

3. O. D. Scharer, Nucleotide excision repair in eukaryotes. Cold Spring Harb Perspect Biol 5, a012609 (2013).

4. K. W. Caldecott, Single-strand break repair and genetic disease. Nat Rev Genet 9, 619–631 (2008).

5. M. R. Lieber, The mechanism of double-strand DNA break repair by the nonhomologous DNA end-joining pathway. Annu Rev Biochem 79, 181–211 (2010).

6. O. V. Iarovaia et al., Dynamics of double strand breaks and chromosomal translocations. Mol Cancer 13, 249 (2014).

7. N. W. Ashton, E. Bolderson, L. Cubeddu, K. J. O’Byrne, D. J. Richard, Human single- stranded DNA binding proteins are essential for maintaining genomic stability. BMC Mol Biol 14, 9 (2013).

8. L. V. Croft et al., Human single-stranded DNA binding protein 1 (hSSB1, OBFC2B), a critical component of the DNA damage response. Semin Cell Dev Biol 86, 121–128 (2019).

9. L. A. Yates et al., A structural and dynamic model for the assembly of Replication Protein A on single-stranded DNA. Nat Commun 9, 5447 (2018).

10. D. J. Richard et al., Single-stranded DNA-binding protein hSSB1 is critical for genomic stability. Nature 453, 677–681 (2008).

11. J. Huang, Z. Gong, G. Ghosal, J. Chen, SOSS complexes participate in the maintenance of genomic stability. Mol Cell 35, 384–393 (2009).

12. Y. Li et al., HSSB1 and hSSB2 form similar multiprotein complexes that participate in DNA damage response. J Biol Chem 284, 23525–23531 (2009).

13. E. Bolderson et al., Human single-stranded DNA binding protein 1 (hSSB1/NABP2) is required for the stability and repair of stalled replication forks. Nucleic Acids Res 42, 6326–6336 (2014).

14. N. Paquet et al., hSSB1 (NABP2/OBFC2B) is regulated by oxidative stress. Sci Rep 6, 27446 (2016).

15. A. Marechal, L. Zou, DNA damage sensing by the ATM and ATR kinases. Cold Spring Harb Perspect Biol 5, (2013).

16. I. S. Mohiuddin, M. H. Kang, DNA-PK as an Emerging Therapeutic Target in Cancer. Front Oncol 9, 635 (2019).

17. D. Menolfi, S. Zha, ATM, ATR and DNA-PKcs kinases-the lessons from the mouse models: inhibition not equal deletion. Cell Biosci 10, 8 (2020).

18. S. Matsuoka et al., ATM and ATR substrate analysis reveals extensive protein networks responsive to DNA damage. Science 316, 1160–1166 (2007).

19. M. B. Smolka, C. P. Albuquerque, S. H. Chen, H. Zhou, Proteome-wide identification of in vivo targets of DNA damage checkpoint kinases. Proc Natl Acad Sci U S A 104, 10364–10369 (2007).

20. M. T. M. van Jaarsveld et al., Cell-type-specific role of CHK2 in mediating DNA damage-induced G2 cell cycle arrest. Oncogenesis 9, 35 (2020).

21. E. M. Bahassi et al., The checkpoint kinases Chk1 and Chk2 regulate the functional associations between hBRCA2 and Rad51 in response to DNA damage. Oncogene 27, 3977–3985 (2008).

22. J. Drouet et al., Interplay between Ku, Artemis, and the DNA-dependent protein kinase catalytic subunit at DNA ends. J Biol Chem 281, 27784–27793 (2006).

23. V. Meltser, M. Ben-Yehoyada, Y. Shaul, c-Abl tyrosine kinase in the DNA damage response: cell death and more. Cell Death Differ 18, 2–4 (2011).

24. K. Burger, M. Schlackow, M. Gullerova, Tyrosine kinase c-Abl couples RNA polymerase II transcription to DNA double-strand breaks. Nucleic Acids Res 47, 3467–3484 (2019).

25. F. Michelini et al., Damage-induced lncRNAs control the DNA damage response through interaction with DDRNAs at individual double-strand breaks. Nat Cell Biol 19, 1400–1411 (2017).

26. S. Alberti, A. Gladfelter, T. Mittag, Considerations and Challenges in Studying Liquid- Liquid Phase Separation and Biomolecular Condensates. Cell 176, 419–434 (2019).

27. F. Frottin et al., The nucleolus functions as a phase-separated protein quality control compartment. Science 365, 342–347 (2019).

28. M. Boehning et al., RNA polymerase II clustering through carboxy-terminal domain phase separation. Nat Struct Mol Biol 25, 833–840 (2018).

29. B. R. Sabari et al., Coactivator condensation at super-enhancers links phase separation and gene control. Science 361, (2018).

30. Y. Shin, C. P. Brangwynne, Liquid phase condensation in cell physiology and disease. Science 357, (2017).

31. F. Pessina et al., Functional transcription promoters at DNA double-strand breaks mediate RNA-driven phase separation of damage-response factors. Nat Cell Biol 21, 1286–1299 (2019).

32. S. Kilic et al., Phase separation of 53BP1 determines liquid-like behavior of DNA repair compartments. EMBO J 38, e101379 (2019).

33. X. J. Fan et al., NONO phase separation enhances DNA damage repair by accelerating nuclear EGFR-induced DNA-PK activation. Am J Cancer Res 11, 2838–2852 (2021).

34. N. Descostes et al., Tyrosine phosphorylation of RNA polymerase II CTD is associated with antisense promoter transcription and active enhancers in mammalian cells. Elife 3, e02105 (2014).

35. N. Shah et al., Tyrosine-1 of RNA Polymerase II CTD Controls Global Termination of Gene Transcription in Mammals. Mol Cell 69, 48–61 e46 (2018).

36. A. Alagia, R. F. Ketley, M. Gullerova, Proximity Ligation Assay for Detection of R- Loop Complexes upon DNA Damage. Methods Mol Biol 2528, 289–303 (2022).

37. D. F. Allison, G. G. Wang, R-loops: formation, function, and relevance to cell stress. Cell Stress 3, 38–46 (2019).

38. R. Kato, K. Miyagawa, T. Yasuhara, The role of R-loops in transcription-associated DNA double-strand break repair. Mol Cell Oncol 6, 1542244 (2019).

39. C. Touma et al., A structural analysis of DNA binding by hSSB1 (NABP2/OBFC2B) in solution. Nucleic Acids Res 44, 7963–7973 (2016).

40. W. Ren et al., Structural basis of SOSS1 complex assembly and recognition of ssDNA. Cell Rep 6, 982–991 (2014).

41. E. W. Martin, A. S. Holehouse, Intrinsically disordered protein regions and phase separation: sequence determinants of assembly or lack thereof. Emerg Top Life Sci 4, 307–329 (2020).

42. W. Borcherds, A. Bremer, M. B. Borgia, T. Mittag, How do intrinsically disordered protein regions encode a driving force for liquid-liquid phase separation? Curr Opin Struct Biol 67, 41–50 (2021).

43. H. Lu et al., Phase-separation mechanism for C-terminal hyperphosphorylation of RNA polymerase II. Nature 558, 318–323 (2018).

44. W. K. Cho et al., Mediator and RNA polymerase II clusters associate in transcription- dependent condensates. Science 361, 412–415 (2018).

45. L. M. Appel et al., PHF3 regulates neuronal gene expression through the Pol II CTD reader domain SPOC. Nat Commun 12, 6078 (2021).

46. Y. Shin et al., Spatiotemporal Control of Intracellular Phase Transitions Using Light- Activated optoDroplets. Cell 168, 159–171 e114 (2017).

47. N. W. Ashton et al., hSSB1 phosphorylation is dynamically regulated by DNA-PK and PPP-family protein phosphatases. DNA Repair (Amst) 54, 30–39 (2017).

48. M. Sebesta et al., Role of PCNA and TLS polymerases in D-loop extension during homologous recombination in humans. DNA Repair (Amst) 12, 691–698 (2013).

49. K. Stejskal, D. Potesil, Z. Zdrahal, Suppression of peptide sample losses in autosampler vials. J Proteome Res 12, 3057–3062 (2013).

50. M. R. Lamprecht, D. M. Sabatini, A. E. Carpenter, CellProfiler: free, versatile software for automated biological image analysis. Biotechniques 42, 71–75 (2007).

