## Supplementary Tables for "Phosphorylated trimeric SOSS1 complex and RNA polymerase II trigger liquid-liquid phase separation at double-strand breaks"

**Supplementary table 1: List of oligonucleotides used in *in vitro* work**

| Name | Sequence 5'-3' | Use |
|---|---|---|
| T7F | TAATACGACTCACTATAGGG | Sequencing |
| T7R | GCTAGTTATTGCTCAGCGG | Sequencing |
| M13F | CCCAGTCACGACGTTGTAAAACG | Bacmid screening |
| M13R | AGCGGATAACAATTTCACACAGG | Bacmid screening |
| FB01 | CCTATAACTATTCCGGATTATTCATACCGTC | Sequencing |
| FB02 | CAGGTTCAGGGGGAGGTGTG | Sequencing |
| pR199 | TTATCCACTTCCAATGTTATTACTATCTCTTGCTGCTCCTC | Amplification of hSSB1 |
| pR490 | TACTTCCAATCCAATGCAATGACGACGGAGACCTTTG | Amplification of hSSB1 |
| pR202 | TACTTCCAATCCAATGCAATGGCAGCAAACTCTTCAG | Amplification of c9orf80 |
| pR203 | TTATCCACTTCCAATGTTATTACTATTCTGGGTCAAGGCG | Amplification of c9orf80 |
| pR204 | TACTTCCAATCCAATGCAATGGAGTTGCAGAAGGGAAA | Amplification of INTS3 |
| pR205 | TTATCCACTTCCAATGTTATTACTAGTCACTGTCAGAGCC | Amplification of INTS3 |
| pR206 | TTGGCAGAAAGTGTTCTGGA | Sequencing of INTS3 |
| pR207 | AAATGTCGCTGCCTCCAAT | Sequencing of INTS3 |
| pR208 | CAGAAGGGGAGTGATACGGA | Sequencing of INTS3 |
| pR354 | CTCCCACTACCAATGCCGTCACTGTCAGAGCCCAC | Reverse primer for amplification of INTS3 for mCerulean-tagging |
| pR356 | CTCCCACTACCAATGCCTCTCTTGCTGCTCCTCCG | Reverse primer for amplification of hSSB1 for mOrange-tagging |
| pR468 | TTATCCACTTCCAATGTTATTATTACTTGTACAGCTCG TCCA | Reverse primer for amplification of mOrange-containing cassettes |
| pR469 | TTATCCACTTCCAATGTTATTATTACTACTTGTACAGCTCG TC | Reverse primer for amplification mCerulean-containing cassettes |
| pR563 | CTCCCACTACCAATGCCACAGCTCGCTTGGACATG | Reverse primer for amplification of $INTS3^{1-958}$ for mCerulean-tagging |
| pR217 | [Cy3]- GACGCTGCCGAATTCTACCAGTGCCTTGCTAGGACATCTT TGCCCACCTGCAGGTTCACCC | EMSA experiments |
| pR356 | CTCCCACTACCAATGCCTCTCTTGCTGCTCCTCCG | Part of the following substrates: RNA:DNA-hybrid, R-loop |
| pR219 | GGGTGAACCTGCAGGTGGGCGGCTGCTCATCGTAGGTTA GTTGGTAGAATTCGGCAGCGTC | EMSA experiments |
| pR469 | TTATCCACTTCCAATGTTATTATTACTACTTGTACAGCTCG TC | Part of the following substrate: R-loop substrate for EMSA experiments |
| pR223 | AAAGAUGUCCUAGCAAGGCAC | EMSA experiments |
| pR217 | [Cy3]- GACGCTGCCGAATTCTACCAGTGCCTTGCTAGGACATCTT TGCCCACCTGCAGGTTCACCC | Part of the following substrates: RNA:DNA-hybrid, R-loop |
| pR630 | [Cy3]-GTGCCTTGCTAGGACATCTTT | EMSA experiments |

**Supplementary table 2: List of plasmids used in *in vitro* work**

| Name | Addgene # | Plasmid | Tags / Fusion Proteins | Bacterial Resistance(s) |
|---|---|---|---|---|
| 2BT | 29666 | pET His6 TEV LIC cloning vector | His6-TEV (N-terminal on backbone) | Ampicilin |
| 2BcT | 37236 | pET His6 TEV LIC cloning vector | His6-TEV (C-terminal on backbone) | Ampicilin |
| H6-mOrange | 29723 | pET Biotin His6 TEV mOrange LIC cloning vector | Biotin-His6-TEV (N-terminal on backbone)<br>mOrange (C-terminal on backbone) | Kanamycin |
| H6-mCerulean | 29726 | pET Biotin His6 TEV mCerulean LIC cloning vector | Biotin-His6-TEV (N-terminal on backbone)<br>mCerulean (C-terminal on backbone) | Kanamycin |
| 438B | 55219 | pFastBac His6 TEV cloning vector with BioBrick PolyPromoter LIC Subcloning | His6-TEV (N-terminal on backbone) | Ampicilin |
| 438C | 55220 | pFastBac His6 MBP Asn10 TEV cloning vector with BioBrick PolyPromoter LIC Subcloning | His6-MBP-Asn10-TEV (N-terminal on backbone) | Ampicilin |

**Supplementary table 3: List of recombinant plasmids used in *in vitro* work**

| Name | Backbone | ORF(s) |
|---|---|---|
| pMS1-1 | 2BT | hSSB1 |
| pMS1-2 | 438B | INTS3 |
| pMS1-3 | 438B | c9orf80 |
| pMS1-4 | 438B | hSSB1 |
| pMS1-6 | 438B | INTS3, hSSB1, c9orf80 |
| pMS1-7 | 438B | INTS3, hSSB1, c9orf80 |
| pMS1-8 | H6-mCerulean | INTS3 |
| pMS1-9 | H6-mCerulean | INTS3$^{1-958}$ |
| pMS1-10 | H6-mOrange | hSSB1 |
| pMS1-11 | 438B | INTS3-mCerulean, hSSB1, c9orf80 |
| pMS1-12 | 438B | INTS3$^{1-958}$-mCerulean, hSSB1, c9orf80 |
| pMS1-13 | 438B | INTS3-mCerulean, hSSB1-mOrange, c9orf80 |

**Supplementary table 4: List of antibodies used in *in vivo* work**

| Antibody Name | Species | Company | Catalogue Number | Application/Used Concentration |
|---|---|---|---|---|
| RNA Polymerase II RPB1-8WG16 | Mouse monoclonal | Biolegend | 664912 | PLA 1:500 |
| Anti-RNA polymerase II CTD repeat YSPTSPS antibody | Rabbit polyclonal | Abcam | ab26721 | PLA 1:1000 |
| RNA polymerase II CTD repeat YSPTSPS (phospho S2) | Rabbit polyclonal | Abcam | ab5095 | PLA 1:1000 |
| RNA polymerase II CTD repeat YSPTSPS (phospho S5) | Rabbit polyclonal | Abcam | ab5131 | PLA 1:1000 |
| Y1P-conj | Rat monoclonal (covent to Rabbit) | null | homemade* | PLA 1:500 |
| RNA Pol II CTD phospho Tyr1 antibody (mAb) | Rat monoclonal | Active Motif | 61383 | WB 1:1000 |
| AbFlex® RNA Pol II CTD phospho Tyr1 antibody (rAb) | Mouse recombinant | Active Motif | 92129 | PLA/IF 1:1000 |
| phospho-Histone H2A.X (Ser139) | Mouse monoclonal | Sigma | 05-636 | PLA/WB/IF 1:1000 |
| Anti-gamma H2A.X (phospho S139) | Rabbit polyclonal | Abcam | ab11174 | PLA 1:1000 |
| hSSB1 | Rabbit Polyclonal | Bethyl | A301-938A | PLA/WB 1:1000 |
| hSSB1 | Rabbit polyclonal | Abcam | ab85752 | PLA 1:500 |
| hSSB1 | Mouse monoclonal | Lifespan BioSciences | LS-C173584 | PLA 1:500 |
| INTS3 | Rabbit polyclonal | bethyl | A302-051A | PLA/WB 1:1000 |
| GFP [PABG1] | Rabbit polyclonal | Chromotek | PABG1-10 | PLA 1:1000 |
| GFP | Mouse monoclonal | Santa Cruz | sc-9996 | PLA/WB 1:200 |
| ANTI-DNA-RNA HYBRID, CLONE S9.6 | Mouse monoclonal | Sigma | MABE1095 | PLA 1:500 |
| cABL1 | Rabbit polyclonal | Abcam | ab15130 | PLA 1:50 |
| cABL | Rabbit polyclonal | Cell Signalling | 2862S | PLA 1:500 |
| Phospho-c-Abl (Tyr245) | Rabbit polyclonal | Invitrogen | 44250 | PLA 1:250 |
| phospho-Tyrosine Monoclonal Antibody (pY20) | Mouse monoclonal | Invitrogen | 14500182 | WB 1:500 |
| BRCA1 | Mouse monoclonal | Santa Cruz | sc-6954 | WB 1:500 |
| H3 | Mouse monoclonal | Biolegend | 819414 | WB 1:1000 |
| beta-tubulin | Rabbit polyclonal | Abcam | ab6046 | WB 1:2000 |

*Conjugate Tyrosine-1 of RNA Polymerase II CTD antibody (CKT8 3D12, Helmholtz Zentrum München) with the Duolink™ In Situ Probemaker PLUS kit(DUO92009-1KT, Merck) by using the manufectuer's protocol.

**Supplementary table 5: List of siRNA used in *in vivo* work**

| siRNA | Name | Company | Catalogy Number |
|---|---|---|---|
| siControl | ON-TARGETplus, Dharmacon SMARTpool | Dharmacon | D-001810-03-05 |
| siBRCA1 | ON-TARGETplus, Dharmacon SMARTpool | Dharmacon | J-003461-09-0005 |
| sihSSB1 | ON-TARGETplus, Dharmacon SMARTpool | Dharmacon | L-014288-01-0005 |
| siINTS3 | ON-TARGETplus, Dharmacon SMARTpool | Dharmacon | L-018360-01-0005 |

**Supplementary table 6: List of plasmids used in *in vivo* work**

| Plasmid | Source | Catalogy Number | Used Lipofectamine | Used Amount(μg) |
|---|---|---|---|---|
| pRNH1-GFP | NJP Lab | N/A | Lipofectamine LTX | 1 |
| pRNH1$^{D210N}$-GFP | NJP Lab | N/A | Lipofectamine LTX | 1 |
| pRNH1$^{WKKD}$-GFP | NJP Lab | N/A | Lipofectamine LTX | 1 |
| NABP2 | Sino Biological | HG22790-ACG-SIB-1Unit | Lipofectamine 3000/LTX | 1 |
| 102A NABP2 | this study | N/A | Lipofectamine 3000/LTX | 1 |
| 115A NABP2 | this study | N/A | Lipofectamine 3000/LTX | 1 |
| 102&115A NABP2 | this study | N/A | Lipofectamine 3000/LTX | 1 |

| | | | | |
|---|---|---|---|---|
| pFRT-TODestRFP_RNAseH1 | Addgene | Plasmid #65785 | Lipofectamine LTX | 1 |
| pCBASceI | Addgene | Plasmid #26477 | Lipofectamine 3000 | 1.5 |
| pHR-mCh-Cry2WT | Addgene | Plasmid #101221 | Lipofectamine 3000 | 1 |
| pHR-FUSN-mCh-Cry2WT | Addgene | Plasmid #101223 | Lipofectamine 3000 | 1 |
| pHR-HNRNPA1C-mCh-Cry2WT | Addgene | Plasmid #101226 | Lipofectamine 3000 | 1 |
| pHR-hSSB1-mCh-Cry2WT | this study | N/A | Lipofectamine 3000 | 1 |
| pHR-INTS3-mCh-Cry2WT | this study | N/A | Lipofectamine 3000 | 1 |

**Supplementary table 7: List of primers used for mutagenesis in this work**

| Primer Name | Primer Sequence (5'-3') |
|---|---|
| Y102A.GCC.FOR | GAATTCTGTATGGTTGCCTCTGAGGTTCCTAAC |
| Y102A-GCC.REV | TCCAATCTTCTGCAGATCA |
| Y115A-GCC.FOR | GAGCCAAACCCAGAGGCCAGCACCCAGCAGGCACCC |
| Y115A.GCC-REV | ACTGAAGTTAGGAACCTCAGAATAAACCATACAGAATTC |

**Supplementary table 8: List of primers used for Gibson cloning in this work**

| Primer Name | Primer Sequence (5'-3') |
|---|---|
| OPTO_1_CLO_FWD | cccggaggagcagcaagagaggaggaatggtgtctaaagg |
| OPTO_1_CLO_REV | ccacaccctaactgacacacattccacagctgcattaatg |
| OPTO_1_ACL_FWD | cattaatgcagctgtggaatgtgtgtcagttagggtgtgg |
| OPTO_1_ACL_REV | acaaaggtctccgtcgtcatggtagctccggatccactgtc |
| OPTO_SSB1_FWD | acagtggatccggagctaccatgacgacggagacctttgtgaag |
| OPTO_SBB1_REV | cctttagacaccattcctcctctcttgctgctcctccggg |
| OPTO_3_CLO_FWD | cattaatgcagctgtggaatgtgtgtcagttagggtgtgg |
| OPTO_3_CLO_REV | tttcccttctgcaactccatggtagctccggatccactgtc |
| OPTO_3_ACL_FWD | cagtgggctctgacagtgacggaggaatggtgtctaaagg |
| OPTO_3_ACL_REV | ccacaccctaactgacacacattccacagctgcattaatg |
| OPTO_INTS3_FWD | acagtggatccggagctaccatggagttgcagaagggaaaag |
| OPTO_INTS3_REV | cctttagacaccattcctccgtcactgtcagagcccactg |

**Supplementary table 9: List of cell lines used in this work.**

| Cell lines | Sources |
|---|---|
| HeLa | ATCC |
| DRGFP HeLa | This study |
| EJ5 HeLa | This study |
| hSSB1-GFP HeLa | This study |
| 102A hSSB1-GFP HeLa | This study |
| 115A hSSB1-GFP HeLa | This study |
| 102A&115A hSSB1-GFP HeLa | This study |
| pHR-mCh-Cry2WT HeLa | This study |
| pHR-FUSN-mCh-Cry2WT HeLa | This study |
| pHR-HNRNPA1C-mCh-Cry2WT heLa | This study |
| pHR-hSSB1-mCh-Cry2WT HeLa | This study |
| pHR-INTS3-mCh-Cry2WT HeLa | This study |
